# Treatment with commonly used antiretroviral drugs induces a type I/III interferon signature in the gut in the absence of HIV infection

**DOI:** 10.1101/701961

**Authors:** Sean M. Hughes, Claire N. Levy, Fernanda L. Calienes, Joanne D. Stekler, Urvashi Pandey, Lucia Vojtech, Alicia R. Berard, Kenzie Birse, Laura Noël-Romas, Brian Richardson, Jackelyn B. Golden, Michael Cartwright, Ann C. Collier, Claire E. Stevens, Marcel E. Curlin, Timothy H. Holtz, Nelly Mugo, Elizabeth Irungu, Elly Katabira, Timothy Muwonge, Javier R. Lama, Jared M. Baeten, Adam Burgener, Jairam R. Lingappa, M. Juliana McElrath, Romel Mackelprang, Ian McGowan, Ross D. Cranston, Mark J. Cameron, Florian Hladik

## Abstract

Tenofovir disoproxil fumarate (TDF) and emtricitabine (FTC) are used for HIV treatment and prevention. Previously, we found that topical rectal tenofovir gel caused immunological changes in the mucosa. Here we assessed the effect of oral TDF/FTC in three HIV pre-exposure prophylaxis trials, two with gastrointestinal and one with cervicovaginal biopsies. TDF/FTC induced type I/III interferon-related (IFN I/III) genes in the gastrointestinal tract, but not blood, with strong correlations between the two independent rectal biopsy groups (Spearman r=0.91) and between the rectum and duodenum (r=0.81). Gene set testing also indicated stimulation of type I/III pathways in the ectocervix, as well as of cellular proliferation in the duodenum. mRNA sequencing, digital droplet PCR, proteomics, and immunofluorescence staining confirmed IFN I/III pathway stimulation in the gastrointestinal tract. Thus, oral TDF/FTC stimulates an IFN-I/III signature throughout the gut, which could increase antiviral efficacy but also cause chronic immune activation in HIV prevention and treatment settings.

## Introduction

Despite highly active antiretroviral treatment (ART), persons living with HIV (PLWH) are more likely than HIV-uninfected individuals to experience non-AIDS related morbidities, such as non-AIDS defining cancers, cardiovascular disease, osteoporosis, frailty and other conditions associated with aging^1^. This increased morbidity appears to be associated, at least in part, with low-level immune activation persisting even in the face of complete suppression of plasma viremia^2^. The extent to which ART itself may contribute to this chronic immune activation has been a nagging concern without adequate investigation.

The two nucleoside/nucleotide reverse transcriptase inhibitors (NRTIs) tenofovir and emtricitabine are common components of ART, which must be taken life-long to prevent clinical progression to AIDS. Tenofovir and emtricitabine in a single combination pill also compose the only licensed oral pre-exposure prophylaxis (PrEP) intervention for uninfected persons at risk of HIV infection^3,4^. While tenofovir and emtricitabine are generally well tolerated and considered safe, their effects specifically on the immune system have been little studied. In particular, no studies have evaluated their contribution to immune dysregulation in the gastrointestinal (GI) tract. This is surprising, given that the GI tract harbors the largest HIV burden on and off ART^5^ and is a major source of immune dysfunction in PLWH^2^. Moreover, two trials of tenofovir application directly to the female genital tract or the rectum as topical PrEP demonstrated a pro-inflammatory effect on the mucosa^6,7^. Analogous to these studies of topical PrEP, oral PrEP offers a unique opportunity to define the immunological effects of tenofovir-emtricitabine without interference by HIV infection, HIV-associated comorbidities or the additional drugs PLWH must take. Because oral use of tenofovir and emtricitabine is more common than topical use of tenofovir by orders of magnitude, lacking information on its mucosal impact in fact constitutes a startling knowledge gap.

To fill this gap, we received and tested blood and mucosal specimens from three human HIV PrEP studies of oral tenofovir-emtricitabine or tenofovir alone, two pre-licensure (NCT00557245, NCT01687218)^8,9^ and one post-licensure (NCT02621242). Participants were HIV-negative, generally healthy and took no additional antiretroviral medications such as integrase or protease inhibitors, allowing us to evaluate the drugs’ effects with minimal confounders. We used transcriptomics and proteomics for broad discovery and confirmed results by focused PCR assays and immunofluorescence staining.

## Results

We measured the effect of oral tenofovir disoproxil fumarate (TDF) and emtricitabine (FTC), or oral TDF alone, on the gastrointestinal tract, the female reproductive tract, and blood from three clinical trials of oral PrEP in HIV-uninfected individuals: the Microbicide Trials Network trial 017^9^ (MTN-017); ACTU-3500 (first reported here, see Methods); and the Genital Mucosal Substudy (GMS)^10^ of the Partners PrEP Study^8^. We analyzed gene expression by microarray, RNA sequencing (RNAseq) and digital droplet PCR (ddPCR); and protein expression by mass spectrometry and microscopy. Paired gastrointestinal biopsies were obtained before and during treatment from two studies: MTN-017 (rectal biopsies) and ACTU-3500 (rectal and duodenal biopsies). Paired female reproductive tract biopsies were obtained during and after treatment from one GMS cohort (GMS A; vaginal and ectocervical biopsies). Paired blood was used from ACTU-3500 and GMS A. Finally, blood was also obtained from a second GMS cohort in order to compare treatment to placebo (GMS B).Samples and assays are described in **Table 1** and in the Methods. RNA quality was assessed for all samples with the RNA integrity number, as presented in Supplemental Table 1.

**Table 1.** Characteristics of studies and cohorts from which samples were obtained. GMS Genital Mucosal Substudy of the Partners PrEP study, MTN-017 Microbicide Trials Network study 017, ACTU-3500 AIDS Clinical Trial Unit Study 3500, TDF tenofovir disoproxil fumarate, FTC emtricitabine, PBMC peripheral blood mononuclear cells.

| Study | Cohort | Gender | Control | Treatment | Drug | Sample | N | Assays |
| --- | --- | --- | --- | --- | --- | --- | --- | --- |
| NCT01687218<br>Ref <sup>9</sup> | MTN-017 | Men (34), trans women (2) | Pre-treatment, paired | 2 months of treatment | TDF/FTC | Rectum | 36 pairs | Microarray<br>RNAseq<br>ddPCR<br>Proteomics |
| NCT02621242 | ACTU-3500 | Men | Pre-treatment, paired | 2 months of treatment | TDF/FTC | Duodenum | 8 pairs | Microarray<br>ddPCR<br>Microscopy |
|  |  |  |  |  | TDF/FTC | Rectum | 8 pairs | Microarray<br>ddPCR<br>Microscopy |
|  |  |  |  |  | TDF/FTC | Whole blood | 8 pairs | Microarray<br>ddPCR |
|  |  |  |  |  | TDF/FTC | PBMC | 8 pairs | Microarray<br>ddPCR |
| NCT00557245<br>Ref <sup>8,10</sup> | GMS A | Women | 2 months after treatment cessation, paired | 24-36 months of treatment | TDF/FTC | Vagina | 3 pairs | Microarray |
|  |  |  |  |  | TDF | Vagina | 12 pairs | Microarray<br>ddPCR |
|  |  |  |  |  | TDF | Ectocervix | 9 pairs | Microarray<br>ddPCR |
|  |  |  |  |  | TDF | PBMC | 10 pairs | Microarray<br>ddPCR |
|  | GMS B | Women | Placebo, unpaired | 24-36 months of treatment | TDF | PBMC | 36 drug<br>20 placebo | Microarray |
|  |  |  |  |  | TDF/FTC | PBMC | 26 drug<br>20 placebo | Microarray |

### Differentially expressed genes

Gene expression was measured with microarrays, comparing no treatment and treatment with oral TDF or TDF/FTC, paired within individuals for studies with multiple samples per participant (MTN-017, ACTU-3500, and GMS A). Differential expression was defined by an FDR-adjusted p-value less than 0.05, with up-regulation meaning higher expression during treatment.

Differential expression was quantified with log2-fold changes. Differentially expressed genes were found in two studies: MTN-017 rectal samples (13 genes up, n = 36 pairs of samples) and ACTU-3500 duodenal samples (116 genes up and 135 genes down, n = 8 pairs of samples) (**Table 2**). No differentially expressed genes were found in any of the other study arms by our chosen FDR-adjusted p-value threshold of 0.05. Gene lists for every study arm and sample type are in Supplemental File 1.

**Table 2.** Differentially expressed genes. Differentially expressed genes are defined by an FDR-adjusted p-value less than 0.05, with up indicating higher expression during drug treatment. All analyses were paired within individuals except for GMS B, where samples from treated individuals were compared to samples from individuals receiving placebo.

| Study | Drug | Sample | Genes up | Genes down | Total genes | Off treatment n | On treatment n |
| --- | --- | --- | --- | --- | --- | --- | --- |
| MTN-017 | TDF/FTC | Rectum | 13 | 0 | 21683 | 36 | 36 |
| ACTU-3500 | TDF/FTC | Duodenum | 116 | 135 | 16321 | 8 | 8 |
| ACTU-3500 | TDF/FTC | Rectum | 0 | 0 | 16399 | 8 | 8 |
| ACTU-3500 | TDF/FTC | Whole blood | 0 | 0 | 13922 | 8 | 8 |
| ACTU-3500 | TDF/FTC | PBMC | 0 | 0 | 14937 | 8 | 8 |
| GMS A | TDF/FTC | Vagina | 0 | 0 | 18397 | 3 | 3 |
| GMS A | TDF | Vagina | 0 | 0 | 18397 | 12 | 12 |
| GMS A | TDF | Ectocervix | 0 | 0 | 17578 | 9 | 9 |
| GMS A | TDF | PBMC | 0 | 0 | 16553 | 10 | 10 |
| GMS B | TDF | PBMC | 0 | 0 | 20121 | 20 | 36 |
| GMS B | TDF/FTC | PBMC | 0 | 0 | 20121 | 20 | 26 |

All 13 genes differentially expressed in the rectum in MTN-017 were expressed more highly during treatment with TDF/FTC. As shown in **Table 3**, seven of these 13 genes are members of the gene ontology biological process “type I interferon signaling pathway” (GO:0060337): IFI27, IFI6, IFIT1, ISG15, RSAD2, MX1 and OAS1. Four of the other six have been reported in the literature to be induced by type I interferon: DDX60^11,12^, SAMD9^11^, IFI27L1^13^, and HERC6^11^. Thus, only two of the thirteen genes (CCDC77 and the pseudogene MROH3P) have no reported roles related to type I interferon. Gene ontology overrepresentation analysis of these thirteen genes revealed that these thirteen genes were highly overrepresented in biological processes related to type I interferon and response to virus (Supplemental File 1).

**Table 3.**
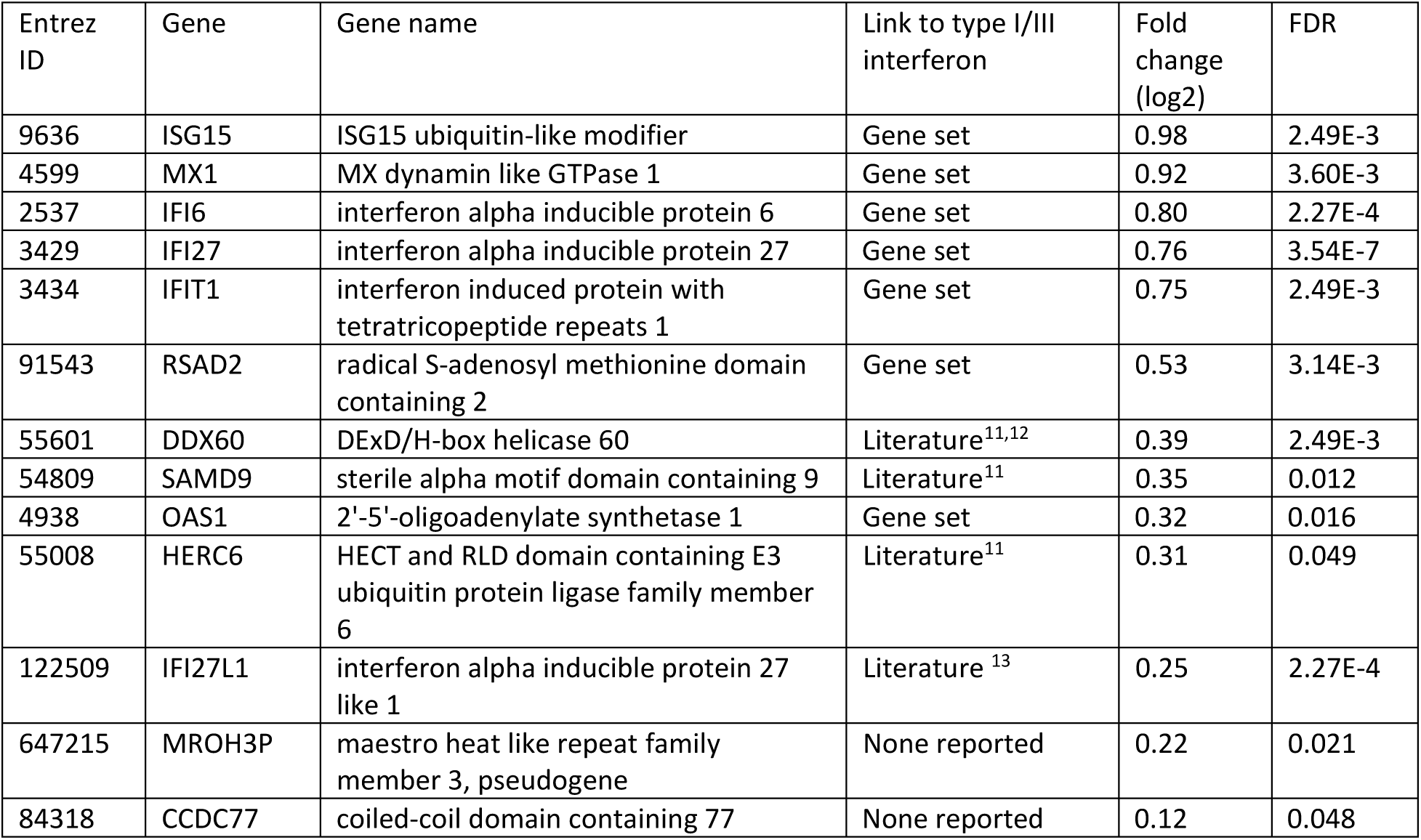
Genes differentially expressed (higher during daily oral TDF/FTC use) in rectal biopsies in MTN-017. “Gene set” indicates membership in GO:0060337. “Literature” indicates that a link to type I/III interferon has been reported in the indicated articles.

| Entrez ID | Gene | Gene name | Link to type I/III interferon | Fold change (log2) | FDR |
| --- | --- | --- | --- | --- | --- |
| 9636 | ISG15 | ISG15 ubiquitin-like modifier | Gene set | 0.98 | 2.49E-3 |
| 4599 | MX1 | MX dynamin like GTPase 1 | Gene set | 0.92 | 3.60E-3 |
| 2537 | IFI6 | interferon alpha inducible protein 6 | Gene set | 0.80 | 2.27E-4 |
| 3429 | IFI27 | interferon alpha inducible protein 27 | Gene set | 0.76 | 3.54E-7 |
| 3434 | IFIT1 | interferon induced protein with tetratricopeptide repeats 1 | Gene set | 0.75 | 2.49E-3 |
| 91543 | RSAD2 | radical S-adenosyl methionine domain containing 2 | Gene set | 0.53 | 3.14E-3 |
| 55601 | DDX60 | DExD/H-box helicase 60 | Literature<sup>11,12</sup> | 0.39 | 2.49E-3 |
| 54809 | SAMD9 | sterile alpha motif domain containing 9 | Literature<sup>11</sup> | 0.35 | 0.012 |
| 4938 | OAS1 | 2'-5'-oligoadenylate synthetase 1 | Gene set | 0.32 | 0.016 |
| 55008 | HERC6 | HECT and RLD domain containing E3 ubiquitin protein ligase family member 6 | Literature<sup>11</sup> | 0.31 | 0.049 |
| 122509 | IFI27L1 | interferon alpha inducible protein 27 like 1 | Literature<sup>13</sup> | 0.25 | 2.27E-4 |
| 647215 | MROH3P | maestro heat like repeat family member 3, pseudogene | None reported | 0.22 | 0.021 |
| 84318 | CCDC77 | coiled-coil domain containing 77 | None reported | 0.12 | 0.048 |

In the duodenum in ACTU-3500, the top overrepresented biological processes for down-regulated genes were related to cellular metabolism, and for up-regulated genes were related to a variety of biological processes including RNA splicing and phospholipid transport (Supplemental File 1).

### Correlation of gene expression across study arms

The log2-fold changes of the 13 differentially expressed genes from the rectum in MTN-017 strongly correlated with the fold changes of these same genes in the rectum in ACTU-3500 (**Figure 1A**, left) (Spearman correlation coefficient r=0.91, as compared to 0.07 for all other genes). None of these genes had adjusted p-values < 0.05 in the rectum in ACTU-3500 (two had unadjusted p < 0.05), possibly due to the much smaller sample size.

**Figure 1.**
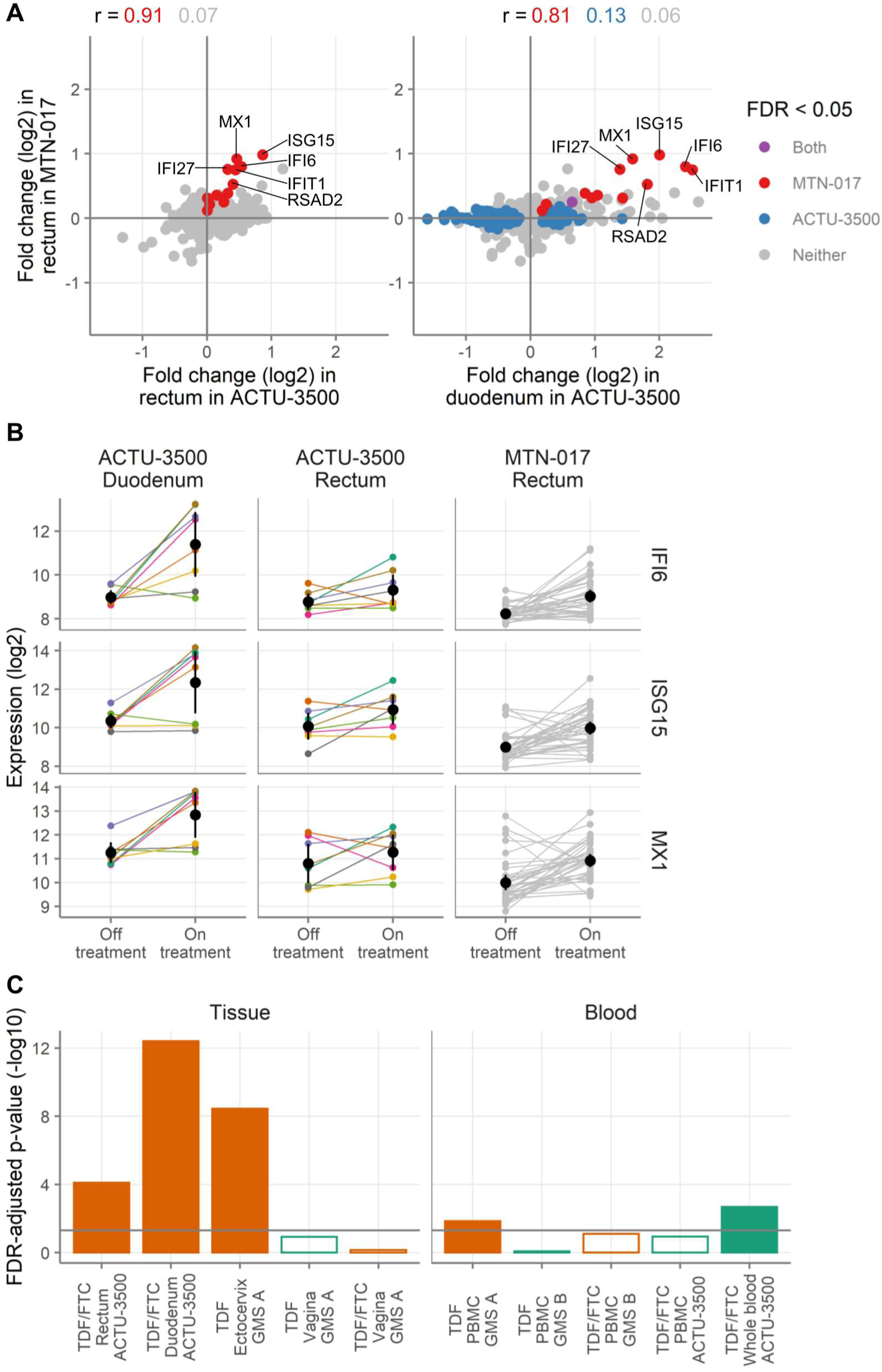
Gene expression across study arms by microarray analysis. **(A)** Fold changes of all genes detected in the rectal samples in MTN-017 compared to the rectal samples from ACTU-3500 (left) and the duodenal samples from ACTU-3500 (right). Colors indicate genes with FDR-adjusted p-values less than 0.05 in MTN-017 (red), ACTU-3500 (blue), both (purple), or neither (gray). Spearman correlation coefficients for the genes falling into each subset are shown. **(B)** Expression levels of IFI6, ISG15, and MX1 in individual samples. Small points indicate measurements from a single biopsy, with lines connecting the matching observation from the same donor. For ACTU-3500, the color of the lines and symbols indicate the participant and are consistent across the panels. Black symbols and vertical lines show the mean and 95% confidence interval of the mean. **(C)** Gene set testing of a custom gene set composed of differentially expressed genes from the rectum in MTN-017 performed in the other study arms in the mucosa (left) and blood (right), comparing expression of genes in the set to all other detected genes within each study arm. Bars indicate the result of a test against the study labeled on the x-axis. Filled bars indicate an FDR-adjusted p-value less than 0.05 and open symbols the opposite, with bar height showing the -log10 of the FDR-adjusted p-value. Colors indicate the direction of change, with orange meaning more expression during product use and green meaning the opposite. The horizontal grey line shows an FDR-adjusted p-value of 0.05.

Similarly, there was a strong correlation between the fold changes of the 13 differentially expressed genes in the rectum from MTN-017 with the fold changes of the same genes in the duodenum in ACTU-3500 (**Figure 1A**, right) (Spearman correlation coefficient r=0.81, as compared to 0.13 for all genes that were differentially expressed in the duodenum, and 0.06 for all other genes). Only one of these genes had an adjusted p-value < 0.05 in the duodenum in ACTU-3500 (though all 13 had unadjusted p < 0.05), possibly due to the much smaller sample size.

Taken together, these results show that oral TDF or TDF/FTC affects the expression of relatively few genes. In particular, we did not find evidence of differential gene expression in the blood. We did detect differentially expressed genes in the rectum and the duodenum. The 13 genes that were differentially expressed in the rectum in MTN-017 were strongly correlated in the duodenum and rectum in ACTU-3500. We additionally performed gene set testing across each other study arm using these 13 genes as a custom gene set, comparing their expression with the expression of genes outside of the set. This custom 13-gene set derived from the rectum in MTN-017 was strongly enriched in the rectum and duodenum in ACTU-3500 (FDR = 8E-5 and 4E-13, **Figure 1B** and complete gene set testing results in Supplemental File 1), as well as in the ectocervix. This suggests that those 13 genes, 11 of which are known to be type I/III interferon-related, may represent an underlying biological pathway that is affected by TDF/FTC in the gastrointestinal tract.

### Gene set testing of Hallmark gene sets

To assess higher level biological effects, we performed gene set testing on the fifty Hallmark gene sets^14^ within each study arm and specimen type, comparing the expression of genes in each set to the expression of genes not in the set (i.e. all other detected genes). Each gene set comprises genes that are involved in a biological state or process. To reduce the number of gene sets displayed and focus on those gene sets that were affected in multiple studies, we show the gene sets that had adjusted p-values below 0.05 in at least two study arms for tissue and blood in **Figure 2A-B**. Complete gene set testing results are provided in Supplemental File 1. For tissue (**Figure 2A**), thirteen gene sets had adjusted p-values below 0.05 in at least two study arms. Of these gene sets, four were related to immunity (allograft rejection, interferon-α and -γ responses, and TNF-α signaling via NF-κB), and four were related to cell proliferation (E2F targets, G2M checkpoint, and two MYC targets gene sets). In the mucosa, the immune related gene sets tended to be elevated during product use, with the interferon-α response being the strongest, not only in the duodenum and rectum but also the ectocervix. Only two gene sets had adjusted p-values <0.05 in at least two study arms for blood samples (**Figure 2B**). Both gene sets were immune-related (complement, interferon-γ response, and TNF-α signaling via NF-κB). In both cases, these gene sets were lower during product use. Thus, TDF/FTC seemed to induce interferon-α responses in the gastrointestinal tract and ectocervix but had a somewhat dampening effect on inflammatory responses in the blood.

**Figure 2.**
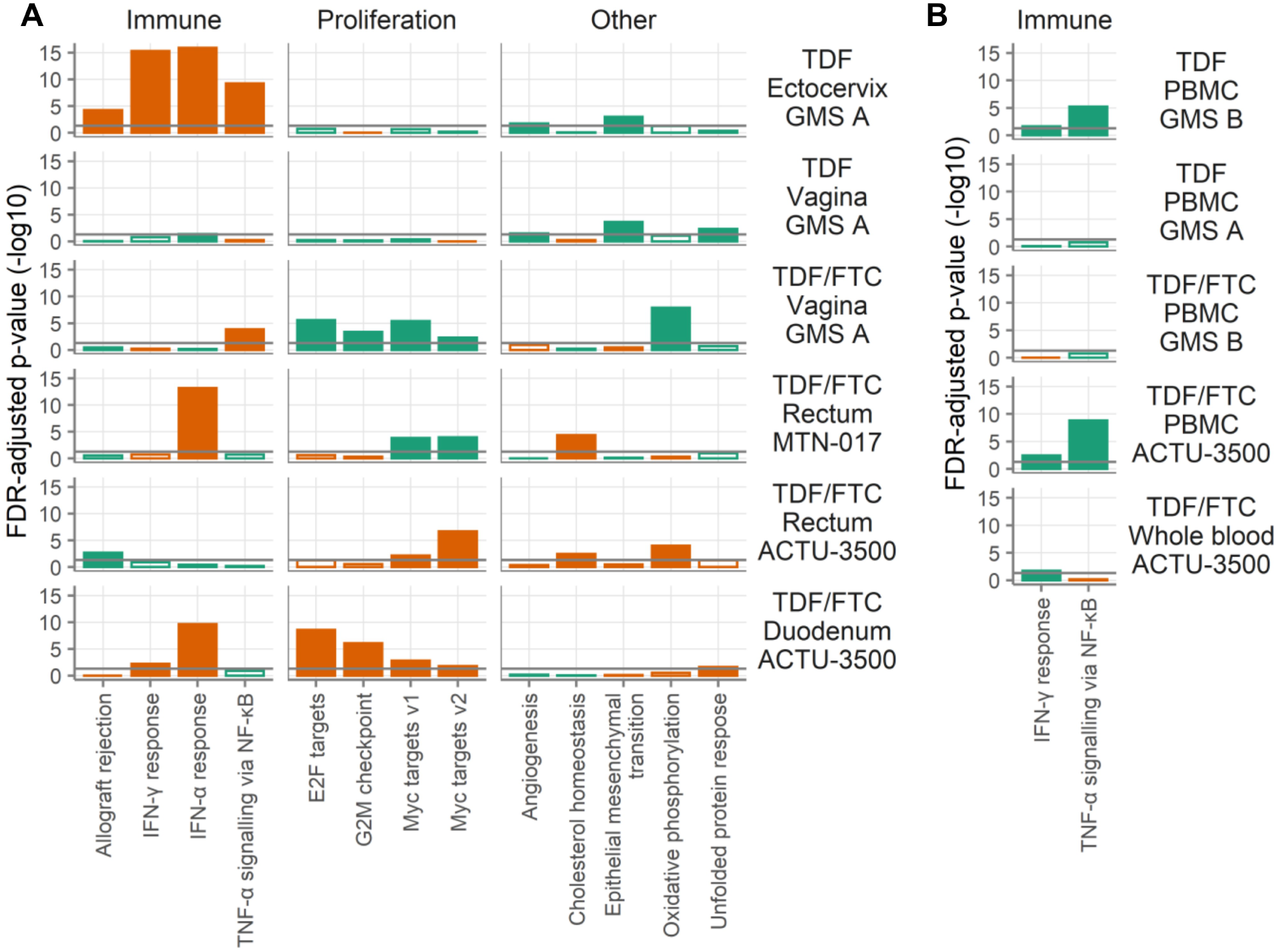
Hallmark gene sets. Gene set testing was performed on the Hallmark gene sets in the mucosa **(A)** and the blood **(B)**. To reduce the number of gene sets displayed and focus on those gene sets that were affected in multiple studies, we show only those gene sets that had an FDR-adjusted p-value less than 0.05 in at least two of the mucosa (A) or blood (B) study arms. Bars indicate the result of a gene set test for the gene set shown on the x-axis tested against the study shown at right. Filled bars indicate an FDR-adjusted p-value less than 0.05 and open bars the opposite, with bar length proportional to the -log10 of the FDR-adjusted p-value. Colors indicate the direction of change, with orange meaning more expression during product use and green meaning less. The horizontal grey lines show an FDR-adjusted p-value of 0.05. The gene sets are grouped into categories as labeled at the top.

### Corroboration of microarray data by RNA sequencing, mass spectrometry and ddPCR

In addition to microarrays, we analyzed the RNA from MTN-017 participants by RNA sequencing. Two genes were differentially expressed (FDR-adjusted p<0.05) in the RNAseq data, both higher during TDF/FTC treatment than at baseline: IFIT1, which was also differentially expressed by microarray, and SLC6A20, which was not. Fold expression changes of the 13 genes identified as significantly upregulated in the microarrays correlated strongly with their fold expression changes by RNAseq (Spearman correlation 0.84 as compared to 0.34 for all other genes, **Figure 3A**). By virtue of having both microarray and RNAseq data on the same samples, we were able to look at genes that had similar fold changes by both methodologies. We looked at genes that had log2-fold changes below -0.25 or above 0.25 by both assays. Only twelve genes fell into the downregulated group and none had adjusted p-values below 0.05 by either assay but, strikingly, eight were metallothioneins (MT1A, 1E, 1F, 1G, 1H, 1M, 1X, and 2A), which bind to heavy metal ions. There were 28 genes with fold changes above 0.25 by both microarray and RNAseq, including all the differentially expressed genes from the microarray, except for the two non-type-I/III-interferon-related genes (i.e., 11 of 13 microarray genes). Many of the additional genes were also related to type I/III interferon signaling: IRF7, IRF9, OAS2, OAS3, IFITM1, and IFI44L, for example. Overrepresentation analysis of these 28 genes (with or without the 11 differentially expressed genes from the microarray) again yielded many gene ontology biological processes related to type I/III interferon signaling.

**Figure 3.**
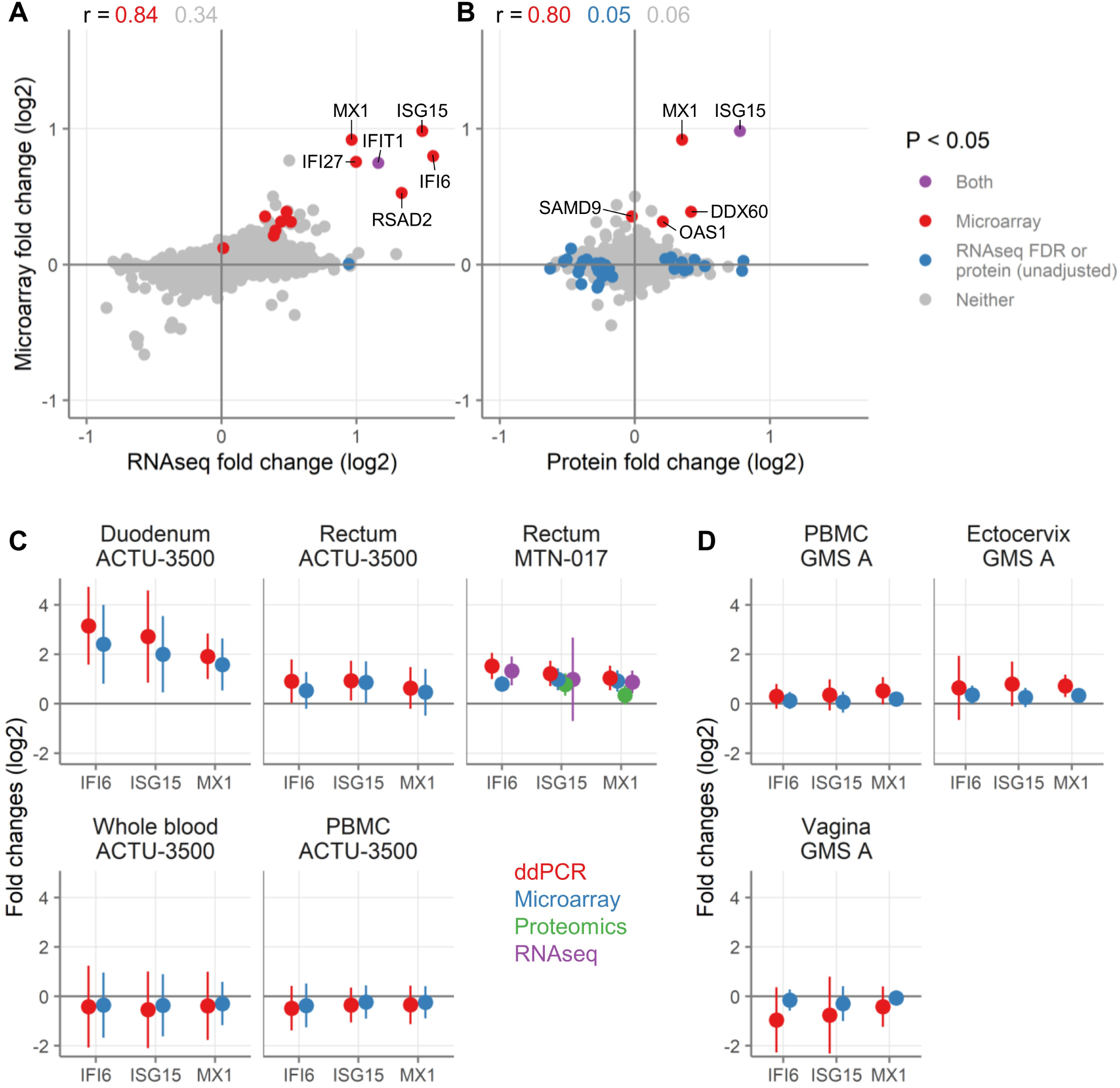
Comparison of gene expression changes measured by ddPCR, RNAseq, proteomics, and microarray. **(A-B)** Correlation of fold changes of genes as detected by microarray (y-axis) with genes detected by RNAseq **(A)** or proteins detected by proteomics **(B)** from the rectal samples from MTN-017. Colors indicate genes with FDR-adjusted p-values less than 0.05 in microarray transcripts (red), RNAseq genes (blue, by FDR, A) or proteins (blue, unadjusted p-value, B), both (purple, FDR for microarray and RNAseq, unadjusted p-value for protein), or neither (gray). Spearman correlation coefficients for the genes falling into each subset are shown. Selected genes are labeled. **(C-D)** Fold changes in gene expression of three genes (IFI6, ISG15, or MX1) as detected by ddPCR (red), microarray (blue), proteomics (green), and RNAseq (purple) after treatment with TDF/FTC **(C)** or TDF alone **(D)**. Symbols show the mean across all participants and vertical lines show the 95% confidence intervals of the mean. IFI6 stands for interferon alpha inducible protein 6, ISG15 for ISG15 ubiquitin-like modifier, and MX1 for MX dynamin like GTPase 1. A positive fold change means higher expression during treatment and a negative fold change means higher expression off of treatment. For ddPCR, the expression of each gene of these three genes was normalized to the expression of ubiquitin C (UBC), which was chosen as reference due to the stability of its expression across tissues and treatments in the microarray data.

We also analyzed rectal biopsies from MTN-017 by mass spectrometry for protein identification. The biopsies were run in two batches, with the samples from US participants in one batch and samples from Thai participants in the other. In both cases, no proteins were detected as differentially expressed after adjustment for multiple comparisons (complete list of protein fold changes in Supplemental File 1). None of the proteins from the 13 genes of interest was consistently detected in the US participants, due to the low abundance of these proteins. We therefore could not analyze the US batch for these proteins. However, five of these proteins (ISG15, MX1, OAS1, DDX60 and SAMD9) were detected in all Thai samples. All but one (SAMD9) had positive fold changes and there was a strong correlation between the fold changes detected by microarray and mass spectrometry (Spearman 0.80; **Figure 3B**).

Finally, we used ddPCR to measure the expression of three selected IFN-I/III genes (IFI6, ISG15, and MX1). The fold changes for ddPCR and microarray are shown in **Figure 3C-D**. By ddPCR, linear scale fold changes for these three genes ranged from 3.77 to 8.89 in the duodenum and 1.56 to 2.88 in the rectum, while they were little changed in the blood, ectocervix, and vagina. Changes measured by ddPCR were always in the same direction as those by microarray and the magnitudes tended to be larger by ddPCR. The participant-level fold changes calculated from ddPCR measurements correlated well with the microarray data (Pearson correlation 0.93 for MX1, 0.92 for ISG15, and 0.90 for IFI6). Taking each gene separately and stratifying by study, sample type and gene, Pearson correlations ranged from 0.54 to 0.99, with a mean of 0.86 and median of 0.92. Thus, the ddPCR data confirmed that oral TDF/FTC induces genes related to type I/III interferon throughout the gut and mildly in the ectocervix, but not the vagina and blood.

Taken together, the data obtained from the rectum and the duodenum after TDF/FTC treatment for two months, in two different studies (MTN-017 and ACTU-3500), and by several different assays (microarray, RNAseq, ddPCR, and mass spectrometry-based proteomics) all indicate upregulation of factors related to type I/III IFN signaling in the gastrointestinal tract.

### Immunofluorescence microscopy of gastrointestinal tissues

To further assess type I/III interferon induction, we stained duodenal and rectal tissue sections from ACTU-3500 for ISG15. Slides were evaluable from 8 pairs of duodenal biopsies and 6 pairs of rectal biopsies (before and during treatment). ISG15 was expressed by few if any stromal cells but within the columnar epithelium, some cells stained intensely positive (**Figure 4A**). Whereas the intensity of ISG15 staining did not change with TDF/FTC use (**Figure 4B**), the mean percentage of ISG15 bright cells increased in both the rectum (increase of 0.43 percentage points or 2.76-fold, one-sided paired t-test p = 0.0023) and the duodenum (0.43 percentage points or 1.37-fold, one-sided paired t-test p = 0.06) (**Figure 4C**). Co-staining of duodenal biopsies from three individuals with ISG15 and glycoprotein 2 (GP2), a cell surface receptor reported to identify microfold (M) cells, a special type of intestinal epithelial cell with heightened immunological activity^15-17^, revealed a startling overlap between the two markers (**Figure 4D**). Thus, TDF/FTC use increased the proportion of cells expressing high levels of ISG15, and at least some of these cells may be M cells.

**Figure 4.**
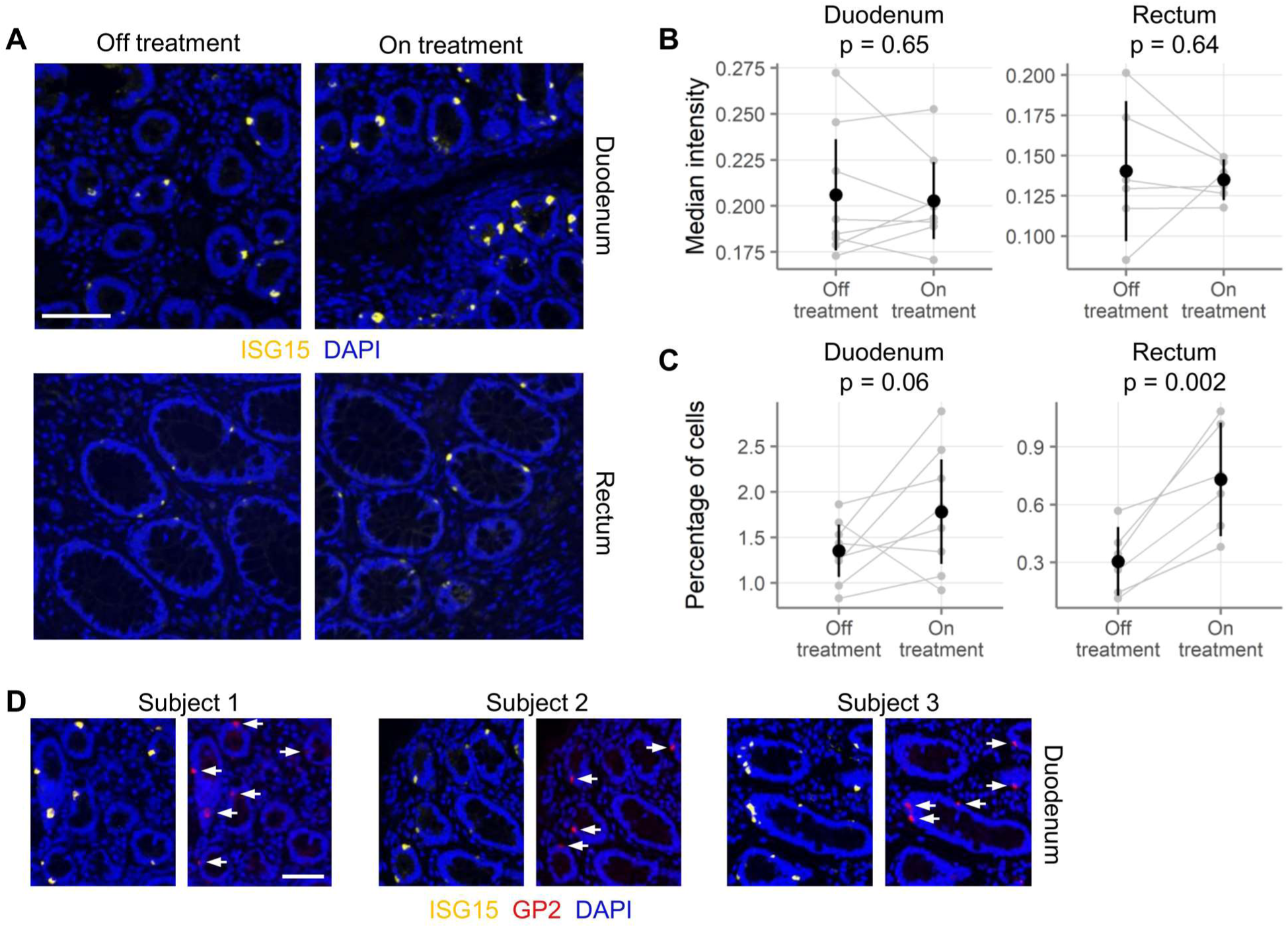
Immunofluorescence microscopy staining for ISG15. **(A)** 20X magnification images of duodenal (top) and rectal (bottom) biopsies, stained for ISG15 (yellow) and DAPI (blue). Biopsies from pre-treatment (left) and at the end of two months of treatment (right) are shown. Scale bars are 100 µm. Duodenal biopsies came from one donor and rectal biopsies came from a second. **(B)** ISG15 intensity on ISG15 positive cells was measured in paired duodenal (n = 8 donors) and rectal (n = 6) biopsies from ACTU-3500. The median intensity of all cells measured is shown. **(C)** The percentage of ISG15 positive cells out of all epithelial cells is shown for the same biopsies as in (B). In (B) and (C), gray points indicate measurements from a single biopsy, with gray lines connecting the matching observation from the same donor. Black symbols and vertical lines show the mean and 95% confidence interval of the mean. **(D)** Co-staining of duodenal biopsies from three individuals for ISG15 and glycoprotein 2 (GP2). Anti-GP2 was raised against a peptide component of pancreatic secretory (zymogen) granules and has some cross-reactivity with microfold (M) cells. GP2 positive cells co-express ISG15 (white arrows) but not all ISG15 positive cells are GP2 positive. Scale bar is 100 µm.

## Discussion

We found that oral treatment with TDF/FTC or TDF alone had limited global effects on host gene expression, with no differentially expressed genes in the blood or female reproductive tract and tens to hundreds in the gastrointestinal tract. Notably, though, genes related to type I/III interferon signaling were consistently induced in the gut, with good agreement between microarray, ddPCR and RNA sequencing. Protein-level data by mass spectrometry, and immunohistology of gut sections for ISG15, were in congruence.

The overall limited changes we found indicate that TDF/FTC have few off-target effects on host gene expression, suggesting that TDF/FTC treatment is largely benign. In light of the widespread use of TDF/FTC among HIV-infected individuals for treatment and HIV-uninfected individuals for prevention, this is reassuring. Moreover, increased type I/III interferon signaling could signify increased innate immune readiness, enhancing the anti-viral preventative or treatment efficacy of TDF/FTC. In fact, induction of a type I/III interferon signature by TDF/FTC primarily in the gastrointestinal mucosa could contribute to oral PrEP’s greater efficacy in preventing rectal over vaginal HIV transmission^18^. Other proposed explanations for this observation include pharmacokinetic differences in tenofovir levels between vaginal and rectal tissues^19^ and perturbations of tenofovir metabolism by a dysbiotic vaginal microbiome^20^.

Stimulation of interferon pathways by TDF/FTC could have detrimental effects too. It is well known that continued disinhibition of IFN-I/III responses plays an important role in chronic inflammatory diseases^21-23^. Long-term use of TDF/FTC with ongoing stimulation of IFN-I/III pathways could predispose individuals to chronic immune activation, including people living with HIV (PLWH). Interestingly, two recent papers in HIV-infected humanized mice showed that blocking IFN-I/III pathways lead not only to less immune activation but also to a lower HIV reservoir and delayed HIV rebound following ART interruption^24,25^. Oral TDF/FTC in PLWH could therefore paradoxically promote HIV reservoir persistence, especially in the gut where >99% of the latent reservoir is thought to reside^5^ and where TDF/FTC’s effect is strongest. Thus, selecting ART drugs that don’t stimulate IFN-I/III might decrease morbidity associated with chronic immune activation and even contribute to HIV cure strategies, which will likely be delivered alongside suppressive ART. However, we emphasize that these possible consequences of TDF/FTC treatment in PLWH remain speculative because we did not study these drugs specifically in PLWH. Notably, other factors may cause or contribute to persistent immune activation, such as dysregulation of the gut microbiome (reviewed in ^26^), dysfunction of regulatory immune cells^27,28^, insults created by HIV infection during the untreated disease phase^29^, and a number of other possible mechanisms (reviewed in ^30^).

In addition to changes to immune-related gene expression in the gastrointestinal tract, the other notable gene expression changes were related to cell proliferation. Specifically, we saw some evidence of increased expression of gene sets related to cell proliferation in the duodenum, and reduced expression of these gene sets in the vagina. Proliferation-related gene sets were conflicting in the rectum (two sets up in one study and down in the other). In a prior study of rectal tenofovir 1% gel, we noted induction of pro-proliferative pathways on both the transcriptomic and proteomic level^7^. It is difficult to speculate about the clinical relevance of changes to cell proliferation pathways. Factors associated with cell cycle regulation are crucial to balance cell proliferation with cell death, and for cells to respond to DNA damage. TDF/FTC’s effect on cell cycle processes in the duodenum may not be surprising given the increased drug concentrations likely achieved in the upper gastrointestinal tract. Long-term, high concentration oral administration of TDF to mice has been reported to cause a low incidence of duodenal tumors (Canadian product monograph for VIREAD) and liver adenomas (US prescribing information for VIREAD). It is unclear whether our findings with human duodenal biopsies relate in any way to these outcomes in rodents and no such findings have been reported during human use.

It remains speculative how TDF/FTC induce a type I/III signature in the gut, but our and another group’s recent data point to some possible mechanisms. First, TDF/FTC increased the relative number of ISG15 bright cells within the columnar epithelium of the gut but not their ISG15 expression levels (Figure 4), suggesting that the drugs have a proliferation-inducing effect especially on these cells. Based on co-expression of glycoprotein 2, they may be M cells, which are highly active immunological sentinels of the intestinal mucosa^15-17^. The increased numbers of these cells suggests that the increased ISG activity we observed is due to a greater number of ISG-producing cells, rather than more ISGs produced by a constant number of cells. Second, TDF treatment has been reported to increase serum IFN-λ3 levels in patients with hepatitis B or HIV as well as IFN-λ3 secretion from cultured colon cancer cell lines^31^, indicating a role for type III IFN. These experiments showed that TDF induction of IFN-λ3 production is unique to gut cells and, further, that TDF-induced IFN-λ3-containing supernatants cause production of MX1 and OAS2. Thus, these experiments provide direct *in vitro* confirmation of our findings that TDF/FTC induce type I/III IFN production specifically in the gastrointestinal tract. Unfortunately, neither type I nor type III interferons were detectable in any of our samples, likely due to inadequate sensitivity of the assays. Lastly, there have been several reports, including our prior finding ^7,32,33^, that TDF inhibits IL-10 production. It has recently been shown that tenofovir monophosphate, an intracellular metabolite of TDF, strongly binds to Akt, a protein kinase broadly involved in intracellular signaling, including lipopolysaccharide (LPS)-induced stimulation of interleukin 10 (IL-10) transcription. Binding by tenofovir monophosphate prevents Akt phosphorylation and translocation to the plasma membrane, interrupting a key event in the LPS/IL-10 signaling cascade^32^. By consequence, TDF treatment reduces anti-inflammatory IL-10 responses to LPS in favor of pro-inflammatory IL-12 responses. Taken together, these data demonstrate specific intracellular effects of tenofovir, with consequences ranging from disinhibition of cellular proliferation to perturbation of innate cytokine networks. However, more studies will be required to tie these findings together in a unified functional model.

Our study has a few limitations. One is the use of within-person comparisons between on- and off-drug. Because participants were aware of when they were or were not taking an intervention, behavioral changes or other factors than the drug itself could explain the gene expression changes we observed. The only study where placebo was compared to treatment is the GMS B study, in which the samples were PBMC and no differentially expressed genes were observed. Secondly, co-existing or new bacterial or viral infections could explain the interferon responses we attributed to the effect of TDF/FTC use. However, it is relatively unlikely for infections to explain these effects, because they would have to occur in a concerted fashion primarily at the end of the study period, during TDF/FTC use, and rarely at baseline, before TDF/FTC initiation. Moreover, the underlying clinical trials were well monitored and included a comprehensive package of HIV prevention counseling and sexually transmitted infection testing throughout the studies. No increase of STIs or other infections was noted during TDF/FTC use by testing or clinical symptoms. Thirdly, we looked for gene expression changes at the bulk cell levels of tissue, PBMC, and whole blood. Had we looked at specific cell types, we may have seen different results. For example, there is evidence that PrEP alters the composition of immune cells in tissues in the female genital tract and the blood^33^. We were also limited in our ability to differentiate effects of FTC from those of TDF, given that only a limited set of participants received TDF alone and none received FTC alone. Finally, and most importantly, our results are limited to HIV-uninfected people, so it is unclear whether our findings extend to PLWH. With the relatively recent emergence of NRTI-sparing, but equally effective, cART regimens^34,35^, it has become possible to conduct a prospective clinical trial comparing immune activation and HIV reservoir decay in PLWH randomized to NRTI-containing versus NRTI-sparing regimens. Our data advocate for such a study.

## Methods

### Studies

Samples were used from four studies, which are described in **Table 1**. Two sets of samples were used from the Genital Mucosal Substudy (GMS)^10^ of the Partners PrEP Study^8^: paired samples during and after treatment (GMS A) and unpaired placebo vs. treatment samples (GMS B). The samples all came from the same parent study, but were processed separately. The Microbicide Trials Network trial 017 (MTN-017) included oral TDF/FTC as well as topical tenofovir; only the oral TDF/FTC samples are included in this analysis. Rectal biopsies taken after two months of oral TDF/FTC use were compared to baseline samples. The study visits for the samples used here were performed at two sites (Pittsburgh, USA and Bangkok, Thailand), with half of the samples coming from each site. ACTU-3500 followed nine men initiating oral PrEP with TDF/FTC in Seattle, with a baseline visit and a visit after two months of PrEP use. Eight men completed both visits. Ethics reviews are published in the primary manuscripts for the GMS and MTN-017 studies (listed in **Table 1**). ACTU-3500 was reviewed through the University of Washington Institutional Review Board, number 49167. Sample sizes varied within each trial depending on drug (TDF/FTC or TDF) and sample type. Complete sample size information is listed in **Table 1**. Study adherence measures are given in the Supplemental Material and Methods.

### Study sampling and laboratory assays

Blood and biopsy sampling are described in the Supplemental Material and Methods. Gene expression was measured at the level of RNA and protein, by the methods indicated in **Table 1** and described in detail in the Supplemental Material and Methods.

### Data analysis

Data were initially processed using instrument-specific software as described in the above sections. Following export from instrument-specific software, data were analyzed using R version 3.5.2^36^. Microarray data is posted on GEO at accession numbers GSE139611, GSE138723, GSE139655, and GSE139411. The raw data for all other assays is available on figshare at doi: 10.6084/m9.figshare.c.4704827. All code necessary to reproduce the analyses and figures is provided in Supplemental File 2.

## Supporting information

Results of analyses

Code to reproduce analyses and figures

## Acknowledgments

We wish to express gratitude to all study volunteers for their participation. We are grateful to Max Abou and Lauren Girard for proteomic wet lab support as well as Stuart McCorrister and Garrett Westmacott for mass spectrometry technical support. We acknowledge the Fred Hutch Experimental Histopathology (Sunni Farley, Savanh Chanthaphavong, Li-Ya Huang) and Genomics (Cassie Sather, Crissa Bennett) core facilities for their assistance.

This work was funded by NIH R01AI116292 (to FH), NIH R01AI111738 (to JRL), Bill and Melinda Gates Foundation #47674 (to JRL), NIH R01AI134293 (to RM), NIH AI027757 (to JMB), NIH AI069481 (to ACC and MJM). MJC and IM were supported by the Microbicide Trials Network (UM1AI068633, Sharon Hillier, PI). The ddPCR portion of this work was supported by a grant from the James B. Pendleton Charitable Trust. The NCI 5 P30 CA015704-44 Cancer Center Support Grant supported the Fred Hutch Experimental Histopathology core facility.

## Disclaimer

The findings and conclusions in this report are those of the authors and do not necessarily represent the official position of the funding agencies. The funders had no role in the study design, data collection and analysis, decision to publish, or preparation of the manuscript. The corresponding author had full access to all the data in the study and had final responsibility for the decision to submit for publication.

## Supplement

**Supplemental File 1**. Results of data analysis.

**Supplemental File 2**. R code used for data analysis.

## Supplemental Tables

**Supplemental Table 1.** Sample quality. RNA Integrity Number (RIN) was determined by Agilent TapeStation. Values are displayed as mean (range).

| Study | Sample | RIN |
| --- | --- | --- |
| ACTU-3500 | Duodenum | 8.4 (7.8-9) |
| ACTU-3500 | Whole blood | 8.1 (7.1-8.7) |
| ACTU-3500 | PBMC | 9.6 (9.3-9.8) |
| ACTU-3500 | Rectum | 7.9 (6.9-8.8) |
| GMS A | Ectocervix | 8.1 (7.1-8.8) |
| GMS A | PBMC | 8.3 (6.5-9.4) |
| GMS A | Vagina | 8.3 (6.5-9.7) |
| GMS B | PBMC | 9.1 (7-9.7) |
| MTN-017 | Rectum | 7.8 (5.5-8.8) |

**Supplemental Table 2.**
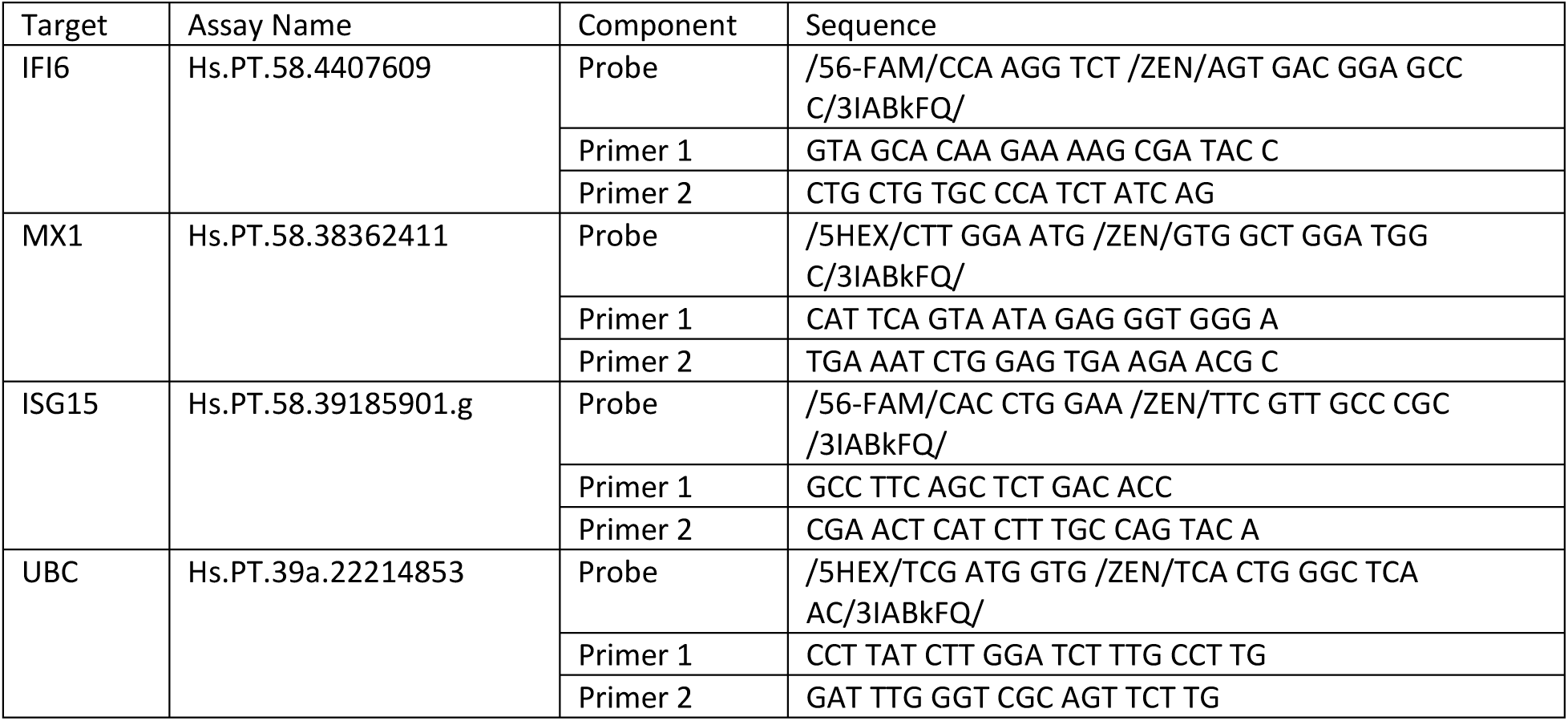
Primers and probes used for ddPCR.

## Notes

https://www.ncbi.nlm.nih.gov/geo/query/acc.cgi?acc=GSE139611

https://www.ncbi.nlm.nih.gov/geo/query/acc.cgi?acc=GSE138723

https://www.ncbi.nlm.nih.gov/geo/query/acc.cgi?acc=GSE139655

https://www.ncbi.nlm.nih.gov/geo/query/acc.cgi?acc=GSE139411

https://doi.org/10.6084/m9.figshare.c.4704827

### Competing Interest Statement

JMB is on advisory boards for Gilead Sciences, Merck and Janssen. IM is the Chief Medical Officer of Orion Biotechnology. All other authors report no conflicts.

