## Supplemental methods for "Treatment with commonly used antiretroviral drugs induces a type I/III interferon signature in the gut in the absence of HIV infection"

### Adherence

Tenofovir levels were measured in the serum for the GMS A and GMS B studies as previously described (Baeten et al., 2014). Samples from the treatment arm without detectable tenofovir were removed, as was one post-treatment sample, where drug was unexpectedly detected. Tenofovir levels were measured in serum in the MTN-017 study. Adherence was high in the oral arm of this study, with 94% of participants taking the daily pill at least 80% of the time (Carballo-Diéguez et al., 2017). Participant report of pill use was largely concordant with serum tenofovir levels, with only 4.4% of serum samples having no detectable tenofovir when participants reported product use. All samples from this study were included. Adherence was determined for the ACTU-3500 study by participant self-report. All participants reported daily use of TDF/FTC throughout the study period.

### Samples and sample storage

Vaginal, cervical, and rectal biopsies were obtained in GMS A and B and MTN-017 as described in the primary manuscripts (Baeten et al., 2012; Cranston et al., 2017; Lund et al., 2016). Rectal biopsies were obtained in ACTU-3500 by anoscopy using Radial Jaw 4 biopsy forceps (Boston Scientific). Duodenal biopsies were obtained by esophagogastroduodenoscopy under light anesthesia also using Radial Jaw 4 biopsy forceps.

Biopsies were placed into RNALater Stabilization Solution (ThermoFisher Scientific, Waltham, MA, USA) and held at 4$^{o}$C for 24 h and then frozen at -80$^{o}$C. PBMC were isolated from whole blood by density gradient centrifugation and then cryopreserved and stored in the vapor phase of a liquid nitrogen freezer. Whole blood was drawn into PAXgene tubes (PreAnalytiX, Hombrechtikon, Switzerland), which were frozen at -20°C for 24 h and then stored at -80°C.

**RNA extraction and quality control**

Biopsies were thawed at room temperature. They were transferred with forceps into 600 µL of Buffer RLT (Qiagen, Hilden, Germany) and homogenized using a Bio-Gen PRO200 homogenizer (PRO Scientific, Oxford, CT, USA) followed by passing 10 times through a needle and syringe. RNA was extracted using the RNeasy fibrous tissue mini kit (Qiagen) automated on a QIAcube (Qiagen). PBMC were thawed, washed by centrifugation, counted, and RNA was extracted using the RNeasy mini kit (Qiagen) on a QIAcube. PAXgene tubes were thawed and held at room temperature for 3 h with occasional mixing, in order to completely lyse red blood cells. RNA was extracted from PAXgene samples using the PAXgene Blood RNA kit (PreAnalytiX) according to the manufacturer's instructions. RNA was stored at -80°C until use.

**Sample quality control**

Cell viability was measured prior to RNA extraction for PBMC samples (Guava Viacount, EMD Millipore, Burlington, MA, USA). The quality of all RNA was determined using the RNA integrity number as calculated from the TapeStation R6K assay (Agilent, Santa Clara, CA, USA, Supplemental Table 1) and the concentration was determined by NanoDrop (ThermoFisher).

**Microarray labeling and hybridization**

For the GMS A and GMS B studies, samples were prepared for microarray using 50 ng of total RNA with the Ovation PicoSL WTA System V2 kit (NuGEN, San Carlos, CA, USA) and labeled with the Encore BiotinIL kit (NuGEN). The Illumina TotalPrep RNA Amplification kit (ThermoFisher) was used to prepare samples for microarray from the ACTU-3500 study (275 ng total RNA as input) and the MTN-017 study (500 ng input), with input sizes chosen based on available RNA.

750 ng of the labeled cDNA (from the NuGEN kits) or cRNA (from the Thermo kit) was hybridized to HumanHT-12 v4 Expression BeadChips (Illumina, San Diego, CA, USA) and scanned by the Fred Hutch Genomics Core facility. Images were converted to expression data using GenomeStudio (Illumina).

**Microarray analysis**

Microarray analysis was done using R. All microarrays were pre-processed within study and sample type using variance stabilizing transformation (Lin et al., 2008) and robust spline normalization from the lumi package (Du et al., 2008). Probes that were rarely expressed in a given study arm were removed.

Differential gene expression was assessed using the limma package (Ritchie et al., 2015), which fits a linear model to each probe measured in the microarray, calculates empirical Bayes moderated t-statistics, and adjusts probe-level p-values for multiple comparisons to control the false discovery rate using the method of Benjamini and Hochberg (Benjamini and Hochberg,

1995). In general, paired models were fit, with modeling done separately for each sample type and study (due to the separate preprocessing of different sample types). For the ACTU-3500 and MTN-017 study, within-participant comparisons were done comparing baseline samples to those obtained at the end of two months of treatment. Similar within-participant comparisons were done for the GMS A study, with the difference that samples during treatment were compared to samples taken two months after the end of treatment. For the GMS B study, treatment samples were compared to samples taken at the same time point from placebo recipients. Individual probes with an adjusted p-value of less than 0.05 were defined as differentially expressed. Gene set testing was done using the camera function from limma,(Wu and Smyth, 2012) using the default inter-gene correlation of 0.01. The camera function is a competitive gene set, meaning that it tests whether the genes in a set are highly ranked as compared to genes outside of the set. The same threshold of 0.05 was used for gene sets, where p-values were adjusted using the false discovery rate for the number of gene sets times the number of study arms tested (e.g. 50 Hallmark gene sets * 11 study arms). Because there are multiple probes for some genes, probes were collapsed into genes by taking the probe with the lowest adjusted p-value for each gene (Ramasamy et al., 2008).

Thirteen cervical samples from the GMS A study clustered separately on principal components analysis plots from the rest of the cervical samples. Differential gene expression analysis comparing this cluster to the rest of the cervical samples revealed tens of thousands of genes to be differentially expressed and suggested that these samples may have included endocervical tissue, rather than only ectocervical as intended, possibly due to cervical ectopy. Keratin genes and gene ontology processes related to keratinocyte and epidermal development were higher in the main group of cervical samples, while the small cluster had higher expression of processes related to cilia movement and development, consistent with the ciliated epithelial cells of the endocervix. Because gene expression differed so dramatically in the thirteen cervical samples in question, we removed them from the analysis.

**Reverse transcription and ddPCR of selected genes**

We used all the paired samples with enough RNA available to repeat measurements of transcript levels for MX dynamin like GTPase 1 (MX1), ISG15 ubiquitin-like modifier (ISG15), and interferon-α inducible protein 6 (IFI6) by ddPCR assay. As the reference gene, we used Ubiquitin C (UBC), selected based on a comparison of commonly used reference genes in the microarray data. Among those genes, UBC was expressed in all sample types and the average fold change (across different sample types and studies) was close to 0 and the standard deviation was small. The samples used for ddPCR were the same as used for microarray, except that the GMS B samples were not used (because they were unpaired), the number of sample pairs was reduced by one each for the vaginal and ectocervical samples from the TDF arm of the GMS A study because of insufficient RNA remaining from those samples. The vaginal samples from the TDF/FTC arm of the GMS A study were not tested by ddPCR because the sample size was so low (only three pairs of samples).

Reverse transcription was performed using 100 ng of RNA per sample in a 20 µL reaction mixture using qScript cDNA Synthesis Kit (QuantaBio, Beverly, MA, USA) according to the manufacturer's instructions. The incubation conditions were 22°C for 5 minutes, 42°C for 30 min, and then 85°C for 5 min. After reverse transcription, the samples were diluted to 100 µL with water and 5 µL (cDNA equivalent of 5 ng RNA) was used per ddPCR well.

The primers and probes used for ddPCR are shown in Supplemental Table 2 and were purchased from Integrated DNA Technologies (Skokie, IL, USA). Assays were run in duplex (IFI6 on the FAM channel with MX1 on the HEX channel in one set of wells and ISG15 on the FAM channel with UBC on the HEX channel in a second set of wells). Each sample was run in duplicate for each assay. Sample pairs (i.e. on- and off-treatment) were always run on the same plates. ddPCR was performed using ddPCR Supermix for Probes (no dUTP), with droplets generated on a QX200 Automated Droplet Generator and droplets read on a QX200 Droplet Reader according to the manufacturer's instructions (Bio-Rad, Hercules, CA, USA).

The ddPCR data was analyzed using QuantaSoft version 1.7.4.0917 (Bio-Rad). The same fluorescence thresholds were applied to all samples across all plates. Wells with fewer than 10,000 droplets were removed. Concentrations of IFI6, MX1 and ISG15 were divided by the concentration of UBC from the corresponding sample to yield copies of each gene per copy of UBC. This value was log2-transformed to convert it to a normal distribution and place it on a comparable scale to the microarray data. Replicate wells were then averaged. Fold changes were calculated by subtracting the expression level from the off-treatment sample from the on-treatment sample.

**RNA sequencing in MTN-017**

Total RNA prepared above was normalized to 300 ng input for library preparation with the TruSeq Stranded Total RNA with Ribo-Zero Globin kit (Illumina). The resulting libraries were assessed on the Agilent Fragment Analyzer with the HS NGS assay (Agilent) and quantified using the KAPA Library Quantification Kit (Roche) on a ViiA 7 Real Time PCR platform (Thermo Fisher). High depth sequencing (50 million reads per sample) was performed with a HiSeq 2500 (Illumina) on two High Output v4 flow cells as a 50 base pair, paired-end run. Raw demultiplexed fastq paired end read files were trimmed of adapters and filtered using the program skewer(Jiang et al., 2014) to remove any reads with an average phred quality score of less than 30 or a length of less than 36 bp. Trimmed reads were aligned using the HISAT2 (Kim et al., 2015) aligner to the Homo sapiens NCBI reference genome assembly version GRCh38 and sorted using SAMtools (Li et al., 2009). Aligned reads were counted and assigned to gene meta-features using the program featureCounts (Liao et al., 2014) as part of the Subread package. Counts data were analyzed analogously to the microarray data, using the voom function from limma and then fitting models for each transcript. Because the samples were processed in two batches, batch number was included in the model in addition to participant ID and treatment.

**Proteomics in MTN-017**

Frozen rectal biopsies from MTN-017 were processed as described previously (Burgener et al., 2013). For protein extraction, tissues were washed 3 times with 10 mM Tris (pH 7.6), placed in 5 mL of a lysis solution consisting of 7 M Urea, 2 M Thiourea, 40 mM Tris, and 10 mM DTT, and homogenized with a gentleMACS Octo Dissociator (RNA02-01M setting, Miltenyi Biotec,

Bergisch Gladbach, Germany). Precipitates were removed by centrifugation at 9000 g for 20 minutes at 4$^{o}$C, transfer of supernatant to a new tube, and a second round of centrifugation at 15,000 g for 20 minutes. Supernatants were stored at -80$^{o}$C. Trypsin digestion was performed as described previously(Birse et al., 2013). Briefly, for each sample, 600 µL of tissue lysate was denatured in urea exchange buffer (8 M Urea in 1:10 0.5 M HEPES:water solution, GE HealthCare, Uppsala, Sweden) and filtered through a 10 kDa membrane. Filtered lysates were alkylated with 50 mM iodoacetamide for 20 minutes, and then washed with 50 mM HEPES buffer. Nucleic acids were removed by treatment with benzonase (150 units/µL in HEPES with $MgCl_2$, Novagen, Darmstadt, Germany) for 30 minutes, and then lysates were washed with HEPES buffer. Trypsin digestion (2 µg trypsin per 100 µg protein, Promega, WI, USA) was performed overnight at 37$^{o}$C. Eluted peptides were dried using a speed vacuum and then stored at -80$^{o}$C. Reverse-phase liquid chromatography using a step-wise gradient was used to remove salts and detergents. Peptide quantification was performed with the LavaPep Fluorescent Protein and Peptide Quantification Kit (Gel Company, San Francisco, CA, USA).

Mass spectrometry was performed using a nano flow liquid chromatography system (Easy nLC, Thermo Fisher) connected inline to a Velos Orbitrap mass spectrometer as described previously (Birse et al., 2015, 2017). One µg of peptide was run for each sample. Feature detection, normalization, and quantification were performed using Progenesis LC-Mass Spectrometry software (Nonlinear Dynamics, Newcastle upon Tyne, UK) with default settings. Peptides were found using Mascot v.2.4.0 (Matrix Science, Boston, MA, USA) to search against the SwissProt database(UniProt Consortium, 2019) restricting taxonomy to Human. Search results were imported into Scaffold (Proteome Software, Portland, OR, USA) for peptide identification, requiring ≤0.1 FDR for protein identification, ≤0.01 FDR for peptide identification, and at least 2 unique peptides identified per protein. Samples were run in two batches, with US participants in one batch and Thai participants in the other. Data from the two cohorts were combined using Combat (Johnson et al., 2007).

**Immunofluorescence microscopy**

Pairs of rectal and duodenal biopsies from eight subjects were stained for ISG15 protein for immunofluorescence microscopy. Duodenal biopsies from two subjects were also stained for glycoprotein 2 to identify M cells (Fabiano et al., 2018). Each pair consisted of one pre- and one on-treatment (~60 days) sample from the ACTU-3500 study. Two rectal biopsies were of poor quality, so they and their pairs were excluded from analysis, reducing the sample size for the rectal biopsies to six pairs. The biopsies were collected into RNAlater Stabilization Solution (ThermoFisher), held at 4 $^{o}$C overnight, and then stored at -80 $^{o}$C. Prior to use, biopsies were thawed, fixed in 10% neutral buffered formalin for 3 days, and stored in 70% ethanol until paraffin embedding. Four micron thick sections were cut, attached to positively-charged slides and baked at 60 $^{o}$C for 1 h, with each slide holding one pre- and one on-treatment tissue section from the same participant. The histopathologist and data analyst were blinded to treatment status. Staining was performed using the procedure described previously (Paulson et al., 2018). Primary antibodies were anti-ISG15 (Atlas Antibodies Cat#HPA004627, RRID: AB_1079152) and anti-glycoprotein 2 (GP2; Thermo Fisher, RRID: AB_2608499).

Slides were scanned on an Aperio FL (Version 2; Leica Biosystems). Exposure times were 125 ms for ISG15 and 64 ms for DAPI. Images were analyzed with HALO v2.2 image analysis software (Indica Labs, Albuquerque, NM, USA) with the CytoNuclear FL v1.4 algorithm. Images were annotated manually to select stroma or epithelium, which were analyzed separately. Individual cells were identified by the software via DAPI-stained nuclei in conjunction with cell-defining parameters including nuclear contrast threshold, minimum nuclear intensity, nuclear segmentation aggressiveness, nuclear size, minimum nuclear roundness and maximum nuclear radius. These parameters were set to be optimal for each pair (i.e. settings were the same for the on- and off-treatment pairs for each person and sample type). The ISG15 signal was so bright that the software identified surrounding cells as positive for ISG15, despite manual inspection clearly showing that only one central cell was positive. To correct for this, hierarchical clustering was used on the spatial positions of the cells identified as positive. A distance threshold was empirically determined to count adjacent cells as a single positive cell, while still identifying nearby but distinct positive cells as distinct. The output of this analysis was verified by comparison to manual counting. Moreover, there was a strong correlation between the percentage of cells that were bright for ISG15 before and after adjustment for falsely positive surrounding cells ($r = 0.98$ for duodenum and 0.95 for rectum), the numbers were simply lower (and more reflective of manual inspection) after adjustment.

**Data analysis**

The following R packages from CRAN or Bioconductor (Huber et al., 2015) were used: AnnotationDbi (Pagès et al., 2018), Biobase (Huber et al., 2015), broom (Robinson, 2016), conflicted (Wickham, 2018), edgeR (Robinson et al., 2010), ggrepel (Slowikowski, 2018), here (Müller, 2017), limma (Ritchie et al., 2015), lumi (Du et al., 2008), msigdbr (Dolgalev, 2018), org.Hs.eg.db (Carlson, 2018), pander (Daróczi and Tsegelskyi, 2018), patchwork (Pedersen, 2017), plater (Hughes, 2016), RColorBrewer (Neuwirth, 2014), tidyverse (Wickham, 2016), and writexl (Ooms, 2018). R was run through RStudio version 1.1.463.
