## Supplementary material for "Treatment with commonly used antiretroviral drugs induces a type I/III interferon signature in the gut in the absence of HIV infection": Code to reproduce analyses and figures: make_paper.R

```
# This file runs all of the scripts that analyze the data and generate the statistics and figures
# for the paper. 

# This script is the only place containing external file paths , so it is the only file that may 
# need manual changes to reproduce the analysis. (For example, after downloading the data from
# GEO, you may need to adjust the file paths to point to the right places.) This script assumes
# that your working directory is the same folder in which it is contained. 

# Load all packages and utility functions
source("R/source_all.R")

###################################################################################################
# Analyze raw data
###################################################################################################
# GMSB study
microarray_GMSB_file <- "data/raw/GSE139411_GMSB_BeadStudio_output.txt.gz"
microarray_GMSB_metadata <- "data/raw/GSE139411_GMSB_sample_metadata.xlsx"
source(here("R/analysis/microarray_GMSB.R"))

# GMSA study
microarray_GMSA_file <- "data/raw/GSE139655_GMSA_BeadStudio_output.txt.gz"
microarray_GMSA_metadata <- "data/raw/GSE139655_GMSA_sample_metadata.xlsx"
source(here("R/analysis/microarray_GMSA.R"))

# MTN-017 study
microarray_mtn017_file <- "data/raw/GSE138723_MTN017_BeadStudio_output.txt.gz"
microarray_mtn017_metadata <- "data/raw/GSE138723_MTN017_sample_metadata.xlsx"
source(here("R/analysis/microarray_MTN017.R"))

# ACTU3500 study
microarray_actu3500_file <- "data/raw/GSE139611_ACTU3500_BeadStudio_output.txt.gz"
microarray_actu3500_metadata <- "data/raw/GSE139611_ACTU3500_sample_metadata.xlsx"
source(here("R/analysis/microarray_ACTU3500.R"))

# Combine microarray data across studies
source(here("R/analysis/microarray_combine_studies.R"))

# ddPCR assays
ddpcr_data_directory <- "data/raw/raw_ddPCR_data"
source(here("R/analysis/ddpcr_comparison.R"))

# microscopy
microscopy_unblinded_log <- "data/raw/microscopy_unblinded_log.csv"
microscopy_isg15_file <- "data/raw/ISG15_microscopy.xlsx"
source(here("R/analysis/microscopy_ACTU3500.R"))

# proteomics
proteomics_mtn017_file <- "data/raw/Proteomics_MTN017.xlsx"
source(here("R/analysis/proteomics_mtn017.R"))

# RNAseq
rnaseq_mtn017_file <- "data/raw/RNAseq_MTN017.xlsx"
source(here("R/analysis/RNAseq_mtn017.R"))

###################################################################################################
# Make figures and statistics
###################################################################################################

# Generate figures
source(here("R/figures.R"))

# Make statistics PDFs
rmarkdown::render(here("R/statistics.Rmd"), output_file = here("R/statistics.pdf"))

source(here("R/generate_supplemental_data.R"))
```
