## Supplementary material for "Treatment with commonly used antiretroviral drugs induces a type I/III interferon signature in the gut in the absence of HIV infection": Code to reproduce analyses and figures: ddpcr_comparison.R

```
quantasoft_directory <- paste0(ddpcr_data_directory, "/ddPCR")
plate_layouts_directory <- paste0(ddpcr_data_directory, "/plate_layouts")
sample_key_file <- paste0(ddpcr_data_directory, "/SampleKey.csv")

# clean plate name and generate other columns
clean_plate <- function(data, files) {
    data %>% 
        mutate(ddPcrPlate = files[as.numeric(ddPcrPlate)], 
            ddPcrPlate = str_extract(ddPcrPlate, "[^\\/]*$"), 
            ddPcrPlate = str_replace(ddPcrPlate, "ddPCR_plate_", ""), 
            ddPcrPlate = str_replace(ddPcrPlate, "[_ubc_second_half]?\\.csv", "")) %>% 
        separate(ddPcrPlate, c("ddPcrPlate", "Fam", "Hex")) %>% 
        mutate(Hex = ifelse(Fam == "isg15", "ubc", "mx1"))
}

# exclude plate 9, the redos
find_files <- function(x) x[str_detect(x, "csv") & !str_detect(x, "9_all")]

# load ddPCR data
files <- list.files(quantasoft_directory, recursive = TRUE, full.names = TRUE) %>% 
    find_files()   

# warning is okay, one file name is longer
data <- map_dfr(.x = files, .f = quiet_read_csv, .id = "ddPcrPlate") %>% 
    clean_plate(files) %>% 
    select(ddPcrPlate:Well, TargetType, Concentration, AcceptedDroplets, Threshold)

# load key
plate_map_files <- list.files(plate_layouts_directory, 
    recursive = TRUE, full.names = TRUE) %>% 
    find_files()

plate_maps <- lapply(plate_map_files, 
    function(x) read_plate(x, well_ids_column = "Well")) %>% 
    data.table::rbindlist(idcol = "ddPcrPlate") %>% 
    clean_plate(plate_map_files) %>% 
    select(-Fam, -Hex) %>% 
    canonicalize_study_names()

key <- quiet_read_csv(sample_key_file) %>% 
    select(Study:Ptid)

# combine 
data <- inner_join(key, plate_maps, by = c("Study", "Index")) %>% 
    inner_join(data, by = c("Well", "ddPcrPlate"))

# clean
data <- data %>% 
    mutate(Target = ifelse(TargetType == "Ch1Unknown", Fam, Hex), 
        Target = str_to_upper(Target)) %>% 
    select(-Fam, -Hex, -TargetType)

make_unique_id <- function(data) {
    data %>% 
        mutate(UniqueId = paste0(Study, Visit, Sample, Index, SpecId, Ptid)) 
}

data <- make_unique_id(data)

redos <- quiet_read_csv(paste0(quantasoft_directory, "/ddPCR_plate_9_all.csv")) %>% 
    mutate(ddPcrPlate = as.character(9)) %>% 
    select(Well, TargetType, Concentration, AcceptedDroplets, Threshold, ddPcrPlate) %>% 
    add_plate(paste0(plate_layouts_directory, "/ddPCR_plate_9_all.csv"), well_ids_column = "Well") %>% 
    mutate(Target = ifelse(TargetType == "Ch1Unknown", 
        str_extract(Assays, "^[^\\/]*"), 
        str_extract(Assays, "[^\\/]*$"))) %>% 
    select(-Assays, -TargetType) %>% 
    mutate(Index = as.character(Index)) %>% 
    canonicalize_study_names()

redos <- inner_join(redos, key, by = c("Study", "Index")) %>% 
    select(Study, Visit, Sample, Index, SpecId, Ptid, ddPcrPlate, Well, 
        Concentration, AcceptedDroplets, Threshold, Target) %>% 
    make_unique_id()

data <- data %>% 
    make_unique_id() %>% 
    filter(!UniqueId %in% redos$UniqueId) %>% 
    bind_rows(redos) %>% 
    select(-UniqueId)

data <- data %>% 
    # remove NTC
    filter(Study != "GMS A" | !Index %in% c(117, 174)) %>% 
    # remove the pairs of the NTCs
    filter(!Index %in% c(183, 170)) %>% 
    # remove failed wells
    filter(AcceptedDroplets >= 10000) %>% 
    # average replicates
    group_by(Study, Visit, Sample, Ptid, ddPcrPlate, Target) %>% 
    summarise(Concentration = mean(Concentration))

# get UBC only data
ref <- data %>% 
    ungroup() %>% 
    filter(Target == "UBC") %>% 
    mutate(Ubc = Concentration) %>% 
    select(-Target, -Concentration, -ddPcrPlate)

# normalize other targets to UBC 
data <- inner_join(data, ref, by = c("Study", "Visit", "Sample", "Ptid")) %>% 
    mutate(NormalizedConcentration = Concentration / Ubc) %>% 
    mutate(Log2NormalizedConcentration = log2(NormalizedConcentration))

data <- data %>% 
    ungroup() %>% 
    mutate(Visit = ifelse(str_detect(Visit, "Baseline|post"), "Off-treatment", "On-treatment"))

ddpcr <- data %>% 
    # all treatments were truvada except for in the GMS A study, where only tenofovir samples were run
    mutate(Treatment = ifelse(Study == "GMS A", "Tenofovir", "Truvada")) %>% 
    canonicalize_treatment_names() %>% 
    canonicalize_sample_names() %>% 
    canonicalize_study_names() %>% 
    canonicalize_visit_names() %>% 
    rename(TargetId = Target)

write_csv(ddpcr, "data/clean/ddpcr.csv")

probes <- get_probe_expression_participant_level() %>% 
    filter(TargetId %in% c("IFI6", "ISG15", "MX1", "UBC")) %>% 
    # one of the UBC probes doesn't have an EntrezID listed
    mutate(EntrezId = ifelse(TargetId == "UBC", 7316, EntrezId))

genes <- probes %>% 
    # use the probes from MTN-017
    filter(ProbeId %in% c(1010246, 2100196, 1690066, 3060300)) %>% 
    group_by(Study, Ptid, Sample, Visit, Treatment, TreatmentLength, TargetId, EntrezId) %>% 
    summarise(Expression = mean(Expression))

microarray_vs_ddpcr <- inner_join(ddpcr, genes, by = c("Study", "Visit", "Sample", "Ptid", "TargetId", "Treatment"))

differences <- microarray_vs_ddpcr %>% 
    group_by(Study, Treatment, Sample, TargetId, Ptid) %>% 
    arrange(Visit) %>% 
    summarise(MicroarrayDiff = Expression[2] - Expression[1], 
        PcrDiff = Log2NormalizedConcentration[2] - Log2NormalizedConcentration[1])

write_csv(differences, "data/clean/microarray_vs_ddpcr.csv")
```
