## Supplementary material for "Treatment with commonly used antiretroviral drugs induces a type I/III interferon signature in the gut in the absence of HIV infection": Code to reproduce analyses and figures: microarray_ACTU3500.R

```
###################################################################################################
# load, add sample key, preprocess, filter
###################################################################################################
actu3500_microarray <- read_lumi(microarray_actu3500_file)

key <- readxl::read_excel(microarray_actu3500_metadata)
key <- as.data.frame(key)
rownames(key) <- key$MicroarrayId
actu3500_microarray <- add_pData(actu3500_microarray, key)

# split out by tissue 
actu3500_pbmc <- actu3500_microarray[ , pData(actu3500_microarray)$Tissue == "PBMC"] %>% 
    pre_process_and_filter(7)

actu3500_rectum <- actu3500_microarray[ , pData(actu3500_microarray)$Tissue == "Rectal"] %>% 
    pre_process_and_filter(7)

actu3500_duodenum <- actu3500_microarray[ , pData(actu3500_microarray)$Tissue == "Duodenal"] %>% 
    pre_process_and_filter(7)

actu3500_wholeblood <- actu3500_microarray[ , pData(actu3500_microarray)$Tissue == "PAXgene"] %>% 
    pre_process_and_filter(7)

save_actu3500_expression_levels <- function(eSet) {
    for_probe_level_key <- transmute(pData(eSet), 
        MicroarrayId = MicroarrayId, Study = "ACTU-3500", Ptid = Ptid, Sample = Tissue, 
        Visit = ifelse(Visit == "Enrollment", "Off treatment", "On treatment"), 
        Treatment = "Truvada", TreatmentLength = "60 days")
    
    make_expression_levels(eSet, for_probe_level_key)
}

actu3500_expression_levels <- map_df(list(actu3500_duodenum, actu3500_pbmc, actu3500_rectum, actu3500_wholeblood), 
    save_actu3500_expression_levels)
save(actu3500_expression_levels, file = "data/clean/participant_level_expression/Microarray_actu3500_expression_levels.Rdata")

###################################################################################################
# modeling
###################################################################################################
fit_model <- function(filtered) {
    visit <- factor(pData(filtered)$Visit)
    ptid <- factor(pData(filtered)$Ptid)
    
    design <- model.matrix(~ 0 + visit + ptid)
    
    fit <- lmFit(filtered, design)
    
    contrasts <- makeContrasts(prep = visitVisit2 - visitEnrollment, levels = design)
    
    fit <- contrasts.fit(fit, contrasts)
    
    fit <- eBayes(fit)
    
    fit
}

n_deg <- function(fit) {
    sum(topTable(fit, coef = "prep", n = Inf)$adj.P.Val < 0.05)
}

pbmc <- fit_model(actu3500_pbmc)
rectum <- fit_model(actu3500_rectum)
duodenum <- fit_model(actu3500_duodenum)
wholeblood <- fit_model(actu3500_wholeblood)

save_top_table(pbmc, "prep", "ACTU3500_Truvada_Oral_60Days_Daily_PBMC", "Microarray")
save_top_table(rectum, "prep", "ACTU3500_Truvada_Oral_60Days_Daily_15cmRectum", "Microarray")
save_top_table(duodenum, "prep", "ACTU3500_Truvada_Oral_60Days_Daily_Duodenum", "Microarray")
save_top_table(wholeblood, "prep", "ACTU3500_Truvada_Oral_60Days_Daily_PAXgene", "Microarray")

###################################################################################################
# Camera gene set testing
###################################################################################################
hallmark <- get_hallmark_gene_sets_for_camera()

do_actu3500_camera <- function(eSet, fit, cond, gene_sets, gene_sets_name, assay) {
    identifiers <- as.character(fData(eSet)[ , "ENTREZ_GENE_ID"])
    index <- ids2indices(gene.sets = gene_sets, identifiers)
    
    result <- limma::cameraPR(fit$t[,"prep"], index)
    save_camera(result, gene_sets_name, cond, assay)  
}  

do_actu3500_camera(actu3500_pbmc, pbmc, "ACTU3500_Truvada_Oral_60Days_Daily_PBMC", hallmark, "Hallmark", "Microarray")
do_actu3500_camera(actu3500_rectum, rectum, "ACTU3500_Truvada_Oral_60Days_Daily_15cmRectum", hallmark, "Hallmark", "Microarray")
do_actu3500_camera(actu3500_duodenum, duodenum, "ACTU3500_Truvada_Oral_60Days_Daily_Duodenum", hallmark, "Hallmark", "Microarray")
do_actu3500_camera(actu3500_wholeblood, wholeblood, "ACTU3500_Truvada_Oral_60Days_Daily_PAXgene", hallmark, "Hallmark", "Microarray")

load("data/clean/oral_gene_sets_for_camera.Rda")
do_actu3500_camera(actu3500_pbmc, pbmc, "ACTU3500_Truvada_Oral_60Days_Daily_PBMC", oral_gene_sets_for_camera, "OralDEGs", "Microarray")
do_actu3500_camera(actu3500_rectum, rectum, "ACTU3500_Truvada_Oral_60Days_Daily_15cmRectum", oral_gene_sets_for_camera, "OralDEGs", "Microarray")
do_actu3500_camera(actu3500_duodenum, duodenum, "ACTU3500_Truvada_Oral_60Days_Daily_Duodenum", oral_gene_sets_for_camera, "OralDEGs", "Microarray")
do_actu3500_camera(actu3500_wholeblood, wholeblood, "ACTU3500_Truvada_Oral_60Days_Daily_PAXgene", oral_gene_sets_for_camera, "OralDEGs", "Microarray")
```
