## Supplementary material for "Treatment with commonly used antiretroviral drugs induces a type I/III interferon signature in the gut in the absence of HIV infection": Code to reproduce analyses and figures: microarray_combine_studies.R

```
###################################################################################################
# Top tables
###################################################################################################
# Combine the top tables from all the different studies into a single file
read_top_table <- function(f) {
    data <- read_csv(f, progress = FALSE, col_types = cols(
        ProbeID = col_double(),
        TargetID = col_character(),
        ILMN_GENE = col_character(),
        ENTREZ_GENE_ID = col_double(),
        GI = col_double(),
        ACCESSION = col_character(),
        SYMBOL = col_character(),
        PROBE_ID = col_character(),
        PROBE_START = col_double(),
        PROBE_SEQUENCE = col_character(),
        CHROMOSOME = col_character(),
        PROBE_CHR_ORIENTATION = col_character(),
        PROBE_COORDINATES = col_character(),
        logFC = col_double(),
        AveExpr = col_double(),
        t = col_double(),
        P.Value = col_double(),
        adj.P.Val = col_double(),
        B = col_double()
    )) %>% 
            .[ , c("ProbeID", "TargetID", "ENTREZ_GENE_ID", "logFC", "P.Value", "adj.P.Val", "t")]
    
    # add name of condition
    data$Condition <- f %>% str_remove(".*Microarray_") %>% 
        str_remove(".csv")
    
    data
}

files <- list.files("data/clean/top_tables/", full.names = TRUE) %>% 
    .[str_detect(., "Microarray_")]

all <- map_df(files, read_top_table)

all <- tidyr::separate(all, Condition, 
    into = c("Study", "Treatment", "TreatmentLocation", "TreatmentLength",
        "TreatmentInterval", "Sample"), sep = "_")

# Clean up descriptions of conditions/genes and column names
probe_fold_changes <- all %>% 
    canonicalize_study_names() %>% 
    canonicalize_sample_names() %>% 
    canonicalize_treatment_names() %>% 
    # TreatmentLocation and TreatmentInterval are all the same for these samples
    # (oral and daily)
    select(-TreatmentLocation, -TreatmentInterval) %>% 
    rename(ProbeId = ProbeID, TargetId = TargetID, EntrezId = ENTREZ_GENE_ID, 
        Log2FoldChange = logFC, PValue = P.Value, AdjustedPValue = adj.P.Val) %>% 
    canonicalize_gene_symbols()

write_csv(probe_fold_changes, path = "data/clean/probe_fold_changes.csv")

# fold change data at gene level ----------------------------------------------------

# count differentially expressed genes 
# group into unique study arms (Study, Treatment, TreatmentLength, Sample)
# and into unique genes (TargetId, EntrezId)
# and into whether that probe is significant or not
# So for each target ID within each study arm, you have a group of probes that were significant and a group that weren't
# - If multiple were significant in different directions --> don't count (these genes are dropped)
# - Otherwise, take the probe with the lowest p-value
gene_fold_changes <- probe_fold_changes %>% 
    group_by(Study, Treatment, TreatmentLength, Sample, TargetId, EntrezId, Sig = AdjustedPValue < 0.05) %>% 
    # remove cases where multiple probes are significant in different directions 
    mutate(Direction = ifelse(all(Log2FoldChange < 0) | all(Log2FoldChange > 0) | !Sig, "okay", "problem")) %>% 
    filter(Direction != "problem") %>% 
    # take the probe with the lowest p-value for each gene
    group_by(Study, Treatment, TreatmentLength, Sample, TargetId, EntrezId) %>% 
    arrange(PValue) %>% 
    slice(1) %>% 
    select(Study, Treatment, TreatmentLength, Sample, TargetId, EntrezId, Log2FoldChange, PValue, AdjustedPValue)

write_csv(gene_fold_changes, path = "data/clean/gene_fold_changes.csv")

###################################################################################################
# Hallmark gene sets
###################################################################################################
files <- list.files("data/clean/camera", full.names = TRUE) %>% 
    .[str_detect(., "Microarray_Hallmark")]

hallmarks <- map_df(files, read_csv) %>% 
    select(-X1)

# correctly calculate FDR to adjust for all tests across all studies
hallmarks$FDR <- p.adjust(hallmarks$PValue, method = "fdr")

# tidy up metadata
hallmarks <- hallmarks %>% 
    #remove "HALLMARK_" from the gene set names
    mutate(Gene_Set = str_replace(Gene_Set, "HALLMARK_", "")) %>%
    # create individual columns from Data_Tested
    tidyr::separate(col = Data_Tested, sep = "_", into = 
            c("Study", "Treatment", "TreatmentLength", "TreatmentInterval", 
                "TreatmentLocation", "Sample" )) %>% 
    select(-TreatmentLocation, -TreatmentInterval) %>% 
    canonicalize_treatment_names() %>% 
    canonicalize_study_names() %>% 
    canonicalize_sample_names() %>% 
    # create Data_Tested with only relevant information
    mutate(Data_Tested = paste(Sample, Study, sep = "\n"))

# create column indicating how many study arms are significant for each gene set
hallmarks <- hallmarks %>% 
    group_by(Gene_Set) %>%
    mutate(Sig = sum(FDR < 0.05)) %>% 
    arrange(Sig)

write_csv(hallmarks, "data/clean/hallmark_camera.csv")    

###################################################################################################
# DEG gene sets
###################################################################################################
files <- list.files("data/clean/camera", full.names = TRUE) %>% 
    .[str_detect(., "Microarray_OralDEGs")]

oral_only <- map_df(files, read_csv) %>% 
    select(-X1)

oral_only <- oral_only %>% 
    # create individual columns from Data_Tested
    tidyr::separate(col = Data_Tested, sep = "_", into = 
            c("Study", "Treatment", "TreatmentLength", "TreatmentInterval", 
                "TreatmentLocation", "Sample" )) %>% 
    select(-TreatmentLocation, -TreatmentInterval) %>% 
    canonicalize_treatment_names() %>% 
    canonicalize_study_names() %>% 
    canonicalize_sample_names() %>% 
    # create Data_Tested with only relevant information
    mutate(Data_Tested = paste(Sample, Study, sep = "\n"))

oral_only <- oral_only %>% 
    # remove weird combined gene set
    filter(!(str_detect(Gene_Set, "MTN017/PrEP"))) %>% 
    # remove cases where gene set was tested against its founder set    
    filter(!(Study == "ACTU-3500" & Sample == "Duodenum" & str_detect(Gene_Set, "PrEP-Truvada-60Days-Daily-Oral-Duodenum"))) %>% 
    filter(!(Study == "MTN-017" & Gene_Set == "UP-MTN017-Truvada-56Days-Daily-Oral-15cmRectum"))

# correctly calculate FDR to adjust for all tests across all studies
oral_only$FDR <- p.adjust(oral_only$PValue, method = "fdr")

# better gene set names
oral_only <- oral_only %>% 
    mutate(Gene_Set = fct_recode(Gene_Set, 
        "DOWN\nDuodenum\nACTU-3500" = "DOWN-PrEP-Truvada-60Days-Daily-Oral-Duodenum", 
        "UP\nDuodenum\nACTU-3500" = "UP-PrEP-Truvada-60Days-Daily-Oral-Duodenum", 
        "UP\nRectum\nMTN-017" = "UP-MTN017-Truvada-56Days-Daily-Oral-15cmRectum"))

write_csv(oral_only, "data/clean/deg_sets_camera.csv")

###################################################################################################
# Probe expression data at participant level
###################################################################################################
files <- list.files("data/clean/participant_level_expression", full.names = TRUE) %>% 
    .[str_detect(., "Microarray")]

for (file in files) load(file)

mtn017_expression_levels$Ptid <- as.character(mtn017_expression_levels$Ptid)
GMSA_expression_levels$Ptid <- as.character(GMSA_expression_levels$Ptid)
GMSA_expression_levels$MicroarrayId <- as.character(GMSA_expression_levels$MicroarrayId)
expression <- bind_rows(GMSA_expression_levels, actu3500_expression_levels, mtn017_expression_levels)

expression <- expression %>% 
    canonicalize_treatment_names() %>% 
    canonicalize_sample_names() %>% 
    canonicalize_study_names() %>% 
    canonicalize_visit_names() %>%
    # make column names consistent
    rename(TargetId = Gene)

saveRDS(expression, file = "data/clean/probe_expression.Rdata")
```
