## Supplementary material for "Treatment with commonly used antiretroviral drugs induces a type I/III interferon signature in the gut in the absence of HIV infection": Code to reproduce analyses and figures: microarray_GMSA.R

```
###################################################################################################
# load, add sample key, preprocess, filter
###################################################################################################
GMSA_microarray <- read_lumi(microarray_GMSA_file)

key <- readxl::read_excel(microarray_GMSA_metadata)
key <- as.data.frame(key)
rownames(key) <- key$MicroarrayId
GMSA_microarray <- add_pData(GMSA_microarray, key)

GMSA_microarray <- GMSA_microarray[ , pData(GMSA_microarray)$Paired == "Paired"]

# split out by tissue (ignore endocervical because too few paired)
GMSA_ectocervix <- GMSA_microarray[ , pData(GMSA_microarray)$Tissue == "Ectocervix"] %>% 
    pre_process_and_filter(8)

GMSA_vagina <- GMSA_microarray[ , pData(GMSA_microarray)$Tissue == "Vagina"] %>% 
    pre_process_and_filter(8)

GMSA_pbmc <- GMSA_microarray[ , pData(GMSA_microarray)$Tissue == "PBMC"] %>% 
    pre_process_and_filter(8)

save_GMSA_expression_levels <- function(eSet) {
    for_probe_level_key <- transmute(pData(eSet), 
        MicroarrayId = MicroarrayId, Study = "GMS A", Ptid = Ptid, 
        Sample = Tissue,
        Visit = ifelse(Prep == "post.prep", "Off treatment", "On treatment"),
        Treatment = Drug, TreatmentLength = "60 days")
    
    make_expression_levels(eSet, for_probe_level_key)
}

GMSA_expression_levels <- map_df(list(GMSA_ectocervix, GMSA_vagina, GMSA_pbmc), 
    save_GMSA_expression_levels)
save(GMSA_expression_levels, file = "data/clean/participant_level_expression/Microarray_GMSA_expression_levels.Rdata")

###################################################################################################
# modeling
###################################################################################################
fit_model <- function(filtered, drug) {
    indices <- pData(filtered) %>% 
        filter(Prep %in% c("prep", "post.prep")) %>% 
        group_by(Ptid) %>% 
        filter(n() > 1) %>% 
        .$MicroarrayId
    filtered <- filtered[ , colnames(filtered) %in% indices]
    filtered <- filtered[ , pData(filtered)$Drug == drug]
    
    prep <- factor(pData(filtered)$Prep)
    ptid <- factor(pData(filtered)$Ptid)
    
    design <- model.matrix(~ 0 + prep + ptid)
    
    fit <- lmFit(filtered, design)
    
    contrasts <- makeContrasts(prep = prepprep - preppost.prep, levels = design)
    
    fit <- contrasts.fit(fit, contrasts)
    
    fit <- eBayes(fit)
    
    fit
}

n_deg <- function(fit) {
    sum(topTable(fit, coef = "prep", n = Inf)$adj.P.Val < 0.05)
}

ecto_tenofovir <- fit_model(GMSA_ectocervix, "TDF")
pbmc_tenofovir <- fit_model(GMSA_pbmc, "TDF")
vag_truvada <- fit_model(GMSA_vagina, "FTC/TDF")
vag_tenofovir <- fit_model(GMSA_vagina, "TDF")

save_top_table(ecto_tenofovir, "prep", "GMSA_Tenofovir_Oral_55Days_Daily_Ectocervix", "Microarray")
save_top_table(pbmc_tenofovir, "prep", "GMSA_Tenofovir_Oral_55Days_Daily_PBMC", "Microarray")
save_top_table(vag_tenofovir, "prep", "GMSA_Tenofovir_Oral_55Days_Daily_Vagina", "Microarray")
save_top_table(vag_truvada, "prep", "GMSA_Truvada_Oral_55Days_Daily_Vagina", "Microarray")

###################################################################################################
# Camera gene set testing
###################################################################################################
hallmark <- get_hallmark_gene_sets_for_camera()

do_GMSA_camera <- function(eSet, fit, cond, gene_sets, gene_sets_name, assay) {
    identifiers <- as.character(fData(eSet)[ , "ENTREZ_GENE_ID"])
    index <- ids2indices(gene.sets = gene_sets, identifiers)
    
    result <- limma::cameraPR(fit$t[,"prep"], index)
    save_camera(result, gene_sets_name, cond, assay)  
}  

do_GMSA_camera(GMSA_ectocervix, ecto_tenofovir, "GMSA_Tenofovir_Oral_55Days_Daily_Ectocervix", hallmark, "Hallmark", "Microarray")
do_GMSA_camera(GMSA_pbmc, pbmc_tenofovir, "GMSA_Tenofovir_Oral_55Days_Daily_PBMC", hallmark, "Hallmark", "Microarray")
do_GMSA_camera(GMSA_vagina, vag_tenofovir, "GMSA_Tenofovir_Oral_55Days_Daily_Vagina", hallmark, "Hallmark", "Microarray")
do_GMSA_camera(GMSA_vagina, vag_truvada, "GMSA_Truvada_Oral_55Days_Daily_Vagina", hallmark, "Hallmark", "Microarray")

load("data/clean/oral_gene_sets_for_camera.Rda")
do_GMSA_camera(GMSA_ectocervix, ecto_tenofovir, "GMSA_Tenofovir_Oral_55Days_Daily_Ectocervix", oral_gene_sets_for_camera, "OralDEGs", "Microarray")
do_GMSA_camera(GMSA_pbmc, pbmc_tenofovir, "GMSA_Tenofovir_Oral_55Days_Daily_PBMC", oral_gene_sets_for_camera, "OralDEGs", "Microarray")
do_GMSA_camera(GMSA_vagina, vag_tenofovir, "GMSA_Tenofovir_Oral_55Days_Daily_Vagina", oral_gene_sets_for_camera, "OralDEGs", "Microarray")
do_GMSA_camera(GMSA_vagina, vag_truvada, "GMSA_Truvada_Oral_55Days_Daily_Vagina", oral_gene_sets_for_camera, "OralDEGs", "Microarray")
```
