## Supplementary material for "Treatment with commonly used antiretroviral drugs induces a type I/III interferon signature in the gut in the absence of HIV infection": Code to reproduce analyses and figures: microarray_GMSB.R

```
###################################################################################################
# load, add sample key, preprocess, filter
###################################################################################################
GMSB_microarray <- read_lumi(microarray_GMSB_file)

key <- readxl::read_excel(microarray_GMSB_metadata)
key <- as.data.frame(key)
rownames(key) <- key$MicroarrayId
GMSB_microarray <- add_pData(GMSB_microarray, key)

GMSB_microarray <- pre_process_and_filter(GMSB_microarray, 10)

for_prep <- GMSB_microarray[ , pData(GMSB_microarray)$Inclusion == "Included"]

###################################################################################################
# modeling
###################################################################################################
drug <- as.character(pData(for_prep)$Drug)
drug <- factor(ifelse(drug == "FTC/TDF", "Truvada", drug))

design <- model.matrix(~ 0 + drug)

fit <- lmFit(for_prep, design)

contrasts <- makeContrasts(tdf = drugTDF - drugPlacebo, truvada = drugTruvada - drugPlacebo, 
    levels = design)

fit <- contrasts.fit(fit, contrasts)

fit <- eBayes(fit)

save_top_table(fit, "tdf", "GMSB_Tenofovir_Oral_NA_Daily_PBMC", "Microarray")
save_top_table(fit, "truvada", "GMSB_Truvada_Oral_NA_Daily_PBMC", "Microarray")

###################################################################################################
# Camera gene set testing
###################################################################################################
hallmark <- get_hallmark_gene_sets_for_camera()

do_GMSB_camera <- function(drug, cond, gene_sets, gene_sets_name, assay) {
    identifiers <- as.character(fData(for_prep)[ , "ENTREZ_GENE_ID"])
    index <- ids2indices(gene.sets = gene_sets, identifiers)
    result <- limma::camera(y = exprs(for_prep), index = index, 
        design = design, contrast = contrasts[ , drug, drop = FALSE])
    
    save_camera(result, gene_sets_name, cond, assay)  
}  

do_GMSB_camera("tdf", "GMSB_Tenofovir_NA_Daily_Oral_PBMC", hallmark, "Hallmark", "Microarray")
do_GMSB_camera("truvada", "GMSB_Truvada_NA_Daily_Oral_PBMC", hallmark, "Hallmark", "Microarray")

load("data/clean/oral_gene_sets_for_camera.Rda")
do_GMSB_camera("tdf", "GMSB_Tenofovir_NA_Daily_Oral_PBMC", oral_gene_sets_for_camera, "OralDEGs", "Microarray")
do_GMSB_camera("truvada", "GMSB_Truvada_NA_Daily_Oral_PBMC", oral_gene_sets_for_camera, "OralDEGs", "Microarray")
```
