## Supplementary material for "Treatment with commonly used antiretroviral drugs induces a type I/III interferon signature in the gut in the absence of HIV infection": Code to reproduce analyses and figures: microarray_MTN017.R

```
###################################################################################################
# load, add sample key, preprocess, filter
###################################################################################################
mtn017_microarray <- read_lumi(microarray_mtn017_file)

key <- readxl::read_excel(microarray_mtn017_metadata)
key <- key %>% 
    separate(SampleTitle, into = c("Study", "Ptid", "Sample", "Drug", "Treatment", "Visit"), sep = "_") %>% 
    mutate(Ptid = paste(Study, Ptid, sep = "_"))
key <- as.data.frame(key)
rownames(key) <- key$MicroarrayId
mtn017_microarray <- add_pData(mtn017_microarray, key)

mtn017_microarray <- pre_process_and_filter(mtn017_microarray, 10)

for_probe_level <- mtn017_microarray[ , pData(mtn017_microarray)$Treatment %in% c("Baseline", "Oral")]

for_probe_level_key <- transmute(pData(for_probe_level), 
    MicroarrayId = MicroarrayId, Study = "MTN-017", Ptid = Ptid, Sample = "15cmRectum", 
    Visit = ifelse(Treatment == "Baseline", "Off treatment", "On treatment"), 
    Treatment = "Truvada", TreatmentLength = "60 days")

mtn017_expression_levels <- make_expression_levels(for_probe_level, for_probe_level_key)
save(mtn017_expression_levels, file = "data/clean/participant_level_expression/Microarray_mtn017_expression_levels.Rdata")

###################################################################################################
# modeling
###################################################################################################
trt_ <- pData(mtn017_microarray)$Treatment %>% factor()
ptid <- factor(pData(mtn017_microarray)$Ptid)
design <- model.matrix(~ 0 + trt_ + ptid)

fit <- lmFit(mtn017_microarray, design)
contrasts <- makeContrasts(prep = trt_Oral - trt_Baseline,
    levels = design)
fit <- contrasts.fit(contrasts, fit = fit)
fit <- eBayes(fit)

save_top_table(fit, "prep", "MTN017_Truvada_Oral_60Days_Daily_Rectum", "Microarray")

###################################################################################################
# Camera gene set testing
###################################################################################################
hallmark <- get_hallmark_gene_sets_for_camera()

do_mtn017_camera <- function(eSet, fit, cond, gene_sets, gene_sets_name, assay) {
    identifiers <- as.character(fData(eSet)[ , "ENTREZ_GENE_ID"])
    index <- ids2indices(gene.sets = gene_sets, identifiers)
    
    result <- limma::cameraPR(fit$t[,"prep"], index)
    save_camera(result, gene_sets_name, cond, assay)  
}  

do_mtn017_camera(mtn017_microarray, fit, "MTN017_Truvada_Oral_60Days_Daily_Rectum", hallmark, "Hallmark", "Microarray")

load("data/clean/oral_gene_sets_for_camera.Rda")
do_mtn017_camera(mtn017_microarray, fit, "MTN017_Truvada_Oral_60Days_Daily_Rectum", oral_gene_sets_for_camera, "OralDEGs", "Microarray")
```
