## Supplementary material for "Treatment with commonly used antiretroviral drugs induces a type I/III interferon signature in the gut in the absence of HIV infection": Code to reproduce analyses and figures: microscopy_ACTU3500.R

```
###############################################################################
# ISG15 microscopy
###############################################################################
bad <- quiet_read_csv(microscopy_unblinded_log) %>%
    filter(Notes == "Bad Sample") %>% .$Ptid

# a couple of rectal slides were bad, have to remove all rectal samples from those donors
remove_bads <- function(d) {
    d %>% 
        filter(Sample == "Duodenum" | !Ptid %in% bad)
}

isg15 <- readxl::read_excel(microscopy_isg15_file, sheet = "ISG15_raw_data") %>% 
    mutate(`Image.Location ISG15` = str_extract(`Image Location`, "[^\\\\]*.afi$")) %>% 
    left_join(quiet_read_csv(microscopy_unblinded_log), by = "Image.Location ISG15") %>% 
    remove_bads() %>% 
    mutate(Isg15Intensity = `Dye 1 Cytoplasm Intensity`, 
        Isg15Classification = `Dye 1 Classification`, 
        SlideId = `Image Location`) %>% 
    select(Ptid, Visit, Sample, SlideId, Isg15Intensity, Isg15Classification, XMin, XMax, YMin, YMax) %>% 
    canonicalize_sample_names() %>% 
    canonicalize_visit_names() 

write_csv(isg15, "data/clean/microscopy_isg15_raw.csv")

summarized_isg15 <- isg15 %>% 
    mutate(Visit = str_replace(Visit, " ", "\n")) %>% 
    mutate(Class = ifelse(Isg15Classification < 2, "Dim/Negative", "Bright")) %>% 
    group_by(Ptid, Visit, Class, Sample) %>% 
    summarise(n = n(), Intensity = median(Isg15Intensity)) %>% 
    group_by(Ptid, Visit, Sample) %>% 
    mutate(Percent = n / sum(n) * 100)

write_csv(summarized_isg15, "data/clean/microscopy_isg15_summarized.csv")

# collapse cells that are bright for ISG15 down to a center one, don't --------
# include adjacent ones -------------------------------------------------------
brights <- isg15 %>% 
    filter(Isg15Classification >= 2)

others <- isg15 %>% 
    filter(Isg15Classification < 2)

do_clustering <- function(d) {
    clustered <- d %>% 
        select(XMin, XMax, YMin, YMax) %>% 
        dist() %>% 
        hclust()
    
    d %>% 
        mutate(Clust = cutree(clustered, h = 75))
}

clustered <- brights %>% 
    group_by(Ptid, Visit, Sample, SlideId) %>% 
    do(do_clustering(.)) %>% 
    bind_rows(others) %>%     
    group_by(Ptid, Visit, Sample) %>% 
    summarise(Isg15 = max(Clust, na.rm = TRUE), Total = n(), Percent = Isg15 / Total * 100) 

write_csv(clustered, "data/clean/microscopy_isg15_clustered_brights.csv")
```
