## Supplementary material for "Treatment with commonly used antiretroviral drugs induces a type I/III interferon signature in the gut in the absence of HIV infection": Code to reproduce analyses and figures: proteomics_MTN017.R

```
###################################################################################################
# load data
###################################################################################################
proteomics <- readxl::read_excel(proteomics_mtn017_file, sheet = "raw_data") 
lookups <- readxl::read_excel(proteomics_mtn017_file, sheet = "protein_lookup")

###################################################################################################
# prepare data for limma
###################################################################################################
proteomics <- proteomics %>% 
    mutate(ID = paste(Description, Ptid, Visit, Treatment, PtidTreatment, sep = "_")) %>% 
    select(ID, everything())

data_matrix <- proteomics %>% 
    select(-(ID:PtidTreatment)) %>% 
    t() %>% 
    log2()
colnames(data_matrix) <- proteomics$ID

pData <- proteomics %>% 
    select(ID:PtidTreatment) %>% 
    as.data.frame()
rownames(pData) <- pData$ID

fData <- tibble(`UniProt Accession ID` = rownames(data_matrix)) %>% 
    inner_join(lookups, by = "UniProt Accession ID") %>% 
    rename(UniProtAccessionId = "UniProt Accession ID", EntrezId = `Gene ID`, TargetId = `Gene Symbol`) %>% 
    canonicalize_gene_symbols() %>% 
    as.data.frame() %>% 
    .[order(match(.$UniProtAccessionId, rownames(data_matrix))), ]
rownames(fData) <- fData$UniProtAccessionId

thai_proteomics <- Biobase::ExpressionSet(data_matrix, phenoData = Biobase::AnnotatedDataFrame(pData),
    featureData = Biobase::AnnotatedDataFrame(fData))
save(thai_proteomics, file = "data/clean/proteomics_thai.Rda")

###################################################################################################
# save participant level expression levels
###################################################################################################
key <- pData(thai_proteomics) %>% 
    transmute(SampleId = ID, Study = "MTN-017", Ptid = Ptid, Sample = "15cmRectum",
        Visit = ifelse(Treatment == "Baseline", "Off treatment", "On treatment"),
        Treatment = "Truvada", TreatmentLength = "60 days") %>% 
    canonicalize_sample_names() %>% 
    canonicalize_study_names() %>% 
    canonicalize_treatment_names() %>% 
    canonicalize_visit_names()

data <- reshape2::melt(exprs(thai_proteomics))
colnames(data) <- c("UniProtAccessionId", "SampleId", "Expression")

data <- merge(data, key, by = "SampleId")

features <- thai_proteomics %>%
    fData() %>% 
    canonicalize_gene_symbols() %>% 
    select(UniProtAccessionId, TargetId, EntrezId)

proteomics_mtn017_expression_levels <- merge(data, features, by = "UniProtAccessionId")

saveRDS(proteomics_mtn017_expression_levels, file = "data/clean/participant_level_expression/Proteomics_mtn017_expression_levels.Rdata")

###################################################################################################
# linear modeling
###################################################################################################
ptid <- factor(pData(thai_proteomics)$Ptid)
treatment <- factor(pData(thai_proteomics)$Treatment)
design <- model.matrix(~ 0 + treatment + ptid)
model <- lmFit(thai_proteomics, design)

contrasts <- makeContrasts(Drug = treatmentOral - treatmentBaseline, levels = design)
model <- contrasts.fit(model, contrasts = contrasts)
model <- eBayes(model)
tt <- topTable(model, coef = "Drug", n = Inf) %>% 
    rename(AdjustedPValue = adj.P.Val, PValue = P.Value, Log2FoldChange = logFC)

write_csv(tt, "data/clean/proteomics_thai_protein_fold_changes.csv")
```
