## Supplementary material for "Treatment with commonly used antiretroviral drugs induces a type I/III interferon signature in the gut in the absence of HIV infection": Code to reproduce analyses and figures: rnaseq_mtn017.R

```
###################################################################################################
# load, add sample key and gene annotation
###################################################################################################
# exprs -------------------------------------------------------------------------------------------
x <- readxl::read_excel(rnaseq_mtn017_file, sheet = "count_matrix")
counts <- as.matrix(x[,2:ncol(x)])
rownames(counts) <- x$Gene
class(counts) <- "numeric"

# pData -------------------------------------------------------------------------------------------
pData <- readxl::read_excel(rnaseq_mtn017_file, sheet = "sample_metadata") %>% as.data.frame()

# put pData in the order that eSet is in
rownames(pData) <- pData$sample.id
all(nrow(pData) == ncol(counts))
pData <- pData[order(match(rownames(pData), colnames(counts))), ]
all(colnames(counts) == rownames(pData))

# fData -------------------------------------------------------------------------------------------
featureData <- readxl::read_excel(rnaseq_mtn017_file, sheet = "transcript_metadata")

lookup <- AnnotationDbi::select(org.Hs.eg.db::org.Hs.eg.db,
        keys = as.character(unique(featureData$TargetId)),
        columns = c("ENTREZID", "SYMBOL"), keytype = "SYMBOL") %>% 
    # some symbols have duplicate EntrezIds, only a few, remove
    group_by(SYMBOL) %>% 
    filter(n() == 1) %>% 
    rename(TargetId = SYMBOL, EntrezId = ENTREZID) %>% 
    ungroup()

featureData <- left_join(featureData, lookup, by = c("TargetId", "EntrezId")) %>% 
    as.data.frame()

# put fData in the order that eSet is in
rownames(featureData) <- featureData$GeneId
nrow(featureData) == nrow(counts)
featureData <- featureData[order(match(rownames(featureData), rownames(counts))), ]
all(rownames(counts) == rownames(featureData))

x2 <- DGEList(counts = counts, samples = pData, genes = featureData)

###################################################################################################
# preprocess, filter
###################################################################################################
trt_ <- x2$samples$TreatmentCode %>% str_replace_all(" ", "_") %>% factor()
ptid <- factor(x2$samples$donor.id)
batch <- factor(x2$samples$batch)

# hist(cpm(x2, log = TRUE), main = "prefilter")
# # library size similar across samples
# plot(x2@.Data[[2]]$lib.size)
# plot(prcomp(t(counts))$x[,1:2], col = factor(trt_))
# # 
# # # PC1 directly proportional to library size
# plot(prcomp(t(counts))$x[,1] ~ x2@.Data[[2]]$lib.size)

design <- model.matrix(~ 0 + trt_ + ptid + batch)

keep <- edgeR::filterByExpr(x2, design)
x2 <- x2[keep, , keep.lib.sizes = FALSE]

# plot(prcomp(t(x2$counts))$x[,1] ~ x2@.Data[[2]]$lib.size)
# plot(prcomp(t(x2$counts))$x[,1:2], col = factor(trt_))
# 
# 
# hist(cpm(x2, log = TRUE), main = "postfilter")

x2 <- calcNormFactors(x2)
x2 <- voom(x2, design, plot = FALSE)

###################################################################################################
# save participant level expression levels
###################################################################################################
for_probe_level <- x2[ , x2$targets$TreatmentCode %in% c("Baseline", "Oral")]

for_probe_level_key <- transmute(for_probe_level$targets,
    SampleId = sample.id, Study = "MTN-017", Ptid = donor.id, Sample = "15cmRectum",
    Visit = ifelse(TreatmentCode == "Baseline", "Off treatment", "On treatment"),
    Treatment = "Truvada", TreatmentLength = "60 days") %>% 
    canonicalize_sample_names() %>% 
    canonicalize_study_names() %>% 
    canonicalize_treatment_names() %>% 
    canonicalize_visit_names()

data <- reshape2::melt(for_probe_level$E)
colnames(data) <- c("GeneId", "SampleId", "Expression")

data <- merge(data, for_probe_level_key, by = "SampleId")

features <- for_probe_level$genes %>% 
    select(TargetId, EntrezId, GeneId)

rnaseq_mtn017_expression_levels <- merge(data, features, by = "GeneId")

saveRDS(rnaseq_mtn017_expression_levels, file = "data/clean/participant_level_expression/RNAseq_mtn017_expression_levels.Rdata")

###################################################################################################
# modeling
###################################################################################################
fit <- lmFit(x2, design)
contrasts <- makeContrasts(prep = trt_Oral - trt_Baseline,
    levels = design, trend = TRUE)
fit <- contrasts.fit(contrasts, fit = fit)
fit <- eBayes(fit)

tt <- topTable(fit, coef = "prep", n = Inf) %>% 
    rename(AdjustedPValue = adj.P.Val, PValue = P.Value, Log2FoldChange = logFC)

write_csv(tt, "data/clean/rnaseq_gene_fold_changes.csv")

###################################################################################################
# Camera gene set testing
###################################################################################################
hallmark <- get_hallmark_gene_sets_for_camera()

do_mtn017_camera <- function(eSet, fit, cond, gene_sets, gene_sets_name, assay) {
    identifiers <- as.character(x2$genes[ , "EntrezId"])
    index <- ids2indices(gene.sets = gene_sets, identifiers)
    
    result <- limma::cameraPR(fit$t[,"prep"], index)
    save_camera(result, gene_sets_name, cond, assay)  
}  

do_mtn017_camera(mtn017_microarray, fit, "MTN017_Truvada_Oral_60Days_Daily_Rectum", hallmark, "Hallmark", "RNAseq")

load("data/clean/oral_gene_sets_for_camera.Rda")
do_mtn017_camera(mtn017_microarray, fit, "MTN017_Truvada_Oral_60Days_Daily_Rectum", oral_gene_sets_for_camera, "OralDEGs", "RNAseq")
```
