## Supplementary material for "Treatment with commonly used antiretroviral drugs induces a type I/III interferon signature in the gut in the absence of HIV infection": Code to reproduce analyses and figures: figure_helpers.R

```
fdr_neg_log10 <- "FDR-adjusted p-value (-log10)"

# helper functions and constants for ordering the facets of plots separately by tfv and truv
truvada_levels <- c("Duodenum\nACTU-3500", "Rectum\nACTU-3500", "Rectum\nMTN-017",
    "Whole blood\nACTU-3500", "PBMC\nACTU-3500", "PBMC\nGMS B", "Vagina\nGMS A")

tenofovir_levels <- c("PBMC\nGMS B", "PBMC\nGMS A", "Ectocervix\nGMS A", "Vagina\nGMS A")

both_drug_levels <- unique(c(truvada_levels, tenofovir_levels))

both_drug_levels_with_drug <- c(paste0("TDF\n", tenofovir_levels), rev(paste0("TDF/FTC\n", truvada_levels)))

set_up_truvada_facets <- function(d) {
    d %>% 
        filter(Treatment == "TDF/FTC") %>% 
        mutate(facetter = factor(facetter, levels = truvada_levels))
}

set_up_tenofovir_facets <- function(d) {
    d %>% 
        filter(Treatment == "TDF") %>% 
        mutate(facetter = factor(facetter, levels = tenofovir_levels)) 
}

# creates a plot of p-values as bar charts for gene set testing
gene_set_plot <- function(d) {
    
}

plot_correlation <- function(p, xlab, ylab) {
    p + 
        geom_point(size = 2) + 
        ylab(ylab) + xlab(xlab) + 
        coord_quickmap() + 
        geom_hline(yintercept = 0, color = highlightLineColor()) + 
        geom_vline(xintercept = 0, color = highlightLineColor()) + 
        theme_pub()
}

plot_vs_mtn_017 <- function(d, xlab) {
    d %>% 
        mutate(Significant = factor(Significant, levels = c("Both", "MTN-017", "ACTU-3500", "Neither"))) %>% 
        arrange(desc(Significant)) %>% 
        ggplot(aes(ACTU3500, MTN017, color = Significant)) %>%  
        plot_correlation(xlab, "Fold change (log2) in\nrectum in MTN-017") + 
        ylim(c(-1.6, 2.61)) + xlim(c(-1.6, 2.61))
}

plot_oral_deg_enrichment <- function(regex) {
    get_deg_sets_camera() %>% 
        filter(str_detect(Data_Tested, regex), !str_detect(Gene_Set, "\\&")) %>% 
        mutate(Data_Tested = factor(Data_Tested, levels = both_drug_levels)) %>% 
        mutate(Gene_Set = factor(Gene_Set, levels = sort(unique(Gene_Set))[3:1])) %>% 
        gene_set_plot()
}

plot_microarray_vs_ddpcr <- function(d, ncol = 3) {
    ggplot(d, aes(x = TargetId, y = Diff, color = Assay)) + 
        geom_hline(yintercept = 0, color = highlightLineColor()) + 
        stat_summary(fun.data = "mean_cl_normal", geom = "pointrange", 
            position = position_dodge(width = 0.5), fill = "white") + 
        facet_wrap(~ facetter, ncol = ncol, scales = "free_x") +
        ylab("Fold changes (log2)") +
        xlab(NULL) + 
        # control breaks with scale
        scale_y_continuous(breaks = c(-2, 0, 2, 4), labels = c(-2, 0, 2, 4)) + 
        # control zoom with coord_cartesian to not remove points involved in calculation
        # of the summary statistics
        # found limits to use with ggplot_build(p)$layout$panel_scales_y
        coord_cartesian(ylim = c(-2.1, 4.73)) + 
        theme_pub() + x_axis_all_facets() + 
        y_axis_all_facets() + 
        scale_color_brewer(palette = "Set1", guide = FALSE)
}

plot_hallmarks <- function(regex, renamer) {
    d <- to_plot %>% filter(str_detect(Data_Tested, regex)) %>% 
        group_by(Gene_Set) %>% 
        filter(sum(FDR < 0.05) > 1) %>% 
        mutate(Data_Tested = factor(Data_Tested, levels = both_drug_levels)) %>% 
        ungroup() %>% 
        mutate(Gene_Set = renamer[Gene_Set]) 
    
    p <- d %>% 
        ungroup() %>% 
        mutate(zzz = paste(Treatment, Data_Tested, sep = "\n"), 
            zzz = factor(zzz, levels = both_drug_levels_with_drug)) %>%
        ggplot(aes(x = Gene_Set, y = -log10(FDR), alpha = FDR < 0.05, 
            fill = Direction, color = Direction)) + 
        geom_col(width = 0.8) + 
        scale_fill_manual(values = c("#1b9e77", "#d95f02")) + 
        scale_color_manual(values = c("#1b9e77", "#d95f02")) + 
        scale_alpha_manual(values = c("FALSE" = 0, "TRUE" = 1), guide = FALSE) +
        ylab(fdr_neg_log10) + xlab("Gene set") +  
        tilt_x_labels(90, 0.5, 0.5) + 
        geom_hline(yintercept = -log10(0.05), color = highlightLineColor()) +
        theme_pub(strip.text.y = element_text(angle = 0, vjust = 0)) + 
        y_axis_all_facets() + x_axis_all_facets() + xlab(NULL)
    
    if ("Category" %in% colnames(d)) {
        p + facet_grid(zzz ~ Category, scales = "free_x", space = "free_x")
    } else {
        p + facet_grid(zzz ~ ., scales = "free_x", space = "free_x") 
    }
}
```
