## Supplementary material for "Treatment with commonly used antiretroviral drugs induces a type I/III interferon signature in the gut in the absence of HIV infection": Code to reproduce analyses and figures: figures.R

```
source("R/source_all.R")

# correlation of microarray fold changes ----------------------------------------------------
to_plot <- get_gene_fold_changes()

extract_data <- function(d) {
    d %>% 
        group_by(TargetId) %>% 
        # only take genes which are detected in both studies
        filter(n() > 1) %>% 
        arrange(Study) %>% 
        summarise(MTN017 = Log2FoldChange[2], ACTU3500 = Log2FoldChange[1], Significant = 
                case_when(all(AdjustedPValue < 0.05) ~ "Both", 
                    AdjustedPValue[2] < 0.05 ~ "MTN-017", 
                    AdjustedPValue[1] < 0.05 ~ "ACTU-3500", 
                    TRUE ~ "Neither"))
    
}

p1 <- to_plot %>%
    filter(Sample == "Rectum") %>%
    extract_data() %>% 
    plot_vs_mtn_017("Fold change (log2) in\nrectum in ACTU-3500") + 
    scale_color_manual(values = c("#e41a1c", "grey"), guide = FALSE) 

p2 <- to_plot %>%
    filter(Sample == "Duodenum" | Study == "MTN-017") %>%
    extract_data() %>% 
    plot_vs_mtn_017("Fold change (log2) in\nduodenum in ACTU-3500") + 
        scale_color_manual(values = c("#984ea3", "#e41a1c", "#377eb8", "grey"), name = "FDR < 0.05") + 
        ylab(NULL)

# combine with patchwork
p1 + p2

ggsave("figures/images/mtn017_actu3500_correlations.png", width = 6, height = 3, dpi = 600)

# participant level plots ------------------------------------------------------------------------

p <- get_probe_expression_participant_level()

to_plot <- p %>% 
    filter(Study %in% c("MTN-017", "ACTU-3500"), 
        TargetId %in% c("IFI6", "ISG15", "MX1"), 
        Sample %in% c("Duodenum", "Rectum")) %>% 
    # remove one of the two IFI6 probes
    filter(ProbeId != 5090215) %>% 
    mutate(Color = ifelse(Study == "MTN-017", "MTN-017", Ptid))

to_plot %>% 
    mutate(Visit = str_replace(Visit, " ", "\n"), facetter = paste(Study, Sample, sep = "\n")) %>% 
    ggplot(aes(x = Visit, y = Expression)) + 
        geom_line(aes(group = Ptid, color = Color), size = thinLineSize()) +
        geom_point(aes(color = Color), size = smallPointSize())  +
        styled_range() + styled_point() + 
        facet_grid(TargetId ~ facetter, scales = "free_y") + 
        theme_pub() + y_axis_all_facets() + x_axis_all_facets() + 
        xlab(NULL) + ylab("Expression (log2)") + 
        scale_color_manual(values = c("grey75", '#1b9e77','#d95f02','#7570b3','#e7298a','#66a61e','#e6ab02','#a6761d','#666666'), 
            guide = FALSE)

ggsave("figures/images/mtn_017_actu_3500_participant_level.png", width = 4.5, height = 3.5, dpi = 600)

# enrichment of oral DEGs ------------------------------------------------------------------------
get_deg_sets_camera() %>% 
    filter(str_detect(Gene_Set, "MTN-017")) %>% 
    mutate(Gene_Set = factor(Gene_Set, levels = sort(unique(Gene_Set))[3:1])) %>% 
    ungroup() %>% 
    mutate(zzz = paste(Treatment, Data_Tested, sep = "\n"), 
        zzz = factor(zzz, levels = c("TDF/FTC\nRectum\nACTU-3500", "TDF/FTC\nDuodenum\nACTU-3500",
            "TDF\nEctocervix\nGMS A", "TDF\nVagina\nGMS A", "TDF/FTC\nVagina\nGMS A", 
             "TDF\nPBMC\nGMS A",  
            "TDF\nPBMC\nGMS B", "TDF/FTC\nPBMC\nGMS B", "TDF/FTC\nPBMC\nACTU-3500", "TDF/FTC\nWhole blood\nACTU-3500"
        ))) %>%
    mutate(Tissue = ifelse(str_detect(zzz, "B|b"), "Blood", "Tissue"), 
        Tissue = factor(Tissue, levels = c("Tissue", "Blood"))) %>% 
    ggplot(aes(x = zzz, y = -log10(FDR), alpha = FDR < 0.05, 
        fill = Direction, color = Direction)) + 
    geom_col(width = 0.8) + 
    scale_fill_manual(values = c("#1b9e77", "#d95f02"), guide = FALSE) + 
    scale_color_manual(values = c("#1b9e77", "#d95f02"), guide = FALSE) + 
    scale_alpha_manual(values = c("FALSE" = 0, "TRUE" = 1), guide = FALSE) +
    ylab("FDR-adjusted p-value (-log10)") + xlab(NULL) +
    tilt_x_labels(90, 0.5, 0.5) + theme_pub() + 
    geom_hline(yintercept = -log10(0.05), color = highlightLineColor()) +
    facet_grid(.~Tissue, scales = "free_x", space = "free_x") +
    y_axis_all_facets()

ggsave("figures/images/mtn_017_deg_set_camera.png", width = 5.25, height = 2.5, dpi = 600)

# ddpcr vs microarray --------------------------------------------------------------------------------------
microarray_ddpcr <- get_microarray_vs_ddpcr() %>%
    filter(TargetId != "UBC") %>% 
    gather(Assay, Diff, MicroarrayDiff, PcrDiff) %>% 
    mutate(Assay = ifelse(Assay == "PcrDiff", "ddPCR", "Microarray"))

## RNAseq
rnaseq <- get_rnaseq_expression_participant_level() %>% 
    filter(TargetId %in% c("MX1", "IFI6", "ISG15")) %>% 
    mutate(Assay = "RNAseq") %>% 
    group_by(Assay, Study, Treatment, Sample, TargetId, Ptid) %>% 
    arrange(Visit) %>% 
    summarise(Diff = Expression[2] - Expression[1])

proteomics <- get_thai_proteomics_expression_participant_level() %>% 
    filter(TargetId %in% c("MX1", "IFI6", "ISG15")) %>% 
    mutate(Assay = "Proteomics") %>% 
    group_by(Assay, Study, Treatment, Sample, TargetId, Ptid) %>% 
    arrange(Visit) %>% 
    summarise(Diff = Expression[2] - Expression[1])

to_plot <- bind_rows(microarray_ddpcr, rnaseq, proteomics) %>% 
    mutate(facetter = paste(Sample, Study, sep = "\n")) 

to_plot %>% 
    set_up_truvada_facets %>% 
    plot_microarray_vs_ddpcr() 

ggsave("figures/images/microarray_vs_ddpcr_truvada.png", width = 4, height = 3.5, dpi = 600)

to_plot %>% 
    set_up_tenofovir_facets() %>% 
    plot_microarray_vs_ddpcr(ncol = 2)

ggsave("figures/images/microarray_vs_ddpcr_tenofovir.png", width = 2.5, height = 3.5, dpi = 600)

# Hallmark gene sets--------------------------------------------------
to_plot <- get_hallmarks_camera() %>% 
    inner_join(., read_csv("data/clean/hallmark_set_categories.csv", col_types = "cc"),
        by = "Gene_Set") %>% 
    mutate(Category = ifelse(Gene_Set == "TNFA_SIGNALING_VIA_NFKB", "immune", Category), 
        Category = ifelse(Category %in% c("proliferation", "immune"), Category, "other"),
        Category = str_to_sentence(Category),
        Category = factor(Category, levels = c("Immune", "Proliferation", "Other")))

renamer <- c("CHOLESTEROL_HOMEOSTASIS" = "Cholesterol homeostasis", 
    "UNFOLDED_PROTEIN_RESPONSE" = "Unfolded protein respose", 
    "E2F_TARGETS" = "E2F targets", 
    "ALLOGRAFT_REJECTION" = "Allograft rejection", 
    "G2M_CHECKPOINT" = "G2M checkpoint", 
    "OXIDATIVE_PHOSPHORYLATION" = "Oxidative phosphorylation", 
    "EPITHELIAL_MESENCHYMAL_TRANSITION" = "Epithelial mesenchymal\ntransition", 
    "INTERFERON_ALPHA_RESPONSE" = "IFN-\u03B1 response", 
    "MYC_TARGETS_V1" = "Myc targets v1", 
    "TNFA_SIGNALING_VIA_NFKB" = "TNF-\u03B1 signalling via NF-κB", 
    "MYC_TARGETS_V2" = "Myc targets v2", 
    "INTERFERON_GAMMA_RESPONSE" = "IFN-γ response", 
    "ANGIOGENESIS" = "Angiogenesis")

# blood
plot_hallmarks("(PBMC|blood)", renamer) + guides(fill = FALSE, color = FALSE) + ylim(c(0, 16)) + 
    theme_pub(axis.text.x = element_text(angle = 90, hjust = 1)) 
ggsave("figures/images/hallmarks_camera_blood.png", width = 1.8, height = 4.5, dpi = 600)

# tissue
plot_hallmarks("(um|ina|ix)", renamer) + ylim(c(0, 16)) + guides(fill = FALSE, color = FALSE) + 
    theme_pub(axis.text.x = element_text(angle = 90, hjust = 1)) 
ggsave("figures/images/hallmarks_camera_tissue.png", width = 4.75, height = 5, dpi = 600)

# Microscopy figures ------------------------------------------------------------------------------
isg15 <- get_isg15()

intensity <- isg15 %>% 
    filter(Class == "Bright") %>% 
    ggplot(aes(x = Visit, y = Intensity)) + 
    geom_line(aes(group = Ptid), color = "grey75", size = thinLineSize()) +
    geom_point(size = smallPointSize(), color = "grey75")  +
    styled_range() + styled_point() + 
    facet_wrap(~ Sample, ncol = 2, scales = "free_y") + 
    theme_pub() + 
    x_axis_all_facets() + y_axis_all_facets() + 
    ylab("Median intensity") + 
    xlab(NULL) + 
    # add white space to create a bit of gap when plots are combined
    add_white_space(bottom = 5)

percent <- get_clustered_isg15() %>% 
    mutate(Visit = str_replace(Visit, " ", "\n")) %>% 
    ggplot(aes(x = Visit, y = Percent)) + 
    geom_line(aes(group = Ptid), color = "grey75", size = thinLineSize()) +
    geom_point(size = smallPointSize(), color = "grey75")  +
    styled_range() + styled_point() + 
    facet_wrap(~ Sample, ncol = 2, scales = "free_y") + 
    theme_pub() + x_axis_all_facets() + y_axis_all_facets() + 
    ylab("Percentage of cells") + 
    xlab(NULL)

# combine with patchwork package
intensity + percent + plot_layout(ncol = 1)

ggsave("figures/images/microscopy_isg15.png", width = 3.5, height = 3.75, dpi = 600)

# proteomics from MTN-017 -------------------------------------------------------------------------------
to_plot <- get_thai_proteomics_vs_microarray() %>% 
    mutate(Significant = case_when(
        MicroarrayAdjustedPValue < 0.05 & PValue < 0.05 ~ "Both",
        MicroarrayAdjustedPValue < 0.05 ~ "Microarray", 
        PValue < 0.05 ~ "RNAseq FDR or\nprotein (unadjusted)", 
        TRUE ~ "Neither")) %>% 
    mutate(Significant = factor(Significant, levels = c("Both", "Microarray", "RNAseq FDR or\nprotein (unadjusted)", "Neither"))) %>% 
    arrange(desc(Significant))

proteomics_correlation <- ggplot(to_plot, aes(y = MicroarrayLog2FoldChange, x = Log2FoldChange, color = Significant)) %>%  
    plot_correlation("Protein fold change (log2)", "Microarray fold change (log2)") + ylab(NULL) +
        scale_color_manual(values = c("#984ea3", "#e41a1c", "#377eb8", "grey"), name = "P < 0.05") + xlim(c(-1, 1.6)) + ylim(c(-1, 1.6))

# rnaseq from MTN-017 -------------------------------------------------------------------------------
to_plot_rnaseq <- get_rnaseq_vs_microarray() %>% 
    mutate(Significant = case_when(
        MicroarrayAdjustedPValue < 0.05 & AdjustedPValue < 0.05 ~ "Both",
        MicroarrayAdjustedPValue < 0.05 ~ "Microarray", 
        AdjustedPValue < 0.05 ~ "RNAseq FDR or\nprotein (unadjusted)", 
        TRUE ~ "Neither")) %>% 
    mutate(Significant = factor(Significant, levels = c("Both", "Microarray", "RNAseq FDR or\nprotein (unadjusted)", "Neither"))) %>% 
    arrange(desc(Significant))

rnaseq_correlation <- ggplot(to_plot_rnaseq, aes(y = MicroarrayLog2FoldChange, x = Log2FoldChange, color = Significant)) %>%  
    plot_correlation("RNAseq fold change (log2)", "Microarray fold change (log2)") + 
    scale_color_manual(values = c("#984ea3", "#e41a1c", "#377eb8", "grey"), guide = FALSE) + xlim(c(-1, 1.6)) + ylim(c(-1, 1.6)) 

rnaseq_correlation + proteomics_correlation

ggsave("figures/images/rnaseq_proteomics_correlations.png", dpi = 600, width = 7, height = 3)
```
