## Supplementary material for "Treatment with commonly used antiretroviral drugs induces a type I/III interferon signature in the gut in the absence of HIV infection": Code to reproduce analyses and figures: generate_supplemental_data.R

```
source("R/source_all.R")

# Save gene lists for each study -----------------------------------------------------------------------------------
genes <- get_gene_fold_changes()

# function to remove the / in TDF/FTC for file names
sanitize_treatment <- function(x) str_replace(x, "/", "")

# make one data frame per treatment arm
nested_genes <- genes %>% 
    group_by(Study, Treatment, Sample) %>% 
    arrange(PValue) %>% 
    mutate(Name = paste0("Microarray_", unique(Study), "_", sanitize_treatment(unique(Treatment)), "_", unique(Sample))) %>% 
    group_by(Name) %>% 
    nest()

microarray_fold_changes <- nested_genes$data
names(microarray_fold_changes) <- nested_genes$Name

# add proteomics and RNAseq
other_fold_changes <- list("Proteomics_MTN-017_TDFFTC_Rectum" = get_thai_proteomics_gene_fold_changes() %>% arrange(PValue),
    "RNAseq_MTN-017_TDFFTC_Rectum" = get_rnaseq_gene_fold_changes() %>% arrange(PValue))

fold_changes <- c(other_fold_changes, microarray_fold_changes)

# Excel has a 31 character limit for worksheet names, need to rename some

full_names <- names(fold_changes)
short_names <- names(fold_changes) %>% str_sub(1, 31)
names(fold_changes) <- short_names

# create data dictionary        
data_dictionary <- tibble(
    Worksheet = short_names,
    Full = full_names
) %>% 
    separate(Full, into = c("Assay", "Study", "Drug", "Sample"), sep = "_") %>% 
    list("data_dictionary" = .)

fold_changes <- c(data_dictionary, fold_changes)

# save as Excel file with multiple worksheets
writexl::write_xlsx(fold_changes, "supplemental/fold_changes.xlsx")

# Save DEG sets gene set testing -----------------------------------------------------------------------------------
get_deg_sets_camera() %>% 
    mutate(Gene_Set = str_replace_all(Gene_Set, "\\n", "_")) %>% 
    rename(GeneSet = Gene_Set) %>% 
    select(-Data_Tested) %>% 
    arrange(FDR) %>% 
    write_csv("supplemental/Gene_set_testing_Microarray_DEGs_from_other_study_arms.csv")

# Save Hallmark gene set testing -----------------------------------------------------------------------------------
get_hallmarks_camera() %>% 
    rename(GeneSet = Gene_Set) %>% 
    select(-Data_Tested, -Sig) %>% 
    arrange(FDR) %>% 
    write_csv("supplemental/Gene_set_testing_Microarray_Hallmark_gene_sets.csv")

# overrepresentation analysis -----------------------------------------------------------------
# only load once the genes I might need
possible_genes <- get_gene_fold_changes() %>% 
    filter(!is.na(EntrezId), Sample %in% c("Duodenum", "Rectum"))

mtn_017_significant <- possible_genes %>% 
    filter(Study == "MTN-017", AdjustedPValue < 0.05, Log2FoldChange > 0) %>% 
    .$EntrezId
universe_genes <- possible_genes %>% filter(Study == "MTN-017") %>% .$EntrezId
mtn017_up <- limma::goana(mtn_017_significant, universe = universe_genes) %>% 
    arrange(P.DE) 

actu_3500_significant <- possible_genes %>% 
    filter(Study == "ACTU-3500", AdjustedPValue < 0.05)
universe_genes <- possible_genes %>% filter(Study == "ACTU-3500", Sample == "Duodenum") %>% .$EntrezId
down_genes <- actu_3500_significant %>% filter(Log2FoldChange < 0) %>% .$EntrezId
up_genes <- actu_3500_significant %>% filter(Log2FoldChange > 0) %>% .$EntrezId

duodenum_down <- limma::goana(down_genes, universe = universe_genes) %>% 
    arrange(P.DE) 
duodenum_up <- limma::goana(up_genes, universe = universe_genes) %>% 
    arrange(P.DE)

ora <- list("MTN-017_TDFFTC_Rectum_up" = mtn017_up, 
    "ACTU-3500_TDFFTC_Duodenum_down" = duodenum_down,
    "ACTU-3500_TDFFTC_Duodenum_up" = duodenum_up)

writexl::write_xlsx(ora, "supplemental/overrepresentation_analysis.xlsx")

# Supplementary table 1 ----------------------------------------------------------------------------------------
gmsa <- readxl::read_excel(microarray_GMSA_metadata) %>% mutate(Study = "GMS A") %>% filter(Paired == "Paired")
gmsb <- readxl::read_excel(microarray_GMSB_metadata) %>% mutate(Study = "GMS B", Tissue = "PBMC") %>% filter(Inclusion == "Included")
mtn017 <- readxl::read_excel(microarray_mtn017_metadata) %>% mutate(Study = "MTN-017", Tissue = "Rectal")
actu3500 <- readxl::read_excel(microarray_actu3500_metadata) %>% mutate(Study = "ACTU-3500")

rins <- data.table::rbindlist(list(gmsa, gmsb, mtn017, actu3500), fill = TRUE) %>% 
    select(Study, Sample = Tissue, RIN)

rins %>% 
    group_by(Study, Sample) %>% 
    summarise(n = n(), mean = round(mean(RIN), 1), min = min(RIN), max = max(RIN)) %>% 
    mutate(summary = paste0(mean, " (", min, "-", max, ")"))
```
