## Supplementary material for "Treatment with commonly used antiretroviral drugs induces a type I/III interferon signature in the gut in the absence of HIV infection": Code to reproduce analyses and figures: get_data.R

```
clean_file_path <- function(f) {
    here(paste0("data/clean/", f))
}

get_probe_fold_changes <- function() {
    read_csv(clean_file_path("probe_fold_changes.csv"),
        col_types = cols(
            Study = col_character(),
            Treatment = col_character(),
            TreatmentLength = col_character(),
            Sample = col_character(),
            ProbeId = col_integer(),
            TargetId = col_character(),
            EntrezId = col_integer(),
            Log2FoldChange = col_double(),
            PValue = col_double(),
            AdjustedPValue = col_double(),
            t = col_double()),
        progress = FALSE
    )
    
}

get_gene_fold_changes <- function() {
    read_csv(clean_file_path("gene_fold_changes.csv"),
        col_types = cols(
            Study = col_character(),
            Treatment = col_character(),
            TreatmentLength = col_character(),
            Sample = col_character(),
            TargetId = col_character(),
            EntrezId = col_integer(),
            Log2FoldChange = col_double(),
            PValue = col_double(),
            AdjustedPValue = col_double()
        ),
        progress = FALSE
    ) 
}

get_probe_expression_participant_level <- function() {
    readRDS(clean_file_path("probe_expression.Rdata")) %>% 
        as_tibble()
}

get_hallmarks_camera <- function() {
    read_csv(clean_file_path("hallmark_camera.csv"), 
        col_types = cols(
            NGenes = col_integer(),
            Direction = col_character(),
            PValue = col_double(),
            FDR = col_double(),
            Gene_Set = col_character(),
            Study = col_character(),
            Treatment = col_character(),
            TreatmentLength = col_character(),
            Sample = col_character(),
            Data_Tested = col_character(),
            Sig = col_integer()
        ))
}

get_deg_sets_camera <- function() {
    read_csv(clean_file_path("deg_sets_camera.csv"), 
        col_types = cols(
            NGenes = col_integer(),
            Direction = col_character(),
            PValue = col_double(),
            FDR = col_double(),
            Gene_Set = col_character(),
            Study = col_character(),
            Treatment = col_character(),
            TreatmentLength = col_character(),
            Sample = col_character(),
            Data_Tested = col_character()
        ))
}

get_microarray_vs_ddpcr <- function() {
    read_csv(clean_file_path("microarray_vs_ddpcr.csv"),
        col_types = cols(
            Study = col_character(),
            Treatment = col_character(),
            Sample = col_character(),
            TargetId = col_character(),
            Ptid = col_character(),
            MicroarrayDiff = col_double(),
            PcrDiff = col_double()
        )
    )
}

get_raw_isg15 <- function() {
    read_csv(clean_file_path("microscopy_isg15_raw.csv"),
        col_types = cols(
            Ptid = col_character(),
            Visit = col_character(),
            Sample = col_character(),
            SlideId = col_character(),
            Isg15Intensity = col_double(),
            Isg15Classification = col_double(),
            XMin = col_double(),
            XMax = col_double(),
            YMin = col_double(),
            YMax = col_double()
        ))
}

get_isg15 <- function() {
    read_csv(clean_file_path("microscopy_isg15_summarized.csv"), 
        col_types = cols(
            Ptid = col_character(),
            Visit = col_character(),
            Class = col_character(),
            Sample = col_character(),
            n = col_double(),
            Intensity = col_double(),
            Percent = col_double()
        ))
}


get_clustered_isg15 <- function() {
    read_csv(clean_file_path("microscopy_isg15_clustered_brights.csv"),
        col_types = cols(
            Ptid = col_character(),
            Visit = col_character(),
            Sample = col_character(),
            Isg15 = col_double(),
            Total = col_double(),
            Percent = col_double()
        ))
}

get_thai_proteomics_expression_set <- function() {
    load(clean_file_path("proteomics_thai.Rda"))
    thai_proteomics
}

get_thai_proteomics_gene_fold_changes <- function() {
    read_csv(clean_file_path("proteomics_thai_protein_fold_changes.csv"), col_types = cols(
        UniProtAccessionId = col_character(),
        TargetId = col_character(),
        EntrezId = col_double(),
        Log2FoldChange = col_double(),
        AveExpr = col_double(),
        t = col_double(),
        PValue = col_double(),
        AdjustedPValue = col_double(),
        B = col_double()
    ))
}

get_thai_proteomics_vs_microarray <- function() {
    get_gene_fold_changes() %>% 
        filter(Study == "MTN-017") %>% 
        setNames(paste0("Microarray", names(.))) %>% 
        inner_join(get_thai_proteomics_gene_fold_changes(), by = c("MicroarrayEntrezId" = "EntrezId"))
}

get_thai_proteomics_expression_participant_level <- function() {
    readRDS(clean_file_path("participant_level_expression/Proteomics_mtn017_expression_levels.Rdata")) %>% 
        as_tibble()
}

get_rnaseq_expression_participant_level <- function() {
    readRDS(clean_file_path("participant_level_expression/RNAseq_mtn017_expression_levels.Rdata")) %>% 
        as_tibble()
}

get_rnaseq_gene_fold_changes <- function() {
    read_csv(clean_file_path("rnaseq_gene_fold_changes.csv"), col_types = cols(
        seqname = col_character(),
        source = col_character(),
        feature = col_character(),
        start = col_double(),
        end = col_double(),
        score = col_character(),
        strand = col_character(),
        frame = col_character(),
        GeneId = col_character(),
        TargetId = col_character(),
        PId = col_character(),
        TranscriptId = col_character(),
        TssId = col_character(),
        EntrezId = col_double(),
        Log2FoldChange = col_double(),
        AveExpr = col_double(),
        t = col_double(),
        PValue = col_double(),
        AdjustedPValue = col_double(),
        B = col_double()
    ))
}

get_rnaseq_vs_microarray <- function() {
    get_gene_fold_changes() %>% 
        filter(Study == "MTN-017") %>% 
        setNames(paste0("Microarray", names(.))) %>% 
        inner_join(get_rnaseq_gene_fold_changes(), by = c("MicroarrayEntrezId" = "EntrezId", "MicroarrayTargetId" = "TargetId"))
}
```
