## Supplementary material for "Treatment with commonly used antiretroviral drugs induces a type I/III interferon signature in the gut in the absence of HIV infection": Code to reproduce analyses and figures: microarray_analysis_helpers.R

```
read_lumi <- function(file) {
    lumiR(file, detectionTh = 0.05, convertNuID = FALSE, checkDupId = FALSE,
        columnNameGrepPattern = list(exprs = 'AVG_SIGNAL', se.exprs = 'BEAD_STDERR',
            detection = 'Detection Pval', beadNum = 'Avg_NBEADS'),
        annotationColumn = c('ILMN_GENE', 'ENTREZ_GENE_ID', 'GI', 'ACCESSION', 'SYMBOL', 'PROBE_ID', 'PROBE_START', 'PROBE_SEQUENCE', 'CHROMOSOME', 'PROBE_CHR_ORIENTATION', 'PROBE_COORDINATES'),
        verbose = FALSE)
}

# adds pData to eSet, ordering pData by eSet
# pData must have a column called MicroarrayId that corresponds to the column
#   names of eSet
add_pData <- function(eSet, pData) {
    rownames(pData) <- pData$MicroarrayId
    stopifnot(nrow(pData) == ncol(eSet))
    
    # put pData in the order that eSet is in
    pData <- pData[order(match(rownames(pData), colnames(eSet))), ]
    stopifnot(all(rownames(pData) == colnames(eSet)))
    
    pData(eSet) <- pData
    
    eSet
}

pre_process_and_filter <- function(eSet, threshold) {
    eSet <- eSet %>%
        lumiB(method = "bgAdjust") %>%
        lumiT(method = "vst") %>%
        lumiN(method = "rsn") %>%
        lumiQ(detectionTh = 0.05)
    
    expressed <- rowSums(detection(eSet) < 0.05) >= threshold
    
    eSet[expressed, ]
}

save_top_table <- function(fit, coef, file, assay) {
    tt <- topTable(fit, coef = coef, n = Inf)
    
    write.csv(tt, file = paste0("data/clean/top_tables/", assay, "_", file, ".csv"), row.names = FALSE)
    
    invisible(NULL)
}

save_camera <- function(result, gene_sets_name, cond, assay) {
    result$Gene_Set <- rownames(result)
    result$Data_Tested <- cond
    rownames(result) <- NULL
    
    write.csv(result, file = paste0("data/clean/camera/", assay, "_", gene_sets_name, "_", cond, ".csv")) 
    
    invisible(NULL)
}

# loads the hallmark gene sets from msigdbr and puts them into the
# format required for limma::camera
get_hallmark_gene_sets_for_camera <- function() {
    # load hallmark genes
    df <- msigdbr::msigdbr(category = "H")
    
    # convert to form desired by camera
    temp <- df %>% group_by(gs_name) %>% summarize(entrez = list(entrez_gene))    
    result <- temp$entrez
    names(result) <- temp$gs_name
    
    result
}

make_expression_levels <- function(eSet, key) {
    data <- reshape2::melt(exprs(eSet))
    colnames(data) <- c("ProbeId", "MicroarrayId", "Expression")
    
    key <- key %>% 
        canonicalize_sample_names() %>% 
        canonicalize_study_names() %>% 
        canonicalize_treatment_names() %>% 
        canonicalize_visit_names()
    
    data <- merge(data, key, by = "MicroarrayId")
    
    features <- eSet %>% 
        fData() %>% 
        select(-SYMBOL) %>% 
        canonicalize_gene_symbols(entrez_id_column = "ENTREZ_GENE_ID", current_symbols_column = "TargetID") %>% 
        transmute(ProbeId = ProbeID, Gene = TargetId, EntrezId = ENTREZ_GENE_ID)
    
    merge(data, features, by = "ProbeId")
}
```
