## Supplementary material for "Treatment with commonly used antiretroviral drugs induces a type I/III interferon signature in the gut in the absence of HIV infection": Code to reproduce analyses and figures: package-citations.bib

@Manual{,
  title = {conflicted: An Alternative Conflict Resolution Strategy},
  author = {Hadley Wickham},
  year = {2018},
  note = {R package version 1.0.1},
  url = {https://CRAN.R-project.org/package=conflicted},
}
@Article{,
  title = {edgeR: a Bioconductor package for differential expression analysis of digital gene expression data},
  author = {Mark D Robinson and Davis J McCarthy and Gordon K Smyth},
  journal = {Bioinformatics},
  volume = {26},
  number = {1},
  pages = {139-140},
  year = {2010},
}

@Article{,
  author = {{McCarthy} and Davis J. and {Chen} and {Yunshun} and {Smyth} and Gordon K.},
  title = {Differential expression analysis of multifactor RNA-Seq experiments with respect to biological variation},
  year = {2012},
  journal = {Nucleic Acids Research},
  volume = {40},
  number = {10},
  pages = {4288-4297},
}
@Manual{,
  title = {msigdbr: MSigDB Gene Sets for Multiple Organisms in a Tidy Data Format},
  author = {Igor Dolgalev},
  year = {2018},
  note = {R package version 6.2.1},
  url = {https://CRAN.R-project.org/package=msigdbr},
}
@Manual{,
  title = {patchwork: The Composer of ggplots},
  author = {Thomas Lin Pedersen},
  year = {2017},
  note = {R package version 0.0.1},
  url = {https://github.com/thomasp85/patchwork},
}
