## Supplementary material for "Treatment with commonly used antiretroviral drugs induces a type I/III interferon signature in the gut in the absence of HIV infection": Code to reproduce analyses and figures: source_all.R

```
library(tidyverse)
library(broom)
library(here)
library(pander)
library(plater)
library(conflicted)
library(readxl)
library(ggrepel)
library(writexl)
library(lumi) # Bioconductor
library(limma) # Bioconductor
library(Biobase) # Bioconductor
library(msigdbr) # Bioconductor
library(edgeR) # Bioconductor
library(patchwork) # https://github.com/thomasp85/patchwork

# if "filter" or "select" is called, assume it's from dplyr
# prevents conflicts with bioconductor packages
conflict_prefer("filter", "dplyr", quiet = TRUE)
conflict_prefer("select", "dplyr", quiet = TRUE) 
conflict_prefer("rename", "dplyr", quiet = TRUE)
conflict_prefer("filter", "dplyr", quiet = TRUE)

source(here("R/util.R"))
source(here("R/style_functions.R"))
source(here("R/get_data.R"))
source(here("R/figure_helpers.R"))
source(here("R/microarray_analysis_helpers.R"))

# generate BIBTEX file of citations for packages
# packages <- c("AnnotationDbi", "Biobase", "broom", "ggrepel", "here", "limma", "lumi", "org.Hs.eg.db", "pander", "plater", "RColorBrewer", "tidyverse")
# cite <- function(x) as.character(toBibtex(citation(x)))
# f <- file("package-citations.bib")
# lapply(packages, cite) %>%
#    unlist() %>%
#    writeLines(f)
# close(f)
```
