## Supplementary material for "Treatment with commonly used antiretroviral drugs induces a type I/III interferon signature in the gut in the absence of HIV infection": Code to reproduce analyses and figures: statistics.pdf

### Oral tenofovir: statistics

#### Differentially expressed genes

Table A: Number of differentially expressed genes from each study

| Study | Treatment | Sample | up | down | total |
|---|---|---|---|---|---|
| ACTU-3500 | TDF/FTC | Duodenum | 116 | 135 | 16321 |
| ACTU-3500 | TDF/FTC | PBMC | 0 | 0 | 14937 |
| ACTU-3500 | TDF/FTC | Rectum | 0 | 0 | 16399 |
| ACTU-3500 | TDF/FTC | Whole blood | 0 | 0 | 13922 |
| GMS A | TDF | Ectocervix | 0 | 0 | 17578 |
| GMS A | TDF | PBMC | 0 | 0 | 16553 |
| GMS A | TDF | Vagina | 0 | 0 | 18397 |
| GMS A | TDF/FTC | Vagina | 0 | 0 | 18397 |
| GMS B | TDF | PBMC | 0 | 0 | 20121 |
| GMS B | TDF/FTC | PBMC | 0 | 0 | 20121 |
| MTN-017 | TDF/FTC | Rectum | 13 | 0 | 21683 |

Table B: Names of all differentially expressed genes from the rectal biopsies after oral administration in MTN-017. All genes were upregulated.

| EntrezId | Symbol | Name | Type1Interferon | Log2FoldChange | AdjustedPValue |
|---|---|---|---|---|---|
| 3429 | IFI27 | interferon alpha inducible protein 27 | TRUE | 0.76 | 3.54e-07 |
| 2537 | IFI6 | interferon alpha inducible protein 6 | TRUE | 0.8 | 2.27e-04 |
| 3434 | IFIT1 | interferon induced protein with tetratricopeptide repeats 1 | TRUE | 0.75 | 2.49e-03 |
| 9636 | ISG15 | ISG15 ubiquitin like modifier | TRUE | 0.98 | 2.49e-03 |
| 91543 | RSAD2 | radical S-adenosyl methionine domain containing 2 | TRUE | 0.53 | 3.14e-03 |
| 4599 | MX1 | MX dynamin like GTPase 1 | TRUE | 0.92 | 3.60e-03 |
| 4938 | OAS1 | 2'-5'-oligoadenylate synthetase 1 | TRUE | 0.32 | 0.02 |
| 122509 | IFI27L1 | interferon alpha inducible protein 27 like 1 | FALSE | 0.25 | 2.27e-04 |
| 55601 | DDX60 | DExD/H-box helicase 60 | FALSE | 0.39 | 2.49e-03 |
| 54809 | SAMD9 | sterile alpha motif domain containing 9 | FALSE | 0.35 | 0.01 |
| 647215 | MROH3P | maestro heat like repeat family member 3, pseudogene | FALSE | 0.22 | 0.02 |
| 84318 | CCDC77 | coiled-coil domain containing 77 | FALSE | 0.12 | 0.05 |
| 55008 | HERC6 | HECT and RLD domain containing E3 ubiquitin protein ligase family member 6 | FALSE | 0.31 | 0.05 |

#### Gene fold change correlations

Table C: Spearman correlation coefficients comparing the fold changes in the rectum in ACTU-3500 vs. the rectum in MTN-017

| Genes | Spearman |
|---|---|
| All detected | 0.07 |
| Significant in MTN-017 | 0.91 |
| Significant in neither | 0.07 |

Table D: Spearman correlation coefficients comparing the fold changes in the duodenum in ACTU-3500 vs. the rectum in MTN-017

| Genes | Spearman |
|---|---|
| All detected | 0.06 |
| Significant in MTN-017 | 0.81 |
| Significant in ACTU-3500 | 0.13 |
| Significant in neither | 0.06 |

#### Gene fold changes in ACTU-3500 rectum and duodenum of genes that were significant in MTN-017

Table E: Fold changes and p-values in ACTU-3500 rectum for the genes that were significant in MTN-017

| Study | Treatment | TreatmentLength | Sample | TargetId | EntrezId | Log2FoldChange | PValue | AdjustedPValue |
|---|---|---|---|---|---|---|---|---|
| ACTU-3500 | TDF/FTC | 60Days | Rectum | ISG15 | 9636 | 0.8645 | 0.02155 | 0.9584 |
| ACTU-3500 | TDF/FTC | 60Days | Rectum | DDX60 | 55601 | 0.3236 | 0.03398 | 0.9584 |
| ACTU-3500 | TDF/FTC | 60Days | Rectum | IFI27L1 | 122509 | 0.2601 | 0.05889 | 0.9584 |
| ACTU-3500 | TDF/FTC | 60Days | Rectum | IFI6 | 2537 | 0.536 | 0.08276 | 0.9584 |
| ACTU-3500 | TDF/FTC | 60Days | Rectum | IFIT1 | 3434 | 0.444 | 0.0853 | 0.9584 |
| ACTU-3500 | TDF/FTC | 60Days | Rectum | RSAD2 | 91543 | 0.405 | 0.09039 | 0.9584 |
| ACTU-3500 | TDF/FTC | 60Days | Rectum | MX1 | 4599 | 0.4615 | 0.2174 | 0.9584 |
| ACTU-3500 | TDF/FTC | 60Days | Rectum | IFI27 | 3429 | 0.3208 | 0.2617 | 0.9584 |
| ACTU-3500 | TDF/FTC | 60Days | Rectum | OAS1 | 4938 | 0.2479 | 0.4508 | 0.963 |
| ACTU-3500 | TDF/FTC | 60Days | Rectum | SAMD9 | 54809 | 0.1475 | 0.7451 | 0.9839 |
| ACTU-3500 | TDF/FTC | 60Days | Rectum | MROH3P | 647215 | 0.04134 | 0.8122 | 0.9868 |
| ACTU-3500 | TDF/FTC | 60Days | Rectum | CCDC77 | 84318 | 0.003953 | 0.9596 | 0.9961 |

| Study | Treatment | TreatmentLength | Sample | TargetId | EntrezId | Log2FoldChange | PValue | AdjustedPValue |
|---|---|---|---|---|---|---|---|---|
| ACTU-3500 | TDF/FTC | 60Days | Rectum | HERC6 | 55008 | 0.007217 | 0.9803 | 0.9977 |

Table F: Fold changes and p-values in ACTU-3500 duodenum for the genes that were significant in MTN-017

| Study | Treatment | TreatmentLength | Sample | TargetId | EntrezId | Log2FoldChange | PValue | AdjustedPValue |
|---|---|---|---|---|---|---|---|---|
| ACTU-3500 | TDF/FTC | 60Days | Duodenum | IFI27L1 | 122509 | 0.6446 | 0.0003644 | 0.04039 |
| ACTU-3500 | TDF/FTC | 60Days | Duodenum | CCDC77 | 84318 | 0.1807 | 0.001141 | 0.06002 |
| ACTU-3500 | TDF/FTC | 60Days | Duodenum | IFI6 | 2537 | 2.404 | 0.001341 | 0.06153 |
| ACTU-3500 | TDF/FTC | 60Days | Duodenum | MX1 | 4599 | 1.584 | 0.001438 | 0.06371 |
| ACTU-3500 | TDF/FTC | 60Days | Duodenum | IFIT1 | 3434 | 2.512 | 0.001442 | 0.06371 |
| ACTU-3500 | TDF/FTC | 60Days | Duodenum | HERC6 | 55008 | 1.431 | 0.001999 | 0.06903 |
| ACTU-3500 | TDF/FTC | 60Days | Duodenum | IFI27 | 3429 | 1.39 | 0.003127 | 0.07828 |
| ACTU-3500 | TDF/FTC | 60Days | Duodenum | RSAD2 | 91543 | 1.812 | 0.00319 | 0.07862 |
| ACTU-3500 | TDF/FTC | 60Days | Duodenum | ISG15 | 9636 | 2.001 | 0.003706 | 0.085 |
| ACTU-3500 | TDF/FTC | 60Days | Duodenum | DDX60 | 55601 | 0.8462 | 0.008887 | 0.1206 |
| ACTU-3500 | TDF/FTC | 60Days | Duodenum | SAMD9 | 54809 | 1.038 | 0.01016 | 0.129 |
| ACTU-3500 | TDF/FTC | 60Days | Duodenum | OAS1 | 4938 | 0.9519 | 0.01713 | 0.1642 |
| ACTU-3500 | TDF/FTC | 60Days | Duodenum | MROH3P | 647215 | 0.2452 | 0.04936 | 0.2716 |

#### Correlations between protein and transcript

Table G: Spearman correlation coefficients comparing the fold changes in the rectum in MTN-017 as detected by microarray and proteomics

| Genes | Spearman |
|---|---|
| All detected | 0.07 |
| Significant by microarray | 0.8 |
| Unadjusted significant in proteomics | 0.05 |
| Significant in neither | 0.06 |

#### Correlations between RNAseq and microarray

Table H: Spearman correlation coefficients comparing the fold changes in the rectum in MTN-017 as detected by microarray and RNAseq

| Genes | Spearman |
|---|---|
| All detected | 0.34 |
| Significant by microarray | 0.84 |
| Significant in neither | 0.34 |

#### ddPCR and microarray correlations

Table I: Pearson correlation coefficients of fold changes measured by ddPCR and microarray.

| TargetId | Pearson |
|---|---|
| IFI6 | 0.8956 |
| ISG15 | 0.9165 |
| MX1 | 0.9269 |

Table J: Pearson correlation coefficients of fold changes measured by ddPCR and microarray.

| Study | Treatment | Sample | IFI6 | ISG15 | MX1 |
|---|---|---|---|---|---|
| ACTU-3500 | TDF/FTC | Duodenum | 0.9601 | 0.9856 | 0.921 |
| ACTU-3500 | TDF/FTC | PBMC | 0.9793 | 0.9372 | 0.9624 |
| ACTU-3500 | TDF/FTC | Rectum | 0.8728 | 0.9114 | 0.9176 |
| ACTU-3500 | TDF/FTC | Whole blood | 0.988 | 0.9936 | 0.9609 |
| GMS A | TDF | Ectocervix | 0.5846 | 0.8238 | 0.7606 |
| GMS A | TDF | PBMC | 0.8073 | 0.8487 | 0.744 |
| GMS A | TDF | Vagina | 0.633 | 0.5376 | 0.544 |
| MTN-017 | TDF/FTC | Rectum | 0.9347 | 0.9782 | 0.976 |

Table K: Summary of Pearson correlation cofficients of fold changes.

| Min. | 1st Qu. | Median | Mean | 3rd Qu. | Max. |
|---|---|---|---|---|---|
| 0.5376 | 0.7956 | 0.9193 | 0.8568 | 0.9658 | 0.9936 |

#### ddPCR effect sizes

Table L: Mean and 95% CI of fold changes from ddPCR

| Study | Sample | TargetId | mean | min | max |
|---|---|---|---|---|---|
| ACTU-3500 | Duodenum | IFI6 | 8.89 | 2.981 | 26.51 |
| ACTU-3500 | Duodenum | ISG15 | 6.577 | 1.803 | 24 |
| ACTU-3500 | Duodenum | MX1 | 3.774 | 1.997 | 7.131 |
| ACTU-3500 | Rectum | IFI6 | 1.871 | 1.014 | 3.451 |
| ACTU-3500 | Rectum | ISG15 | 1.901 | 1.089 | 3.316 |
| ACTU-3500 | Rectum | MX1 | 1.555 | 0.8633 | 2.799 |
| MTN-017 | Rectum | IFI6 | 2.883 | 1.997 | 4.164 |
| MTN-017 | Rectum | ISG15 | 2.33 | 1.628 | 3.333 |
| MTN-017 | Rectum | MX1 | 2.056 | 1.46 | 2.896 |

#### Microscopy

Table M: ISG15 intensity. Positive numbers indicate higher during treatment. One-sided paired t-test.

| Class | Sample | difference | ci | p.value | adjusted |
|---|---|---|---|---|---|
| Dim/Negative | Rectum | 0.001157 | [-0.01, Inf] | 0.37 | 1 |
| Dim/Negative | Duodenum | -0.000219 | [-0.01, Inf] | 0.52 | 1 |
| Bright | Rectum | -0.005367 | [-0.03, Inf] | 0.64 | 1 |
| Bright | Duodenum | -0.003227 | [-0.02, Inf] | 0.65 | 1 |

Table N: Percent of all cells that are bright for ISG15. Positive numbers indicate higher during treatment. One-sided paired t-test.

| Sample | difference | ci | p.value | adjusted |
|---|---|---|---|---|
| Rectum | 0.4256 | [0.25, Inf] | 2.34e-03 | 4.68e-03 |
| Duodenum | 0.4305 | [-0.04, Inf] | 0.06 | 0.12 |

Table O: Fold change of percent of all cells that are bright for ISG15. Positive numbers indicate higher during treatment. 1 indicates no change.

| Sample | Mean | sd | n |
|---|---|---|---|
| Duodenum | 1.367 | 0.5116 | 8 |
| Rectum | 2.756 | 0.984 | 6 |
