## Supplementary material for "Treatment with commonly used antiretroviral drugs induces a type I/III interferon signature in the gut in the absence of HIV infection": Code to reproduce analyses and figures: statistics.Rmd

````
---
title: "Oral tenofovir: statistics"
output: pdf_document
header-includes:
   - \usepackage{alphalph}
toc: true
---

\renewcommand{\thetable}{\AlphAlph{\value{table}}}

```{r, warning=FALSE, message=FALSE, echo=FALSE, results='hide'}
knitr::opts_chunk$set(echo = FALSE)
library(here)
source(here("R/source_all.R"))
```

\pagebreak

## Differentially expressed genes
```{r, echo=FALSE}
# number of genes
get_gene_fold_changes() %>% 
    group_by(Study, Treatment, Sample) %>% 
    summarise(up = sum(Log2FoldChange > 0 & AdjustedPValue < 0.05), 
        down = sum(Log2FoldChange < 0 & AdjustedPValue < 0.05), total = n()) %>% 
    pander(caption = "Number of differentially expressed genes from each study")
```

```{r, message=FALSE}
mtn_017_significant <- get_gene_fold_changes() %>% 
    filter(Study == "MTN-017", AdjustedPValue < 0.05) %>% 
    arrange(AdjustedPValue)

# get up to date entrez id, symbol, and name for all DEGs
gene_list <- AnnotationDbi::select(org.Hs.eg.db::org.Hs.eg.db, 
    keys = as.character(mtn_017_significant$EntrezId), 
    columns = c("ENTREZID", "SYMBOL", "GENENAME"))
# nicely relabel columns
names(gene_list) <- c("EntrezId", "Symbol", "Name")

# look up whether or not the genes are in the type I interferon signaling pathway
gene_ontology_type_one_interferon_signalling_pathway <- "GO:0060337"
go_lookup <- AnnotationDbi::toTable(org.Hs.eg.db::org.Hs.egGO)
ifn_a_pathway_genes <- go_lookup[go_lookup$go_id == gene_ontology_type_one_interferon_signalling_pathway, ]

gene_list <- gene_list %>% 
    mutate(Type1Interferon = EntrezId %in% ifn_a_pathway_genes$gene_id, EntrezId = as.integer(EntrezId)) %>% 
    arrange(!Type1Interferon, EntrezId) %>% 
    inner_join(mtn_017_significant, by = c("EntrezId")) %>% 
    select(EntrezId, Symbol, Name, Type1Interferon, Log2FoldChange, AdjustedPValue) %>% 
    arrange(!Type1Interferon, AdjustedPValue) %>% 
    mutate(Log2FoldChange = round(Log2FoldChange, 2), AdjustedPValue = format_p(AdjustedPValue, 3))

panderOptions("table.split.table", Inf)
pander(gene_list, 
    caption = "Names of all differentially expressed genes from the rectal biopsies after oral administration in MTN-017. All genes were upregulated.")
```

\pagebreak

## Gene fold change correlations

```{r,echo=FALSE}
find_cor <- function(d) round(cor(d$MTN017, d$ACTU3500, method = "spearman"), 2)

rec_vs_rec <- get_gene_fold_changes() %>% 
    filter(Sample == "Rectum") %>%
    group_by(TargetId) %>% 
    filter(n() > 1) %>% 
    arrange(Study) %>% 
    summarise(MTN017 = Log2FoldChange[2], ACTU3500 = Log2FoldChange[1], Significant = 
            case_when(all(AdjustedPValue < 0.05) ~ "Both", 
                AdjustedPValue[2] < 0.05 ~ "MTN017", 
                AdjustedPValue[1] < 0.05 ~ "ACTU3500", 
                TRUE ~ "Neither"))

data.frame(Genes = c("All detected", "Significant in MTN-017", "Significant in neither"),
    Spearman = c(find_cor(rec_vs_rec), find_cor(filter(rec_vs_rec, Significant %in% c("MTN017", "Both"))), 
        find_cor(filter(rec_vs_rec, Significant == "Neither")))) %>% 
    pander(caption = "Spearman correlation coefficients comparing the fold changes in the rectum in ACTU-3500 vs. the rectum in MTN-017")

duod_vs_rec <- get_gene_fold_changes() %>% 
    filter(Sample == "Duodenum" | Study == "MTN-017") %>%
    group_by(TargetId) %>% 
    filter(n() > 1) %>% 
    arrange(Study) %>% 
    summarise(MTN017 = Log2FoldChange[2], ACTU3500 = Log2FoldChange[1], Significant = 
            case_when(all(AdjustedPValue < 0.05) ~ "Both", 
                AdjustedPValue[2] < 0.05 ~ "MTN017", 
                AdjustedPValue[1] < 0.05 ~ "ACTU3500", 
                TRUE ~ "Neither"))

data.frame(Genes = c("All detected", "Significant in MTN-017", "Significant in ACTU-3500", "Significant in neither"),
    Spearman = c(find_cor(duod_vs_rec), find_cor(filter(duod_vs_rec, Significant %in% c("MTN017", "Both"))), 
        find_cor(filter(duod_vs_rec, Significant %in% c("ACTU3500", "Both"))), 
        find_cor(filter(duod_vs_rec, Significant == "Neither")))) %>% 
    pander(caption = "Spearman correlation coefficients comparing the fold changes in the duodenum in ACTU-3500 vs. the rectum in MTN-017")

```

## Gene fold changes in ACTU-3500 rectum and duodenum of genes that were significant in MTN-017

```{r}
g <- get_gene_fold_changes()
mtn_017_significant_entrez <- g %>% 
    filter(Study == "MTN-017", AdjustedPValue < 0.05) %>% 
    pull(EntrezId)

g %>% 
    filter(EntrezId %in% mtn_017_significant_entrez, Study == "ACTU-3500", Sample == "Rectum") %>% 
    arrange(PValue) %>% 
    pander(caption = "Fold changes and p-values in ACTU-3500 rectum for the genes that were significant in MTN-017 ")

g %>% 
    filter(EntrezId %in% mtn_017_significant_entrez, Study == "ACTU-3500", Sample == "Duodenum") %>% 
    arrange(PValue) %>% 
    pander(caption = "Fold changes and p-values in ACTU-3500 duodenum for the genes that were significant in MTN-017 ")

```


## Correlations between protein and transcript 

```{r}
proteomics_vs_microarray <- get_thai_proteomics_vs_microarray()
find_cor <- function(d) round(cor(d$MicroarrayLog2FoldChange, d$Log2FoldChange, method = "spearman"), 2)

data.frame(Genes = c("All detected", "Significant by microarray", "Unadjusted significant in proteomics", "Significant in neither"),
    Spearman = c(find_cor(proteomics_vs_microarray), find_cor(filter(proteomics_vs_microarray, MicroarrayAdjustedPValue < 0.05)), 
        find_cor(filter(proteomics_vs_microarray, PValue < 0.05)), 
        find_cor(filter(proteomics_vs_microarray, PValue > 0.05, AdjustedPValue > 0.05)))) %>% 
    pander(caption = "Spearman correlation coefficients comparing the fold changes in the rectum in MTN-017 as detected by microarray and proteomics")
```

## Correlations between RNAseq and microarray

```{r}
rnaseq_vs_microarray <- get_rnaseq_vs_microarray()
find_cor <- function(d) round(cor(d$MicroarrayLog2FoldChange, d$Log2FoldChange, method = "spearman"), 2)

data.frame(Genes = c("All detected", "Significant by microarray", "Significant in neither"),
    Spearman = c(
        find_cor(rnaseq_vs_microarray), 
        find_cor(filter(rnaseq_vs_microarray, MicroarrayAdjustedPValue < 0.05)), 
        find_cor(filter(rnaseq_vs_microarray, MicroarrayAdjustedPValue > 0.05, AdjustedPValue > 0.05)))) %>% 
    pander(caption = "Spearman correlation coefficients comparing the fold changes in the rectum in MTN-017 as detected by microarray and RNAseq")
```

\pagebreak

## ddPCR and microarray correlations
```{r}
get_microarray_vs_ddpcr() %>% 
    filter(TargetId != "UBC") %>% 
    group_by(TargetId) %>% 
    summarise(Pearson=cor(MicroarrayDiff, PcrDiff)) %>% 
    pander(caption = "Pearson correlation coefficients of fold changes measured by ddPCR and microarray.")

get_microarray_vs_ddpcr() %>% 
    filter(TargetId != "UBC") %>% 
    group_by(Study, Treatment, Sample, TargetId) %>% 
    summarise(Pearson=cor(MicroarrayDiff, PcrDiff)) %>% 
    spread(TargetId, Pearson) %>% 
    pander(caption = "Pearson correlation coefficients of fold changes measured by ddPCR and microarray.")

get_microarray_vs_ddpcr() %>% 
    filter(TargetId != "UBC") %>% 
    group_by(Study, Treatment, Sample, TargetId) %>% 
    summarise(Pearson=cor(MicroarrayDiff, PcrDiff)) %>% .$Pearson %>% summary() %>% 
    pander(caption = "Summary of Pearson correlation cofficients of fold changes.")
```

## ddPCR effect sizes

```{r}
# reported in the abstract
get_microarray_vs_ddpcr() %>% 
    filter(Sample %in% c("Duodenum", "Rectum"), TargetId %in% c("MX1", "IFI6", "ISG15")) %>% 
    group_by(Study, Sample, TargetId) %>% 
    summarise(mean = 2^Hmisc::smean.cl.normal(PcrDiff)[1], 
        min = 2^Hmisc::smean.cl.normal(PcrDiff)[2], 
        max = 2^Hmisc::smean.cl.normal(PcrDiff)[3]) %>% 
        pander(caption = "Mean and 95% CI of fold changes from ddPCR")
```


\pagebreak

## Microscopy

```{r, echo=FALSE}
isg15 <- get_isg15() %>% 
    ungroup() %>% 
    arrange(Ptid, Visit)

isg15 %>% 
    nest(data = c(-Class, -Sample)) %>% 
    mutate(test = map(data, ~ tidy(t.test(.x$Intensity ~ .x$Visit, paired = TRUE, alternative = "less")))) %>% 
    unnest(test) %>% arrange(p.value) %>% 
    adjust_and_ci() %>% 
    select(Class, Sample, difference, ci, p.value, adjusted) %>% 
    pander(caption = "ISG15 intensity. Positive numbers indicate higher during treatment. One-sided paired t-test.")
```

```{r, echo=FALSE}
get_clustered_isg15() %>% 
    ungroup() %>% 
    arrange(Ptid, Visit) %>% 
    select(-Isg15, -Total) %>% 
    nest(data = -Sample) %>% 
    mutate(test = map(data, ~ tidy(t.test(.x$Percent ~ .x$Visit, paired = TRUE, alternative = "less")))) %>% 
    unnest(test) %>% 
    adjust_and_ci() %>% 
    select(Sample, difference, ci, p.value, adjusted) %>% 
    pander(caption = "Percent of all cells that are bright for ISG15. Positive numbers indicate higher during treatment. One-sided paired t-test.")

get_clustered_isg15() %>% 
    ungroup() %>% 
    arrange(Ptid, Visit) %>% 
    select(-Isg15, -Total) %>% 
    spread(Visit, Percent) %>% 
    mutate(FoldChange = `On treatment` / `Off treatment`) %>% 
    group_by(Sample) %>% 
    summarise(Mean = mean(FoldChange), sd = sd(FoldChange), n = n()) %>% 
    pander(caption = "Fold change of percent of all cells that are bright for ISG15. Positive numbers indicate higher during treatment. 1 indicates no change.")
```
````
