## Supplementary material for "Treatment with commonly used antiretroviral drugs induces a type I/III interferon signature in the gut in the absence of HIV infection": Code to reproduce analyses and figures: style_functions.R

```
smallPointSize <- function() 1
pointSize <- function() 2
lineSize <- function() 0.5
thinLineSize <- function() 0.25
axisLineColor <- function() "gray50"
highlightLineColor <- function() axisLineColor()
dpi <- function() 600
halfColumnWidth <- function() 3.75
figHeight <- function() 1.4
colorBlindPalette <- function() c("#009E73", "#D55E00", "#0072B2", "#CC79A7")

styled_range <- function() {
   stat_summary(fun.data = "mean_cl_normal", geom = "linerange", 
      size = lineSize()) 
}

styled_point <- function() {
   stat_summary(fun.y = "mean", geom = "point", size = pointSize())
}

theme_pub <- function(...) {
   theme(
      # backgrounds
      strip.background = element_blank(),
      panel.background = element_blank(),
      legend.key = element_blank(),
      
      # lines
      axis.line = element_line(color = axisLineColor(), size = lineSize()),
      axis.ticks = element_line(color = axisLineColor(), size = lineSize()),
      panel.grid.major = element_line(color = "grey90", size = thinLineSize()),
      panel.grid.minor = element_blank(),
      
      # fonts
      title = element_text(size = 12),
      axis.title = element_text(size = 10),
      strip.text = element_text(size = 10),
      legend.title = element_text(size = 10, face = "plain"),
      legend.text = element_text(size = 8, color = "grey50"),
      axis.text = element_text(size = 8),
      
      # remove whitespace at edges of plots
      plot.margin = grid::unit(c(0, 0, 0, 0), "mm"), 
      
      # center plot titles
      plot.title = element_text(hjust = 0.5),
      ...
   ) 
}

y_axis_all_facets <- function() {
   annotate("segment", color = axisLineColor(), 
      size = lineSize(),
      x = -Inf, xend = -Inf, y = -Inf, yend = Inf)
}

x_axis_all_facets <- function() {
   annotate("segment", color = axisLineColor(), 
      size = lineSize(),
      x = -Inf, xend = Inf, y = -Inf, yend = -Inf)
}

add_white_space <- function(top = 0, right = 0, bottom = 0, left = 0) {
   theme(plot.margin = grid::unit(c(top, right, bottom, left), "mm"))
}

tilt_x_labels <- function(angle = 45, hjust = 1, vjust = 1) {
    # sensible default for 90 degrees
    if (angle == 90 && missing(vjust)) {
        vjust <- 0.25
    }
    ggplot2::theme(axis.text.x =
            ggplot2::element_text(angle = angle, hjust = hjust, vjust = vjust))
}
```
