## Supplementary material for "Treatment with commonly used antiretroviral drugs induces a type I/III interferon signature in the gut in the absence of HIV infection": Code to reproduce analyses and figures: util.R

```
# enforce valid names
enforce_valid_names <- function(vector, valid, failing_message) {
    if (!all(vector %in% valid)) stop(failing_message)
}


# convert study names to canonical form for consistency
canonicalize_study_names <- function(d) {
    d <- d %>% 
        mutate(Study = case_when(
            Study == "MTN017" ~ "MTN-017", 
            Study == "PrEP" ~ "ACTU-3500", 
            Study == "ACTU3500" ~ "ACTU-3500", 
            str_detect(Study, "MEHP") ~ "GMS A",
            Study %in% c("CDC", "GMSB") ~ "GMS B",
            Study == "GMSA" ~ "GMS A",
            TRUE ~ Study
        ))
    
    # require valid names
    enforce_valid_names(d$Study, c("GMS A", "ACTU-3500", "GMS B", "MTN-017"), "Invalid study names")
    
    d
}

# convert sample names to canonical form for consistency
canonicalize_sample_names <- function(d) {
    d <- d %>% 
        mutate(Sample = case_when(
            Sample == "15cmRectum" ~ "Rectum", 
            Sample == "PAXgene" ~ "Whole blood",
            Sample == "Cervical" ~ "Ectocervix",
            Sample == "15cm rectal" ~ "Rectum",
            Sample == "Duodenal" ~ "Duodenum",
            Sample == "Vaginal" ~ "Vagina",
            Sample == "Rectal" ~ "Rectum",
            TRUE ~ Sample
        ))
    
    # require valid names
    enforce_valid_names(d$Sample, c("Vagina", "Rectum", "Whole blood", "PBMC", "Duodenum", "Ectocervix", "Endocervix"), "Invalid sample names")
    
    d
}

# convert visit names to canonical form for consistency
canonicalize_visit_names <- function(d) {
    d <- d %>% 
        mutate(Visit = case_when(
            Visit %in% c("post.prep", "Baseline", "Enrollment") ~ "Off treatment", 
            Visit %in% c("prep", "60 days", "Post-treatment", "Visit 2") ~ "On treatment", 
            TRUE ~ Visit
        ), 
        Visit = str_replace_all(Visit, "-", " "))
    
    # require valid names
    enforce_valid_names(d$Visit, c("Off treatment", "On treatment"), "Invalid visit names")
    
    d
}

# convert treatment names to canonical form for consistency
canonicalize_treatment_names <- function(d) {
    d <- d %>% 
        # Treatment is listed as "baseline" for MTN-017 and ACTU-3500 for the
        # baseline visits, but we want it to say the drug name that they will receive
        # which for both studies is truvada
        mutate(Treatment = case_when(
            Treatment %in% c("FTC/TDF", "Baseline", "Truvada") ~ "TDF/FTC",
            Treatment %in% c("TDF", "Tenofovir") ~ "TDF",
            TRUE ~ Treatment
        ))
    
    # require valid names
    enforce_valid_names(d$Treatment, c("TDF/FTC", "TDF"), "Invalid treatment names")
    
    d
}

# update gene names
# requires d to have a column containing entrez IDs
# adds a column named "TargetId" containing the corresponding gene symbols
canonicalize_gene_symbols <- function(d, entrez_id_column = "EntrezId", 
    current_symbols_column = "TargetId") {
    
    # it breaks below if the input has a column called "SYMBOL"
    stopifnot(all(colnames(d) != "SYMBOL"))
    
    # replace Illumina-supplied gene symbols with up-to-date gene symbols
    lookup <- suppressMessages(AnnotationDbi::select(org.Hs.eg.db::org.Hs.eg.db,
        keys = as.character(unique(d[[entrez_id_column]])),
        columns = c("ENTREZID", "SYMBOL"), keytype = "ENTREZID"))
    
    combined <- merge(d, lookup, by.x = entrez_id_column, by.y = "ENTREZID", all.x = TRUE) 
    
    # some probes don't have an EntrezId associated because they measure 
    # something other than a gene, such as an miRNA or an RNA of another type
    # in these cases, they do have a symbol, so get those symbols to use
    # in cases where there is no entrez ID and so AnnotationDbi::select returns
    # NA for symbol
    gene_names <- ifelse(is.na(combined[[entrez_id_column]]), combined[[current_symbols_column]], combined$SYMBOL)
    
    combined %>%
        mutate(TargetId = gene_names) %>%
        dplyr::select(-SYMBOL)
}

DIGITS_TO_SHOW <- 2

make_ci <- function(low, high) {
    paste("[", round(low, DIGITS_TO_SHOW), ", ", 
        round(high, DIGITS_TO_SHOW), "]", sep = "")
}

format_p <- function(p, digits_to_show = DIGITS_TO_SHOW) {
    ifelse(p < 0.01, 
        format(p, scientific = TRUE, digits = DIGITS_TO_SHOW + 1), 
        as.character(round(p, DIGITS_TO_SHOW))) 
}

adjust_and_ci <- function(d) {
    d %>% 
        arrange(p.value) %>% 
        # adjust for multiple comparisons
        mutate(adjusted = p.adjust(p.value, method = "bonferroni")) %>% 
        # difference = treated minus non-treated, so inverse of current estimate
        mutate(difference = estimate * -1, 
            ci = make_ci(-1 * conf.high, -1 * conf.low), 
            p.value = format_p(p.value), 
            adjusted = format_p(adjusted))
}

# read_csv gives a lot of messages/warnings if you don't supply col_types, which is 
# not always convenient, so this is a quiet version
# also prevents progress bar
quiet_read_csv <- function(...) {
    suppressMessages(suppressWarnings(read_csv(..., progress = FALSE)))
}
```
