## Supplementary material for "Treatment with commonly used antiretroviral drugs induces a type I/III interferon signature in the gut in the absence of HIV infection": Code to reproduce analyses and figures: README.html


### README

This document describes how to reproduce the data analysis and figures.

#### Obtaining the microarray data

There are four microarray datasets to download. All are available on GEO. Visit the four links below and from each one, download two Supplementary Files: `GSExxxxx_xxx_BeadStudio_output.txt.gz` and `GSExxxxx_xxx_sample_metadata.xlsx`. Place all eight files into `data/raw/`.

https://www.ncbi.nlm.nih.gov/geo/query/acc.cgi?acc=GSE139611  
https://www.ncbi.nlm.nih.gov/geo/query/acc.cgi?acc=GSE138723  
https://www.ncbi.nlm.nih.gov/geo/query/acc.cgi?acc=GSE139655  
https://www.ncbi.nlm.nih.gov/geo/query/acc.cgi?acc=GSE139411

#### Obtaining the other data

There are several other data types, which are all available in a figshare collection https://doi.org/10.6084/m9.figshare.c.4704827

Download the following files:

- `raw_ddPCR_data.zip`
- `ISG15_microscopy.xlsx`
- `microscopy_unblinded_log.csv`
- `Proteomics_MTN017.xlsx`
- `RNAseq_MTN017.xlsx`

Place all of these files into `data/raw`. Unzip `raw_ddPCR_data.zip` in that folder.

#### Running the analysis

Open `R/source_all.R` and install all of the packages listed there.

Open `make_paper.R`. Assuming that you have placed all of the data files listed above into `data/raw`, the file paths in this file will be correct. If you have placed them elsewhere, adjust the file paths. This assumes that the working directory is set to the top level of this folder. To reproduce the analysis, source `make_paper.R`. This will run code to:

- Load and clean all of the data files
- Perform statistical analysis
- Generate all figures
- Generate all statistics reported in the manuscript
- Generate the supplementary files

It will take a while to run.

Questions?

#### Session info

The analysis for the paper was run using the following versions of R and the packages:

```
## - Session info -----------------------------------------------------------------------------------
##  setting  value                       
##  version  R version 3.5.2 (2018-12-20)
##  os       Windows 10 x64              
##  system   x86_64, mingw32             
##  ui       RTerm                       
##  language (EN)                        
##  collate  English_United States.1252  
##  ctype    English_United States.1252  
##  tz       America/Los_Angeles         
##  date     2020-02-14                  
## 
## - Packages ---------------------------------------------------------------------------------------
##  package              * version   date       lib source                              
##  affy                   1.60.0    2018-10-30 [1] Bioconductor                        
##  affyio                 1.52.0    2018-10-30 [1] Bioconductor                        
##  annotate               1.60.0    2018-10-30 [1] Bioconductor                        
##  AnnotationDbi          1.44.0    2018-10-30 [1] Bioconductor                        
##  askpass                1.1       2019-01-13 [1] CRAN (R 3.5.2)                      
##  assertthat             0.2.1     2019-03-21 [1] CRAN (R 3.5.3)                      
##  backports              1.1.4     2019-04-10 [1] CRAN (R 3.5.3)                      
##  base64                 2.0       2016-05-10 [1] CRAN (R 3.5.2)                      
##  beanplot               1.2       2014-09-19 [1] CRAN (R 3.5.2)                      
##  bibtex                 0.4.2     2017-06-30 [1] CRAN (R 3.5.2)                      
##  Biobase              * 2.42.0    2018-10-30 [1] Bioconductor                        
##  BiocGenerics         * 0.28.0    2018-10-30 [1] Bioconductor                        
##  BiocManager            1.30.4    2018-11-13 [1] CRAN (R 3.5.2)                      
##  BiocParallel           1.16.6    2019-02-10 [1] Bioconductor                        
##  biomaRt                2.38.0    2018-10-30 [1] Bioconductor                        
##  Biostrings             2.50.2    2019-01-03 [1] Bioconductor                        
##  bit                    1.1-14    2018-05-29 [1] CRAN (R 3.5.2)                      
##  bit64                  0.9-7     2017-05-08 [1] CRAN (R 3.5.2)                      
##  bitops                 1.0-6     2013-08-17 [1] CRAN (R 3.5.2)                      
##  blob                   1.1.1     2018-03-25 [1] CRAN (R 3.5.2)                      
##  broom                * 0.5.2     2019-04-07 [1] CRAN (R 3.5.3)                      
##  bumphunter             1.24.5    2018-12-01 [1] Bioconductor                        
##  cellranger             1.1.0     2016-07-27 [1] CRAN (R 3.5.2)                      
##  cli                    1.1.0     2019-03-19 [1] CRAN (R 3.5.3)                      
##  codetools              0.2-15    2016-10-05 [2] CRAN (R 3.5.2)                      
##  colorspace             1.4-1     2019-03-18 [1] CRAN (R 3.5.3)                      
##  conflicted           * 1.0.1     2018-10-02 [1] CRAN (R 3.5.3)                      
##  crayon                 1.3.4     2017-09-16 [1] CRAN (R 3.5.2)                      
##  data.table             1.12.0    2019-01-13 [1] CRAN (R 3.5.2)                      
##  DBI                    1.0.0     2018-05-02 [1] CRAN (R 3.5.2)                      
##  DelayedArray           0.8.0     2018-10-30 [1] Bioconductor                        
##  DelayedMatrixStats     1.4.0     2018-10-30 [1] Bioconductor                        
##  digest                 0.6.21    2019-09-20 [1] CRAN (R 3.5.3)                      
##  doRNG                  1.7.1     2018-06-22 [1] CRAN (R 3.5.2)                      
##  dplyr                * 0.8.3     2019-07-04 [1] CRAN (R 3.5.3)                      
##  edgeR                * 3.24.3    2019-01-02 [1] Bioconductor                        
##  evaluate               0.14      2019-05-28 [1] CRAN (R 3.5.3)                      
##  forcats              * 0.4.0     2019-02-17 [1] CRAN (R 3.5.2)                      
##  foreach                1.4.4     2017-12-12 [1] CRAN (R 3.5.2)                      
##  genefilter             1.64.0    2018-10-30 [1] Bioconductor                        
##  generics               0.0.2     2018-11-29 [1] CRAN (R 3.5.2)                      
##  GenomeInfoDb           1.18.2    2019-02-12 [1] Bioconductor                        
##  GenomeInfoDbData       1.2.0     2019-02-20 [1] Bioconductor                        
##  GenomicAlignments      1.18.1    2019-01-04 [1] Bioconductor                        
##  GenomicFeatures        1.34.3    2019-01-28 [1] Bioconductor                        
##  GenomicRanges          1.34.0    2018-10-30 [1] Bioconductor                        
##  GEOquery               2.50.5    2018-12-22 [1] Bioconductor                        
##  ggplot2              * 3.2.1     2019-08-10 [1] CRAN (R 3.5.3)                      
##  ggrepel              * 0.8.1     2019-05-07 [1] CRAN (R 3.5.3)                      
##  glue                   1.3.1     2019-03-12 [1] CRAN (R 3.5.3)                      
##  gtable                 0.3.0     2019-03-25 [1] CRAN (R 3.5.3)                      
##  haven                  2.1.1     2019-07-04 [1] CRAN (R 3.5.3)                      
##  HDF5Array              1.10.1    2018-12-05 [1] Bioconductor                        
##  here                 * 0.1       2017-05-28 [1] CRAN (R 3.5.2)                      
##  hms                    0.5.1     2019-08-23 [1] CRAN (R 3.5.3)                      
##  htmltools              0.3.6     2017-04-28 [1] CRAN (R 3.5.2)                      
##  httr                   1.4.1     2019-08-05 [1] CRAN (R 3.5.3)                      
##  illuminaio             0.24.0    2018-10-30 [1] Bioconductor                        
##  IRanges                2.16.0    2018-10-30 [1] Bioconductor                        
##  iterators              1.0.10    2018-07-13 [1] CRAN (R 3.5.2)                      
##  jsonlite               1.6       2018-12-07 [1] CRAN (R 3.5.2)                      
##  KernSmooth             2.23-15   2015-06-29 [2] CRAN (R 3.5.2)                      
##  knitr                  1.25      2019-09-18 [1] CRAN (R 3.5.3)                      
##  lattice                0.20-38   2018-11-04 [2] CRAN (R 3.5.2)                      
##  lazyeval               0.2.2     2019-03-15 [1] CRAN (R 3.5.3)                      
##  lifecycle              0.1.0     2019-08-01 [1] CRAN (R 3.5.3)                      
##  limma                * 3.38.3    2018-12-02 [1] Bioconductor                        
##  locfit                 1.5-9.1   2013-04-20 [1] CRAN (R 3.5.2)                      
##  lubridate              1.7.4     2018-04-11 [1] CRAN (R 3.5.2)                      
##  lumi                 * 2.34.0    2018-10-30 [1] Bioconductor                        
##  magrittr               1.5       2014-11-22 [1] CRAN (R 3.5.2)                      
##  MASS                   7.3-51.1  2018-11-01 [2] CRAN (R 3.5.2)                      
##  Matrix                 1.2-15    2018-11-01 [2] CRAN (R 3.5.2)                      
##  matrixStats            0.54.0    2018-07-23 [1] CRAN (R 3.5.2)                      
##  mclust                 5.4.2     2018-11-17 [1] CRAN (R 3.5.2)                      
##  memoise                1.1.0     2017-04-21 [1] CRAN (R 3.5.2)                      
##  methylumi              2.28.0    2018-10-30 [1] Bioconductor                        
##  mgcv                   1.8-26    2018-11-21 [2] CRAN (R 3.5.2)                      
##  minfi                  1.28.3    2019-01-05 [1] Bioconductor                        
##  modelr                 0.1.5     2019-08-08 [1] CRAN (R 3.5.3)                      
##  msigdbr              * 6.2.1     2018-10-09 [1] CRAN (R 3.5.3)                      
##  multtest               2.38.0    2018-10-30 [1] Bioconductor                        
##  munsell                0.5.0     2018-06-12 [1] CRAN (R 3.5.2)                      
##  nleqslv                3.3.2     2018-05-17 [1] CRAN (R 3.5.2)                      
##  nlme                   3.1-137   2018-04-07 [2] CRAN (R 3.5.2)                      
##  nor1mix                1.2-3     2017-08-30 [1] CRAN (R 3.5.2)                      
##  openssl                1.2.1     2019-01-17 [1] CRAN (R 3.5.2)                      
##  pander               * 0.6.3     2018-11-06 [1] CRAN (R 3.5.2)                      
##  patchwork            * 0.0.1     2019-03-12 [1] Github (thomasp85/patchwork@fd7958b)
##  pillar                 1.4.2     2019-06-29 [1] CRAN (R 3.5.3)                      
##  pkgconfig              2.0.3     2019-09-22 [1] CRAN (R 3.5.3)                      
##  pkgmaker               0.27      2018-05-25 [1] CRAN (R 3.5.2)                      
##  plater               * 1.0.1     2017-06-26 [1] CRAN (R 3.5.3)                      
##  plyr                   1.8.4     2016-06-08 [1] CRAN (R 3.5.2)                      
##  preprocessCore         1.44.0    2018-10-30 [1] Bioconductor                        
##  prettyunits            1.0.2     2015-07-13 [1] CRAN (R 3.5.2)                      
##  progress               1.2.0     2018-06-14 [1] CRAN (R 3.5.2)                      
##  purrr                * 0.3.2     2019-03-15 [1] CRAN (R 3.5.3)                      
##  quadprog               1.5-5     2013-04-17 [1] CRAN (R 3.5.2)                      
##  R6                     2.4.0     2019-02-14 [1] CRAN (R 3.5.2)                      
##  RColorBrewer           1.1-2     2014-12-07 [1] CRAN (R 3.5.2)                      
##  Rcpp                   1.0.2     2019-07-25 [1] CRAN (R 3.5.3)                      
##  RCurl                  1.95-4.11 2018-07-15 [1] CRAN (R 3.5.2)                      
##  readr                * 1.3.1     2018-12-21 [1] CRAN (R 3.5.2)                      
##  readxl               * 1.3.1     2019-03-13 [1] CRAN (R 3.5.3)                      
##  registry               0.5       2017-12-03 [1] CRAN (R 3.5.2)                      
##  reshape                0.8.8     2018-10-23 [1] CRAN (R 3.5.2)                      
##  rhdf5                  2.26.2    2019-01-02 [1] Bioconductor                        
##  Rhdf5lib               1.4.2     2018-12-03 [1] Bioconductor                        
##  rlang                  0.4.0     2019-06-25 [1] CRAN (R 3.5.3)                      
##  rmarkdown              1.11      2018-12-08 [1] CRAN (R 3.5.2)                      
##  rngtools               1.3.1     2018-05-15 [1] CRAN (R 3.5.2)                      
##  rprojroot              1.3-2     2018-01-03 [1] CRAN (R 3.5.2)                      
##  Rsamtools              1.34.1    2019-01-31 [1] Bioconductor                        
##  RSQLite                2.1.1     2018-05-06 [1] CRAN (R 3.5.2)                      
##  rstudioapi             0.10      2019-03-19 [1] CRAN (R 3.5.3)                      
##  rtracklayer            1.42.1    2018-11-21 [1] Bioconductor                        
##  rvest                  0.3.4     2019-05-15 [1] CRAN (R 3.5.3)                      
##  S4Vectors              0.20.1    2018-11-09 [1] Bioconductor                        
##  scales                 1.0.0     2018-08-09 [1] CRAN (R 3.5.2)                      
##  sessioninfo            1.1.1     2018-11-05 [1] CRAN (R 3.5.2)                      
##  siggenes               1.56.0    2018-10-30 [1] Bioconductor                        
##  stringi                1.4.3     2019-03-12 [1] CRAN (R 3.5.3)                      
##  stringr              * 1.4.0     2019-02-10 [1] CRAN (R 3.5.2)                      
##  SummarizedExperiment   1.12.0    2018-10-30 [1] Bioconductor                        
##  survival               2.43-3    2018-11-26 [2] CRAN (R 3.5.2)                      
##  tibble               * 2.1.3     2019-06-06 [1] CRAN (R 3.5.3)                      
##  tidyr                * 1.0.0     2019-09-11 [1] CRAN (R 3.5.3)                      
##  tidyselect             0.2.5     2018-10-11 [1] CRAN (R 3.5.2)                      
##  tidyverse            * 1.2.1     2017-11-14 [1] CRAN (R 3.5.2)                      
##  vctrs                  0.2.0     2019-07-05 [1] CRAN (R 3.5.3)                      
##  withr                  2.1.2     2018-03-15 [1] CRAN (R 3.5.2)                      
##  writexl              * 1.1       2018-12-02 [1] CRAN (R 3.5.3)                      
##  xfun                   0.10      2019-10-01 [1] CRAN (R 3.5.3)                      
##  XML                    3.98-1.17 2019-02-08 [1] CRAN (R 3.5.2)                      
##  xml2                   1.2.2     2019-08-09 [1] CRAN (R 3.5.3)                      
##  xtable                 1.8-3     2018-08-29 [1] CRAN (R 3.5.2)                      
##  XVector                0.22.0    2018-10-30 [1] Bioconductor                        
##  yaml                   2.2.0     2018-07-25 [1] CRAN (R 3.5.2)                      
##  zeallot                0.1.0     2018-01-28 [1] CRAN (R 3.5.3)                      
##  zlibbioc               1.28.0    2018-10-30 [1] Bioconductor                        
## 
## [1] C:/Users/Sean Hughes/Documents/R/win-library/3.5
## [2] C:/Program Files/R/R-3.5.2/library
```
